# A Customizable Low-Cost System for Massively Parallel Zebrafish Behavior Phenotyping

**DOI:** 10.1101/2020.09.08.288621

**Authors:** William Joo, Michael D. Vivian, Brett J. Graham, Edward R. Soucy, Summer B. Thyme

## Abstract

High-throughput behavioral phenotyping is critical to genetic or chemical screening approaches. Zebrafish larvae are amenable to high-throughput behavioral screening because of their rapid development, small size, and conserved vertebrate brain architecture. Existing commercial behavior phenotyping systems are expensive and not easily modified for new assays. Here, we describe a modular, highly adaptable, and low-cost behavior system. Along with detailed assembly and operation instructions, we provide data acquisition software and a robust, parallel analysis pipeline. We validate our approach by analyzing stimulus response profiles in larval zebrafish, confirming prepulse inhibition phenotypes of two previously isolated mutants, and highlighting best practices for growing larvae prior to behavioral testing. Our new design thus allows rapid construction and streamlined operation of many large-scale behavioral setups with minimal resources and fabrication expertise, with broad applications to other aquatic organisms.

## 1 Introduction

High-throughput behavior tracking offers great potential for large-scale mutant phenotyping (Thyme et al., 2019) and drug screening (MacRae and Peterson, 2015). Indeed, drug screens have revealed conserved signaling pathways that regulate complex behaviors in both zebrafish and mammals (Kokel et al., 2010; Rihel et al., 2010; Leung and Mourrain, 2016). Furthermore, larval zebrafish maintained in 96-well plate format execute diverse behaviors including prepulse inhibition, sleep, seizures, prey capture, and responses to visual, acoustic, or thermal stimuli (Burgess and Granato, 2007; Chiu et al., 2016; Randlett et al., 2019; Griffin et al., 2020). Many researchers use commercial systems to test these behaviors, but such solutions are limited in their adaptability and prohibitively costly when many parallel systems are required.

For example, two of the most commonly used commercial systems are the DanioVision from Noldus and the ZebraBox from ViewPoint. Standard versions of these systems possess limited applications as they provide only baseline movement tracking and LED light control. While add-ons such as high-speed cameras and acoustic stimulation are available, they greatly increase system cost. Furthermore, users are often limited to commercially provided analysis code and data processing formats.

To bypass these challenges, we present building plans for a modular behavior setup (Figure 1A), together with software for data acquisition and analysis. Our new design significantly extends systems previously validated in a large-scale mutant screen (Thyme et al., 2019), with more precise control over a broader range of assays and greater ease of construction. This system includes most of the assays of commercially available solutions and easily accommodates additional modules. Our highly efficient analysis software utilizes a high-performance computing cluster for parallel processing of multi-day datasets with hundreds of user-defined events. Additionally, we outline best experimental practices for yielding consistent and reliable behavior data. This fully customizable and modular setup can be easily adapted as new behavior assays are published, significantly lowering barriers to large-scale phenotyping approaches.

**Figure 1.**
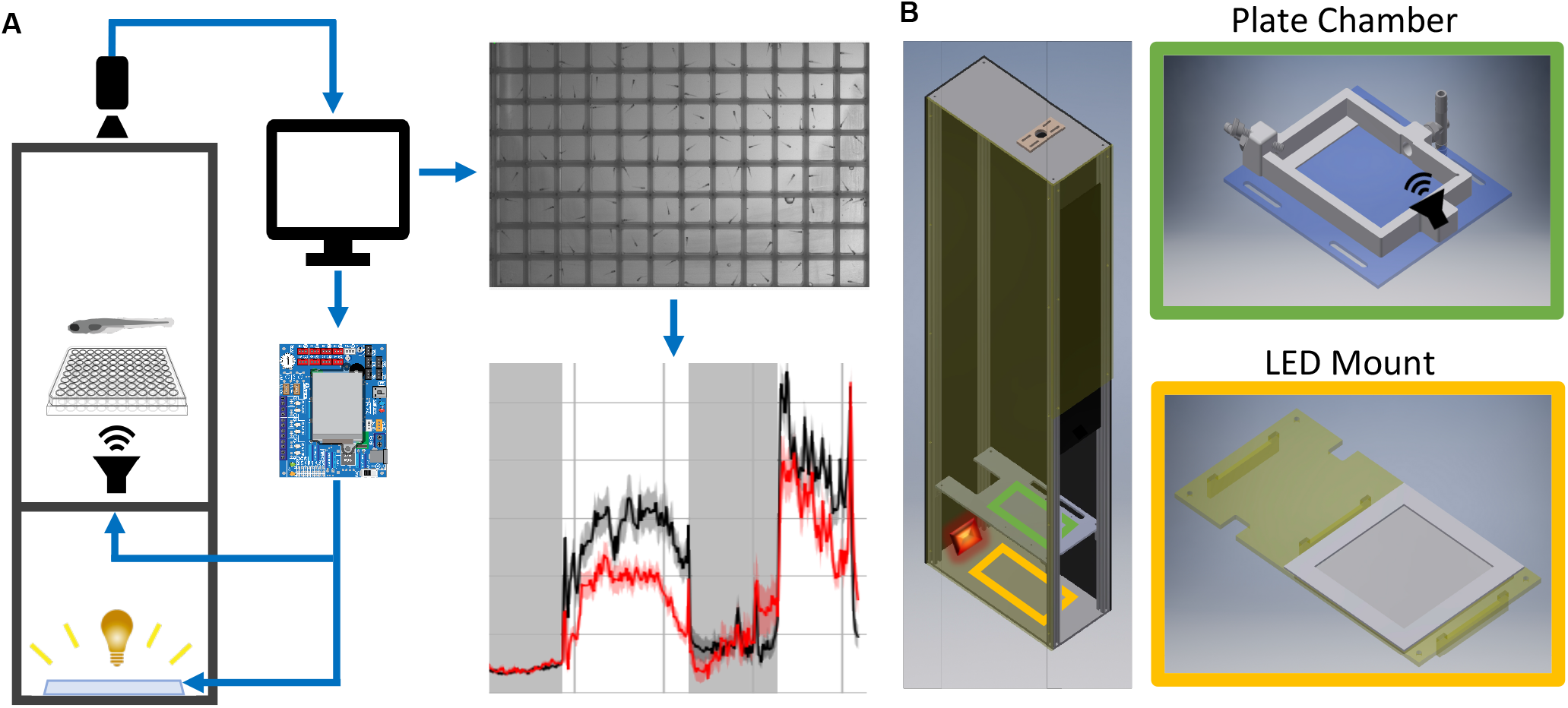
Behavior Box Overview. **(A)** Schematic of the behavior box setup. A high-speed camera is mounted on top of the box and focused on the fish plate. A microcontroller circuit is connected to a white LED panel at the bottom of a box and a surface transducer attached to the plate holder, which deliver visual and acoustic stimuli, respectively. The microcontroller and camera are connected to a desktop computer, which uses custom LabVIEW software for data acquisition and experiment control. **(B)** Left: Setup enclosure is affixed to an aluminum frame with clear acrylic shelving for the plate holder and LED panel. Right: The 3D printed fish chamber includes input/output nozzles for water circulation and a screw stud for the surface transducer. See supplemental files for parts list and assembly instructions (Supplementary Figures 1-8).

## 2 Materials and Methods

### 2.1 Materials

All components and costs are described in the Bill of Materials, and all schematics are included in FabricationFiles.zip. While access to a laser cutter and 3D printer substantially decreases cost and time of construction, online manufacturing websites can easily produce equivalent parts (see Supplementary Material).

### 2.2 Box Assembly

Supplementary Material describes all assembly steps. The setup housing consists of a light-insulated enclosure, a camera to track fish motion, and a computer/electronics setup to deliver stimuli (Figure 1A). The enclosure was laser-cut from high-density polyethylene (HDPE) and fastened with 80/20 rails (Figure 1B).

The enclosure contains a white LED panel to deliver ambient light or stimuli, an infrared (IR) light to visualize animals, and a 3D-printed fish plate holder with a mounted acoustic transducer (Figure 1B). The white LED panel is mounted on an acrylic shelf and illuminates fish from below, while the IR light rests behind and reflects off the white light panel. Fish were detected with a Grasshopper3 camera (FLIR Systems) and a 50 mm fixed focal length lens with an IR filter.

### 2.3 Data Acquisition

See Supplementary Material for detailed operation instructions.

#### Computer hardware

The setup was operated using a standard desktop computer and custom LabVIEW software (Supplementary Software). Minimum hardware requirements for the most computationally demanding assay (acoustic habituation; one-second movies at 285 frames-per-second [fps] acquired every two seconds) were 16.0 GB RAM, an Intel Core i7-9700 processor, Windows 10, and a 1 TB Solid-State Drive.

#### Stimulus Delivery and Data Collection

Acoustic and visual stimuli were controlled by a circuit board that communicates between LabVIEW software and system devices (Figure 2A). A Teensy 3.6 microcontroller and custom Arduino script relays stimulus command strings to LabVIEW (Figure 2B). Each “command string” specifies stimulus parameters such as amplitude (a), frequency (f), duration (d), and delay times (D) (full list in Supplementary Material). The microcontroller then sends voltage changes to the surface transducer or LED light panel to produce stimuli. The “command ID” (Figure 2B) specifies the LabVIEW event type, such as high-speed movie acquisition during stimulus presentation.

**Figure 2.**
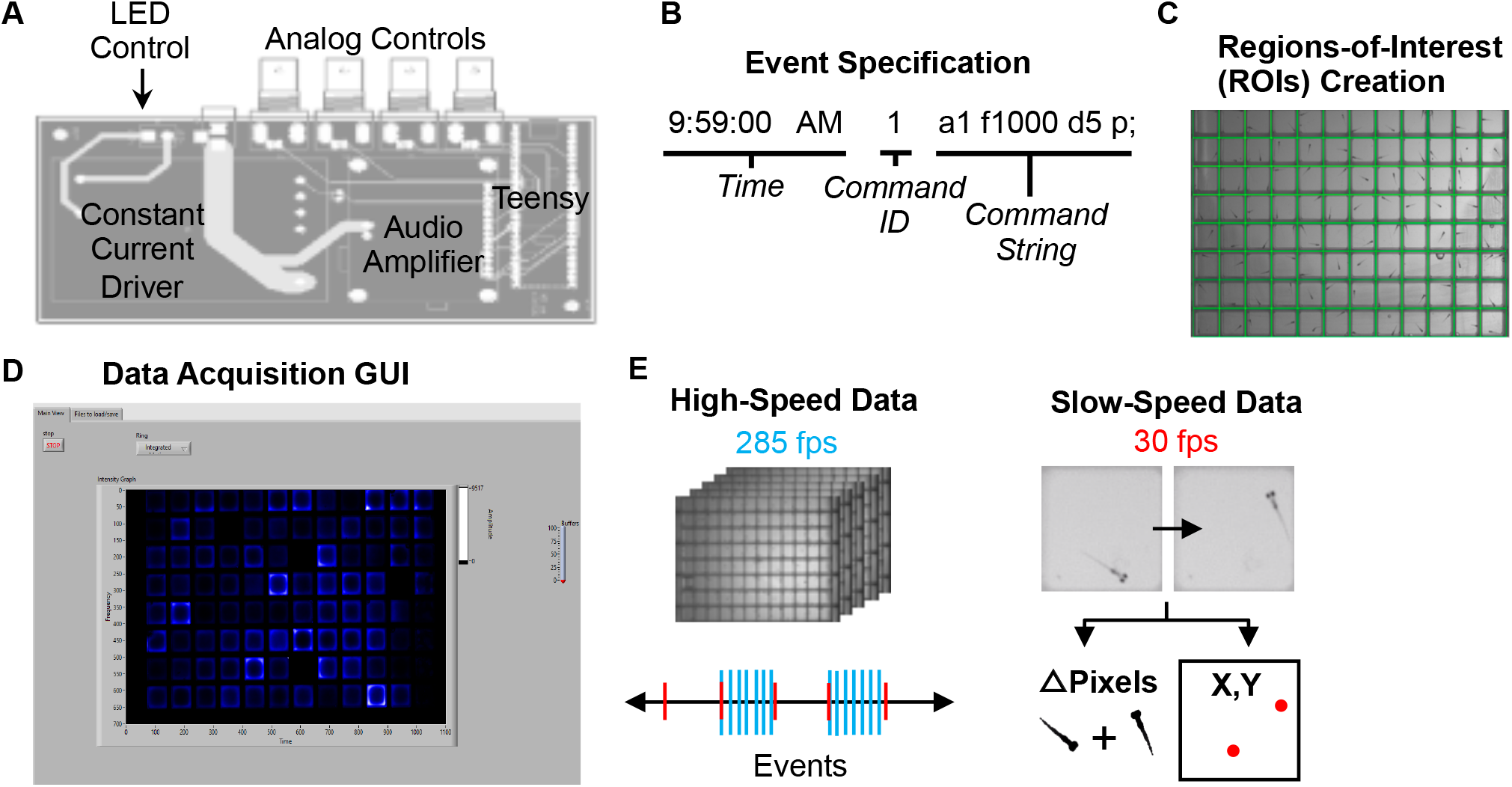
Data Acquisition Control. **(A)** Printed circuit board for electronics control. The LED light panel and surface transducer are manipulated by a Teensy 3.6 microcontroller with a constant current LED driver and an audio amplifier. A custom Arduino script with command options is uploaded to the microcontroller. The board also includes four BNC connectors wired to GPIO pins on the Teensy that support digital input/output, analog input, and other functionality configurable from software. For instance, a photodiode can be connected to calibrate the light panel. **(B)** Example command string to specify an event such as lights-off or high-speed movie acquisition during stimulus. See Supplementary Software for an example events file. **(C)** Users define regions of interest corresponding to each well using a LabVIEW graphical interface. Event parameters and ROIs are then transferred to main experiment software. **(D)** The LabVIEW data acquisition interface can display fish movements in real-time. **(E)** High-speed data is captured as 1 second 285 fps AVI movies as specified in the events file. Slow-speed data is collected at 30 fps to produce motion (delta pixels) and centroid (coordinates of fish centroid) files for the entirety of the experiment. Slow-speed data is continuously acquired regardless of high-speed events. See Supplementary Figures 9-12 for information regarding data acquisition pipeline.

To run an experiment, users 1) construct an **events file** with desired command strings, 2) designate regions of interest (ROIs) using a separate LabVIEW script (“Generate ROIs.vi”) (Figure 2C), which generates an **ROI binary data file** (rois) and **ROI string text file** (rois_string). ROIs can match many different multi-well plate formats. 3) Select the events file, the ROI binary data file, and a png image of the plate using the LabVIEW graphical user interface (GUI) (Figure 2D). Users also define data output names and folders. See Supplementary Material for detailed setup instructions. The ROI string text file is used in later stages of the analysis.

30 fps data is collected for the duration of the experiment in two formats: the change in pixels between each frame within each ROI, and the coordinates of the centroid of each fish in each ROI (**Slow-speed data**, Figure 2E). User-defined LabVIEW events trigger acquisition of one-second movies at 285 fps (**High-speed data**). LabVIEW can also trigger acquisition of 30 fps movies of desired length.

### 2.4 Data Analysis Software

Our analysis pipeline (Figure 3) is based in the Python programming language. All analyses were performed on a high-performance computing cluster due to vastly increased parallel processing capacity. LabVIEW generates slow-speed **motion** (delta pixels) and **centroid** (coordinates) data, while our Python scripts extract motion and centroid from high-speed data (Figure 3A). As in LabVIEW, a centroid for each fish is identified in each ROI to determine coordinates. Typical behavior runs often produce close to one thousand high-speed movies, making parallel processing critical to this tracking step.

**Figure 3.**
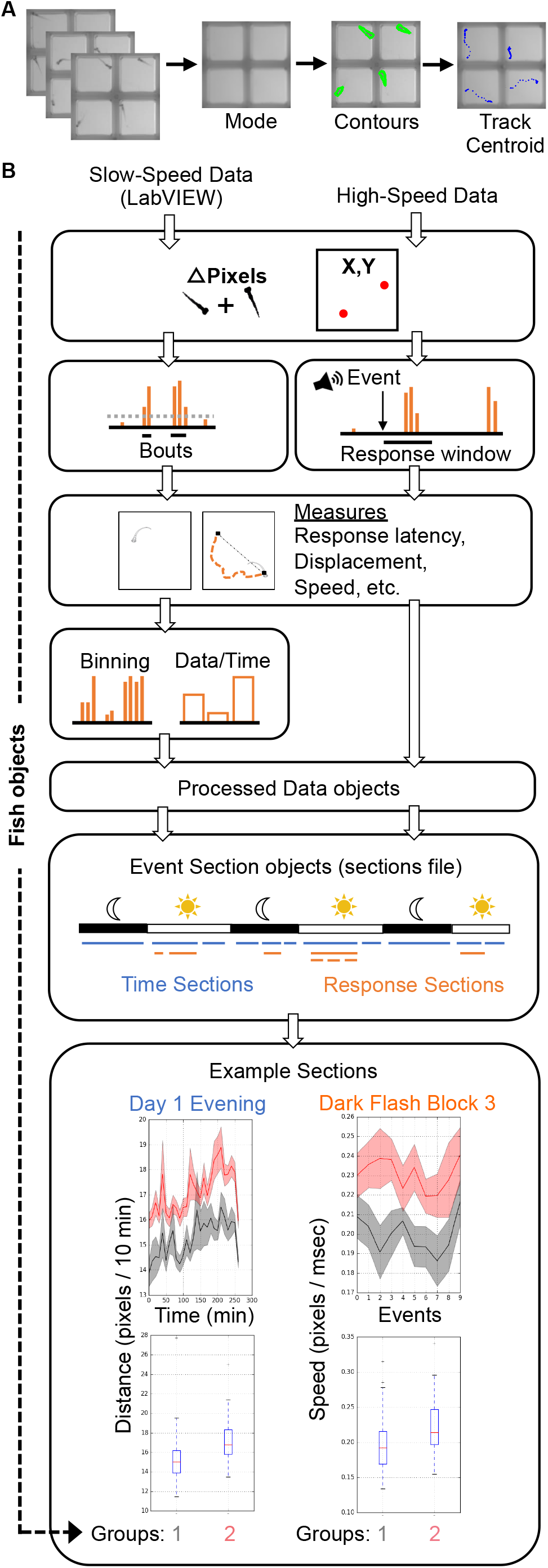
Data Analysis Pipeline. **(A)** The high-speed movies are analyzed by calculating and subtracting a mode image to define frame-by-frame fish contours in each well or ROI, and tracking the centroid of each fish to generate delta pixel and position data. **(B)** Overview of high- and slow-speed data processing and group comparisons. Fish objects contain all metrics for each fish as well as its genotype (gray = group 1, red = group 2). For slow-speed data, bouts are identified using the delta pixel and positional information. Thresholds are set depending on input data type. Stimulus responses are identified analogously, but are identified with high-speed movie frames. Movement and response features are then calculated, binned if slow-speed data, and plotted based on user defined event sections. For example, slow-speed data is processed in sections based on time, such as “Day 1 Evening” or “Day 2 Night”. High-speed data processing considers only the high-speed movie information in a given section, such as the 10 dark flashes in “Dark Flash Block 3”. Sections can be overlapping. Current outputs include both a ribbon plot and a box plot for each metric.

Input data from slow- and high-speed tracking is processed to generate numerous measurements and output graphs (Figure 3B) ranging from classic behaviors such as sleep bouts and waking activity (Chen et al., 2017) to recently published observations such as turn angle preference during dark flash response (Horstick et al., 2020). A **Fish object** is created for each animal and contains all associated slow- and high-speed data as well as genotype. Slow-speed data is converted into movement bouts calculated from both motion data and centroid data. Metrics such as frequency, velocity, and fraction of time in well center are calculated for each bout and binned based on time (such as average velocity / 10 minute). High-speed data is processed based on the type of event and the parameters of the event string. Identification of an event response depends on modality (visual, acoustic) and the time delays in the string. Metrics analogous to bout properties are then calculated for the response. High-speed and binned slow-speed data are returned to the Fish object as **ProcessedData objects**, which are then used to generate graphs according to user-defined **event sections**. The event sections are specified in the **sections file**, which segments the behavior run into time windows for different assays. For example, an acoustic habituation assay would be analyzed separately from the prepulse inhibition assay. Event sections may also correspond to different times of the run such as night or day, and need not include high-speed events. Sections without high-speed events are referred to as “time” sections. An example sections file is included in the Supplementary Software. Data and statistics are saved and a graph is generated for every combination of an EventSections object and ProcessedData object. A Kruskal-Wallis one-way ANOVA is calculated for every metric, and a linear mixed model (Thyme et al., 2019) is also calculated for baseline data with a time component. The code is also available on GitHub (https://github.com/sthyme/ZebrafishBehavior) and will be updated as improvements are made.

### 2.5 Assays

The most common multi-well larval zebrafish assays are based on acoustic and visual stimulation, utilizing the surface transducer and the LED light panel. These include responses to increased light or decreased light (dark flash), dark flash habituation, acoustic responses and thresholds, prepulse inhibition, and acoustic habituation. Our setup can test responses to a broad range of acoustic (sound delivered by surface transducer), visual (whole-field luminance changes such as dark or light flashes), and thermal stimuli (cooling or heating with a water circulation system; see Supplementary Material), and can be further modified for additional assays. To test arousal threshold (Figure 5), we delivered acoustic (20 msec, 625 Hz, square wave-form) or dark flash (1 sec) stimuli at 12 different intensities: acoustic = a0.0005, a0.001, a0.003, a0.0075, a0.01, a0.03, a0.06, a0.075, a0.1, a0.3, a0.5, a1, visual = b245, b240, b230, b220, b210, b200, b175, b150, b125, b100, b50, b0, with baseline light = b250. 30-50 total trials of each intensity were administered in randomized order at 2 minute intervals. Other experiments for mutant and wild-type animals correspond to the event strings in the supplemental events file. The design also includes a mini-projector underneath the fish plate, which can present user-defined movies such as moving gratings to induce the optomotor response (Figure 4) (Naumann et al., 2016). Movies are presented through LabVIEW via the VLC media player (LabVIEW utilizes movie filepaths instead of command strings). Supplementary Software includes example grating movies and Python script to generate gratings. Code to track multiple animals was completed with a custom (http://github.com/docviv/behavior-scripts) based on an algorithm adapted from (Bolton et al., 2019).

**Figure 4.**
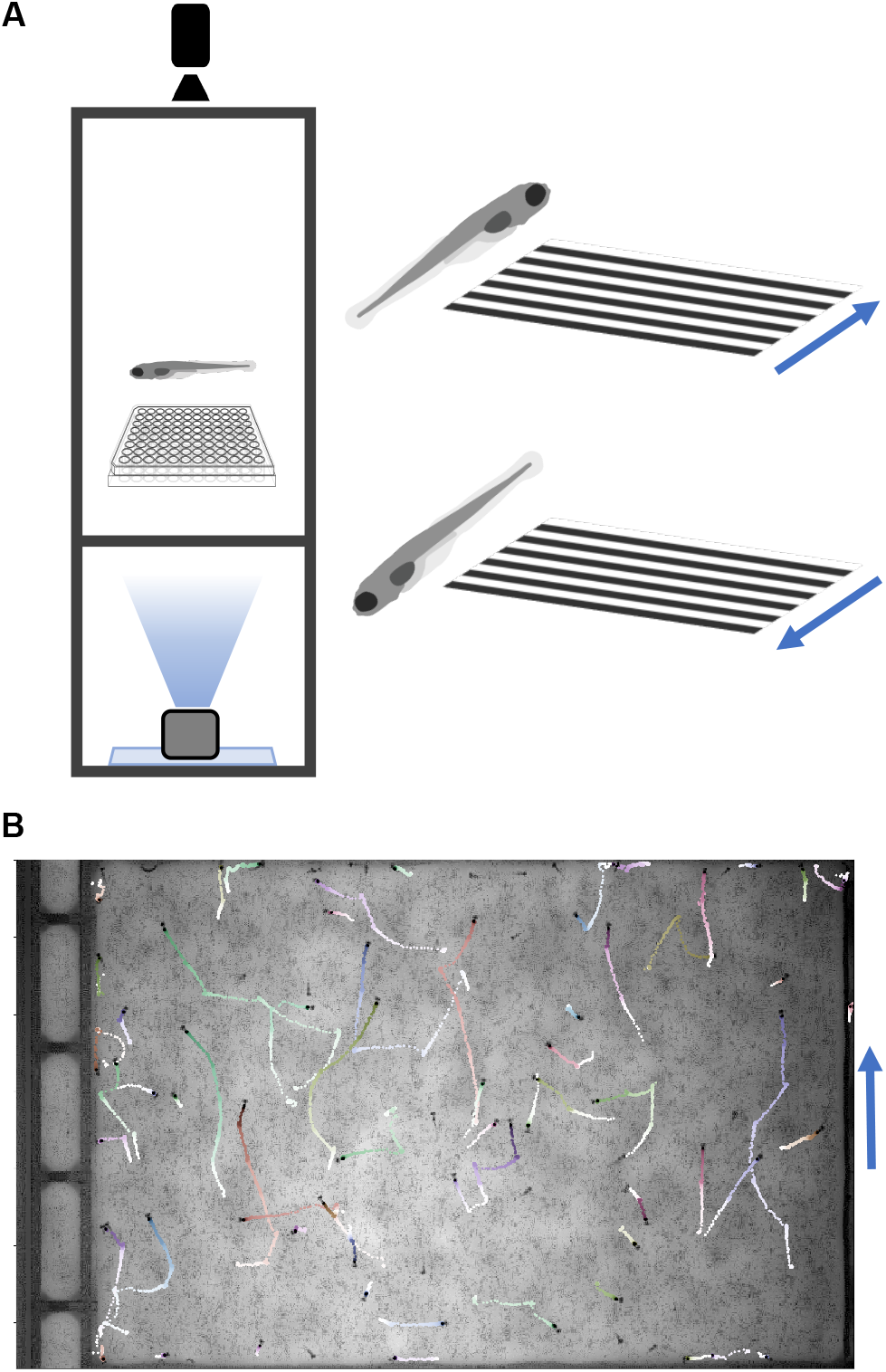
Optomotor Response Assay. **(A)** Diagram of the optomotor response assay. Larvae are tracked while a moving grating is projected from below. Blue arrows denote grating movement direction. **(B)** Fish trajectories (light to dark) during a 25-sec movie (30 fps). Multiple fish were placed in a plate without well dividers. Tracking code is available on GitHub (Methods).

**Figure 5.**
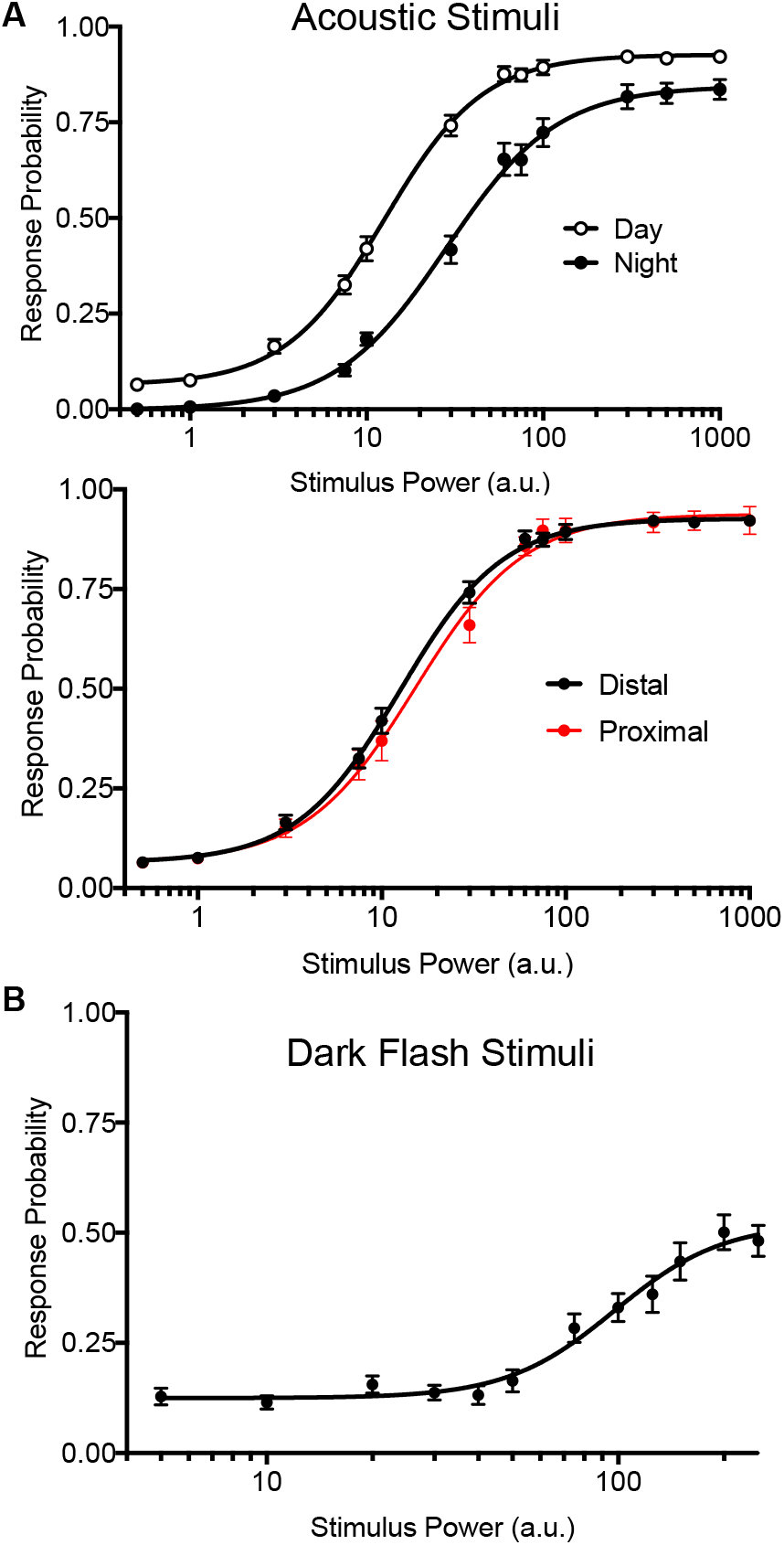
Precise stimulus control. **(A)** *Top*: Acoustic stimulus response curves for wild-type larvae during night and day. Night arousal threshold= 28.30±2.06, Day arousal threshold=12.57±1. N=90 larvae. *Bottom*: Daytime acoustic stimulus response curves comparing location relative to the transducer. Proximal group includes larvae in the three rows closest to the transducer (N=24). **(B)** Dark flash stimulus response curve for wild-type animals. Responses were filtered to include only O-bend startle responses.

### 2.6 Zebrafish Husbandry

All zebrafish were housed in the Zebrafish Research Facility of the University of Alabama at Birmingham and experiments were approved under protocol number IACUC-21744 (UAB Institutional Animal Care and Use Committee; Birmingham, Alabama). All crosses were derived from a single parental pair (mainly Ekkwill strain) to minimize genetic background differences. Arousal threshold assays were conducted in a mixed TL/AB background. Larvae were grown in 150×15 mm petri dishes with standard methylene blue water, at a density of less than 150 fish per plate. Animals were maintained at 28°C and a 14/10 light/dark cycle. Behavior experiments were conducted on the same light/dark cycle. Dead material and debris were removed twice before 4 dpf (afternoons of day 0 and day 2). All behavior assays were conducted on zebrafish larvae 4-7 days post-fertilization (dpf). Zebrafish of any age can be monitored in this setup with an appropriate holding chamber.

### 2.7 Zebrafish Sample Processing

Only healthy larvae with normal swim bladder morphology were included in experiments. Larvae were arrayed in 96-well plates (E&K Scientific Cat#2074, 0.7mL/square well volume) in standard methylene blue water. The plate was placed in an ice-water bath until movement abated and sealed with an air-permeable film (Thermo Fisher Scientific Cat#4311971) to eliminate progressive water evaporation during multi-day experiments (Supplementary Figure 13). Sealing is essential to long-term experiments but incompatible with drug delivery. Accordingly, previous drug screens for sleep modulators refilled evaporated water in the evening and morning (Rihel et al., 2010). Sealed plates were placed into the behavior box and secured tightly (screw in one corner) to prevent movement due to the surface transducer. Temperature inside the setup ranged from 29.5-30.5°C (measured with a wireless Temp Stick), while room temperature was maintained at 23°C. For mutant experiments, larvae were genotyped by 1) noting all dead or unhealthy animals, 2) cooling plate on ice until movement ceased, 3) removing water in wells, 4) immersing in sodium hydroxide and transferring to a PCR plate for DNA extraction and amplification.

## 3 Results

### 3.1 Precise Stimulus Control

Previous versions of our setup used two solenoid tappers and a custom white LED array to deliver acoustic and visual stimuli, respectively (Thyme et al., 2019). Stimulus intensity was inconsistent across setups due to variable construction. For example, solenoid tappers delivered limited and inconsistent tap strengths due to variable height alignment and spring properties, and suffered from artifacts such as inadvertent double or triple tapping (data not shown). The single mounted surface transducer now allows consistent and fine control over a broad range of stimulus durations, voltages, waveforms, and frequencies. Likewise, the new white LED panels deliver consistent luminance across a broad range across setups. We include a simple protocol to calibrate and standardize light levels using a photodiode (see Supplementary Material). To validate these modifications, we monitored larval zebrafish responses to acoustic and dark flash stimuli of variable intensities during day and night. By calculating “dose-response” curves for each type of stimulus, we determined arousal threshold, defined as stimulus strength generating half-maximal response probability (Figure 5). Larvae exhibited significantly higher arousal threshold during night relative to day (Figure 5A, top; night threshold= 28.30±2.06, day threshold=12.57±1). Acoustic stimulus response probabilities did not differ between fish positioned proximally or distally to the transducer, indicating consistent stimulus delivery across the 96-well plate (Figure 5A, bottom). While maximal dark flash responses matched previously reported levels (Figure 5B) (Woods et al., 2014), we observed improved maximal responses to acoustic stimuli relative to previous assays using solenoids (Lee et al., 2017; Singh et al., 2017). Our modifications thus accommodate previously challenging assays and offer improved standardization. Furthermore, our improved analysis code distinguishes clear escape responses (C-bends (Burgess and Granato, 2007) and O-bends (Randlett et al., 2019)) from smaller movements, for more nuanced response quantifications.

### 3.2 Mutant Prepulse Inhibition Phenotypes

We previously demonstrated (Thyme et al., 2019) that mutants for the schizophrenia risk genes *atxn7* and *akt3* (Schizophrenia Working Group of the Psychiatric Genomics Consortium, 2014; Bergeron et al., 2017) exhibit defects in prepulse inhibition (PPI), a sensory-motor gating phenomenon in which a weak prepulse stimulus suppresses an immediately following strong stimulus response. Because previous experiments relied on solenoid tappers, we tested whether the surface transducer recapitulates the PPI assay and phenotypes. Indeed, *atxn7* and *akt3* mutants both exhibited decreased PPI relative to sibling controls as previously observed (Figure 6A). Frequency of response to the strong stimulus in mutants and controls is substantially reduced when preceded by the weak prepulse (Figure 6B), indicating that the prepulse is effectively inhibiting responsiveness. Mutant PPI responses were increased in both frequency and aspects of the response motion (Figure 6C), whereas control larvae responses were not.

**Figure 6.**
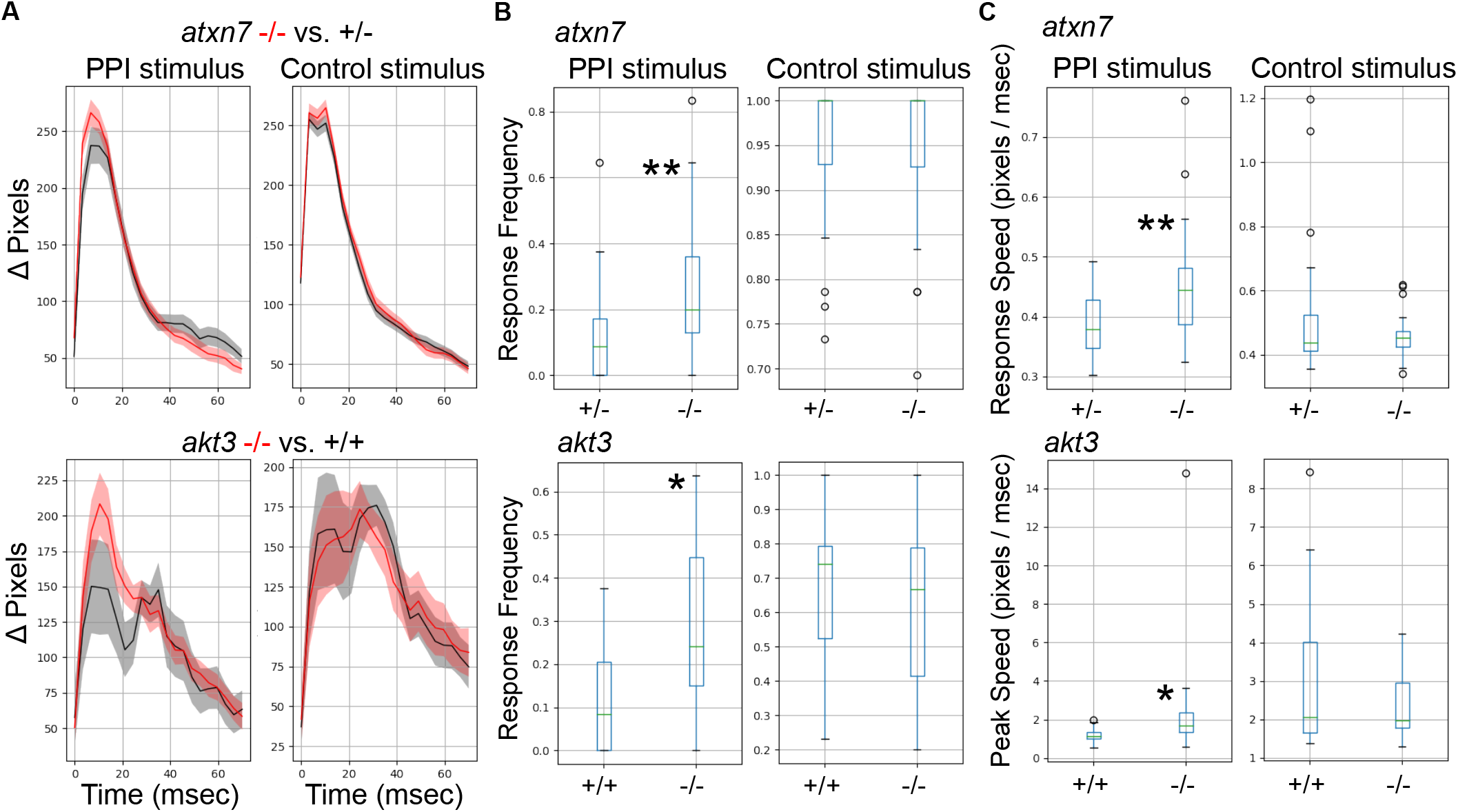
Analysis of Mutants with a Prepulse Inhibition Phenotype. **(A)** Responses to acoustic prepulse inhibition (PPI) strong stimulus and a control isolated strong stimulus not preceded by a prepulse, quantified as change in pixels during a 1-second high-speed movie. The weak prepulse (not shown) does not elicit a significant response and any prepulse-responding larvae are not considered in the calculations. Red, mutant; Gray, sibling control (all 5 dpf). Both mutants display increased response frequency to the PPI strong stimulus but similar response to the control isolated stimulus. **(B)** PPI and control response frequencies for each mutant. *atxn7* PPI response frequency: Kruskal-Wallis p-value = 0.0006. *atxn7* strong response frequency: Kruskal-Wallis p-value = 0.79. *akt3* PPI response frequency: Kruskal-Wallis p-value = 0.012. *akt3* strong response frequency: Kruskal-Wallis p-value = 0.64. **(C)** PPI and control response speed for each mutant. *atxn7* PPI response speed: Kruskal-Wallis p-value = 0.0092. *atxn7* strong response speed: Kruskal-Wallis p-value = 0.83. *akt3* PPI response peak speed: Kruskal-Wallis p-value = 0.016. *akt3* strong response peak speed: Kruskal-Wallis p-value = 0.48. *atxn7* N -/- = 27, +/- = 48. *akt3* N -/- = 16, +/+ = 16. Single asterisk marks p-value < 0.05, double marks p-value < 0.01.

### 3.3 Wild Type Comparisons

While commonly used wild-type zebrafish strains exhibit substantial genetic diversity (Guryev et al., 2006; Brown et al., 2012), few studies explicitly define optimal growth and husbandry conditions that minimize possibly resultant behavioral variability.

As a first step to defining important parameters, we assessed three different conditions on larval zebrafish behavior. 1) To compare separately reared larvae, we divided sibling larvae into two dishes at identical density (Figure 7A). 2) To assess effects of density, we compared sibling larvae reared in two dishes of high or low density. 3) To compare non-siblings, we raised different clutches at identical densities. For each experiment, we also compared within each experimental group as a control and interleaved animals from each condition in the 96-well plate to minimize possible positional effects (Figure 7B).

**Figure 7.**
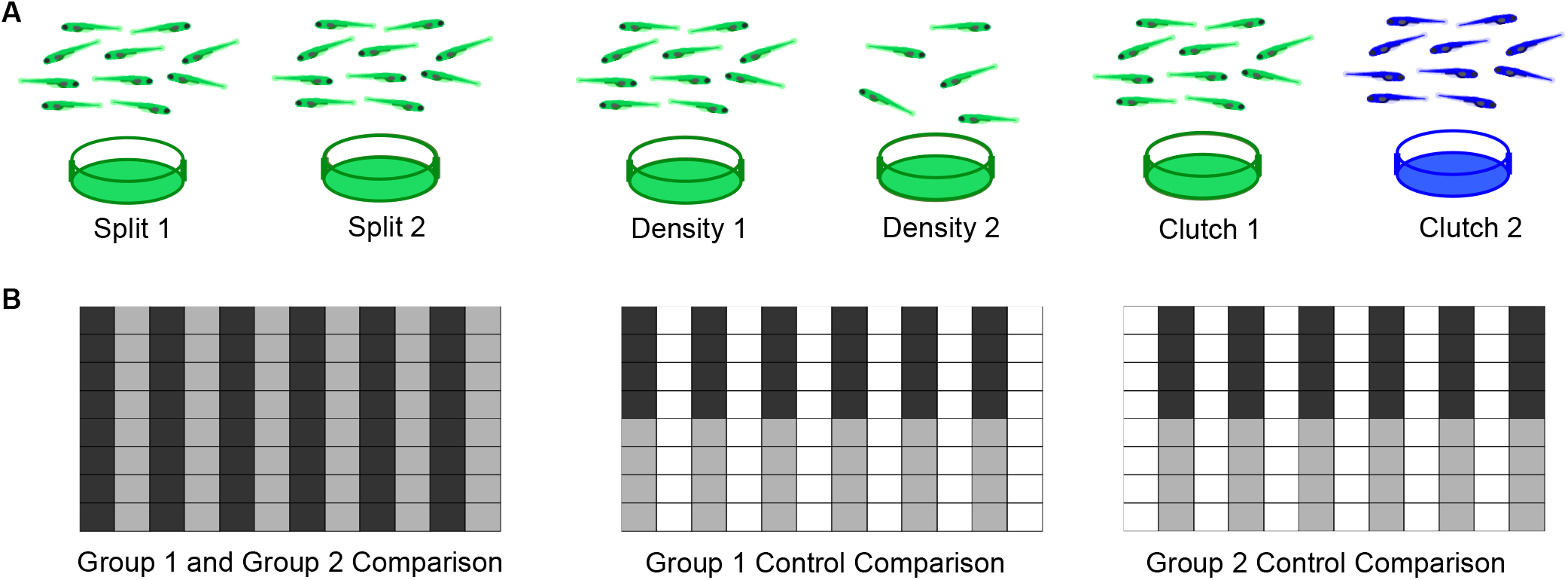
Experimental Design for Wild Type Comparisons. **(A)** *Left*: Comparison of a single clutch split between two petri dishes with a density of 140-150 fish per dish. *Middle*: Comparison of a single clutch split between two petri dishes of different densities: 140-150 fish per dish vs 60-70 fish. *Right*: Comparison of two different clutches with equal densities 140-150 fish per dish). **(B)** Format of the 96-well plate organization for each comparison, where gray and black indicate the two experimental groups loaded in alternating columns. *Left*: Comparison between the two experimental groups; *Middle and Right*: control comparisons within each experimental group.

To estimate behavioral differences, we calculated strictly standardized mean difference (SSMD) values across all behavioral parameters (SSMD of 0 indicates no effect). Growing larvae in separate dishes or at different densities did not affect behavior, as demonstrated by largely overlapping SSMD distributions. However, larvae from different clutches exhibited significantly divergent SSMD distributions relative to control within-clutch comparisons (Figure 8A). For example, non-sibling larvae exhibited significantly different spontaneous movement frequency and dark flash response displacement, in contrast to siblings raised at identical or different densities (Figure 8B). Replicates of non-sibling comparisons generated greater numbers of p-values <0.05 and more divergent kernel density estimate peaks relative to other comparisons (Figure 8C). These results highlight the importance of comparing behavioral phenotypes within the same clutch.

**Figure 8.**
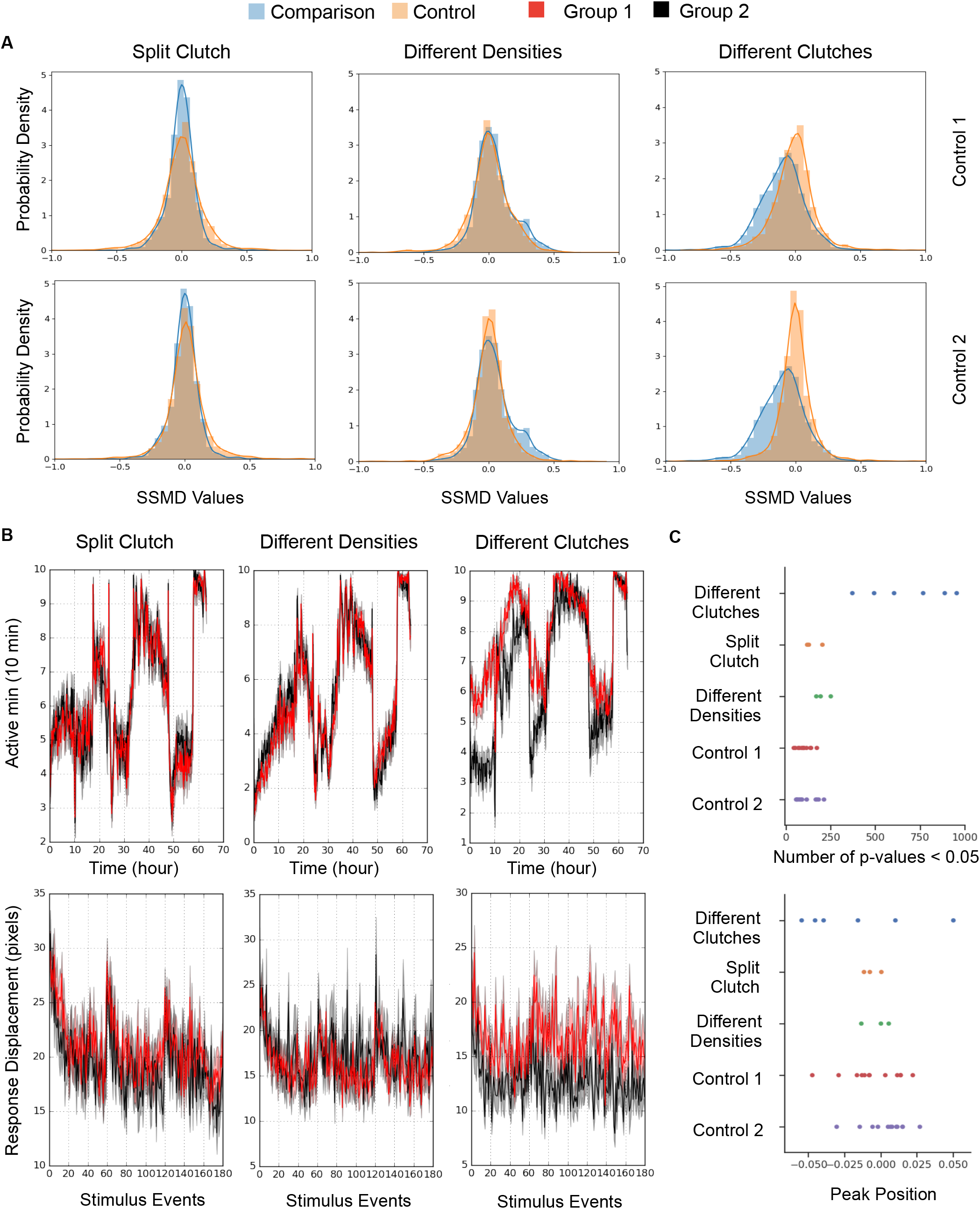
Quantification and Analysis of Wild Type Comparisons. **(A)** Probability density function of strictly standardized mean difference (SSMD) values for all behavioral metrics, according to comparisons outlined in Figure 7. *Left*, Split clutch comparison: Experimental N = 45 and 47 (Group 1 and Group 2), Control 1 N = 21 and 24, Control 2 N = 24 and 23. *Middle*, Different densities comparison: Experimental N = 47 and 43, Control 1 N = 23 and 23, Control 2 N = 21 and 23. *Right*, Different clutches comparison: Experimental N = 47 and 46, Control 1 N = 23 and 24, Control 2 N = 23 and 23. **(B)** Example graphs for two measures included in Figure 7A: movement frequency and dark flash response displacement. Split clutch comparison: Kruskal-Wallis p-value for the Movement Frequency (Active minutes) metric = 0.751. Kruskal-Wallis p-value for the dark flash Stimulus metric (Response Displacement) = 0.196. Different densities comparison: Kruskal-Wallis p-value for the Movement Frequency (Active minutes) metric = 0.415. Kruskal-Wallis p-value for the dark flash Stimulus metric (Response Displacement) = 0.993. Different clutches comparison: Kruskal-Wallis p-value for the Movement Frequency (Active minutes) metric = 9.04×10^−9^. Kruskal-Wallis p-value for the dark flash Stimulus metric (Response Displacement) = 6.05×10^−8^. **(D)** Number of p-values < 0.05 and the peak position of the kernel density estimation (KDE) curve for each comparison. 12 sets of comparisons with respective controls (Different clutch comparison: 4 independent comparisons and 2 replicates, split clutch comparison: 3 independent comparisons, different density comparison: 3 independent comparisons). The total number of p-values ranged from 1,951 to 1,996 depending on the comparison.

## 4 Discussion

The setup described above can test diverse zebrafish behaviors at high throughput and with minimal specialized expertise, equipment, and cost. In addition to baseline motion parameters, the system can assay prepulse inhibition (Figure 6) and responses to acoustic, visual, and thermal stimuli (Figure 5 and data not shown). We also incorporate a mini-projector to test additional visual behaviors such as the optomotor response (Figure 4). Our setup can thus accommodate more sophisticated visual assays including looming stimuli (Temizer et al., 2015), prey capture (Semmelhack et al., 2014), and decision-making based on dot coherence (Bahl and Engert, 2020). Because many components are commercially available, multiple boxes can be completely assembled within two days.

Our modular hardware design supports rapid adaptation for additional assays with adult animals or arena shapes beyond 96-well plate format. Modified camera/lens configurations can produce different resolutions or acquisition speeds. The circuit board design includes multiple BNC connectors capable of triggering or sampling from other devices. For example, these connectors can support optogenetic experiments (Oikonomou et al., 2019) as in the DanioVision system, or deliver electric shocks for conditioning assays (Valente et al., 2012), as in the ZebraBox system.

The software to execute and analyze experiments is also highly adaptable. Existing experiment events can be modified to yield new event types. For example, users can acquire extended movies (Command ID “2”) of a desired length or frame-rate by customizing the event type. These slow-speed movies provide opportunities for a wide range of new analyses that move beyond centroid/motion data, such as machine learning approaches to uncover phenotypes. This approach would have particular utility when monitoring older animals with more complex behaviors than larvae such as mating (Geng and Peterson, 2019). While our Python-based analyses measure motion parameters more comprehensively than any commercially available zebrafish analysis software, machine learning may distinguish additional classes of movements or responses. For example, we do not explicitly distinguish O-bends and C-bends from other movements, but used parameters such as motion velocity to separate responses. Indeed, our analysis pipeline (Figure 3) is currently based solely on motion quantification, but is also highly flexible. A large set of input options can be modified from default settings without coding (Supplementary Material). Furthermore, the code’s object-oriented and modular style permits independent modification of parameters such as the output graph format.

Using this new system, we assessed best practices for raising zebrafish larvae for behavior experiments (Figure 7, Figure 8). First, we found no effect of splitting a clutch across two petri dishes. While we routinely removed all debris from dishes during growth (see Methods), different levels of cleanliness may still influence behavior. Second, we found no effect of growth density up to 150 larvae in a 15×150 mm petri dish. Densities higher than 150 were not tested and may negatively impact growth. Third, and most critical, we found that wild-type animals from different clutches exhibited behavioral differences of similar magnitude to mutants with the strongest behavior phenotypes of 165 mutants (Thyme et al., 2019) versus their respective control siblings (Supplementary Figure 14). These results underscore the importance of comparing results within single clutches. We postulate that inter-clutch differences may contribute to variability in other contexts such as calcium imaging, where data is often collected from many parental pairs.

The zebrafish model continues to increase in popularity (Teame et al., 2019), while recent advances in genome editing technologies lower experimental barriers for non-traditional models. Our adaptable behavioral setup can monitor any small aquatic organism, particularly in multi-well format, and can thus accelerate discovery along both of these avenues. While neuroscientists likely represent the majority of users, our system can also serve as a powerful diagnostic tool for the development and function of other organs such as muscle (Maves, 2014). Finally, genome sequencing continues to link large numbers of genes to human disease (Schizophrenia Working Group of the Psychiatric Genomics Consortium, 2014; Satterstrom et al., 2020). The high throughput approaches outlined here will be critical to establish connections between disease-associated genes and decipher their neurobiological functions.

## Supporting information

Bill of Materials

Fabrication Files

Supplemental Software

Electronics Videos

## Author Contributions

WJ, BJG, ERS, and SBT contributed to design and construction of the behavior setup. SBT, WJ, and MDV wrote the manuscript. ERS built the majority of the LabVIEW software with contributions from MDV and SBT, and SBT built the majority of the Python software with contributions from MDV. BJG built the Arduino interface. SBT, WJ, and MDV conducted the experiments to demonstrate box functionality.

## 7 Acknowledgments

The authors thank Verdion Martina, Gretchen Kioschos, and Emma Jones for assistance in building behavior boxes, Verdion Martina for assisting with data collection, Emma Jones for helpful comments on figures, Ari Ginsparg for assisting with Python environments, the UAB fish facility staff for zebrafish care, the UAB research computing team for providing and maintaining the Cheaha cluster, and the UAB electronics and machine shop.

