## Supplementary figures and images for "A Customizable Low-Cost System for Massively Parallel Zebrafish Behavior Phenotyping"

### schematic.pdf

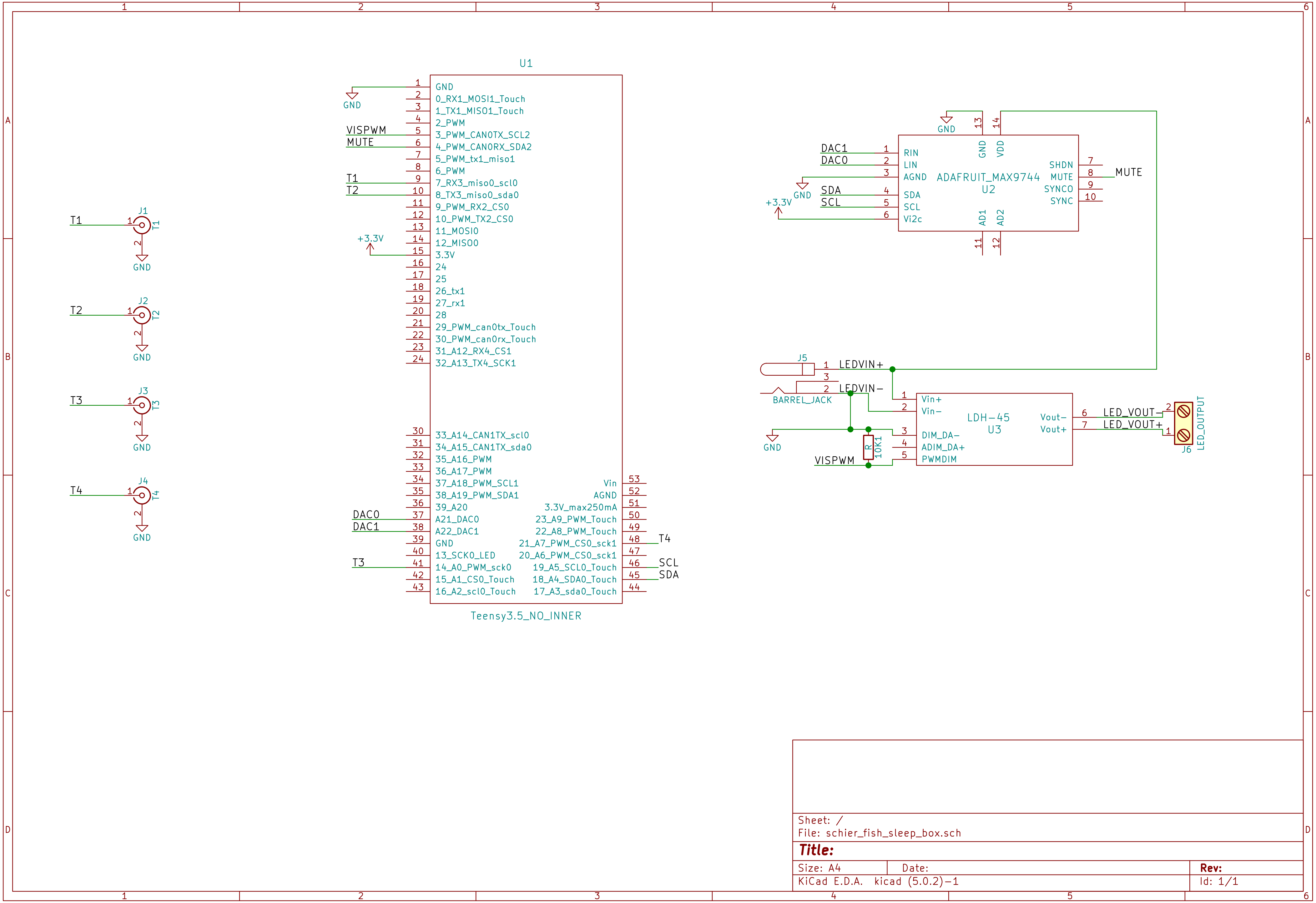
