## Supplemental Software for "A Customizable Low-Cost System for Massively Parallel Zebrafish Behavior Phenotyping": submission_script_gene_het_vs_gene_hom.slurm

```
#!/bin/bash
#SBATCH -p short # Partition to submit to
#SBATCH -n 1 # Number of cores requested
#SBATCH -N 1 # Ensure that all cores are on one machine
#SBATCH -t 600 # Runtime in minutes
#SBATCH --mem=64000 # Memory per cpu in MB (see also --mem-per-cpu)
#SBATCH -o linearmodel_gene_het_vs_gene_hom_%A_%a.out # Standard out goes to this file
#SBATCH -e hostname_%A_%a.err # Standard err goes to this filehostname
module load Anaconda3/5.3.1
cd outputfulldata_gene_het_vs_gene_hom
python /data/project/thymelab/ProcessMotion/processmotiondata.py -t "../testlog.timestamp1.Thu, Apr 16, 2020" -e "../eventsfile" -c "../testlog.centroid1.Thu, Apr 16, 2020" -d "../testlog.motion1.Thu, Apr 16, 2020" -m "../hsmovieThu, Apr 16, 2020_" -g "../gene_het_vs_gene_hom_scripted_inputgenotypeids" -s "../sectionsfile"  -j "../PlotParameters"
```
