## Supplemental Software for "A Customizable Low-Cost System for Massively Parallel Zebrafish Behavior Phenotyping": highspeedmovieanalysis.py

```
#!/usr/bin/python -tt
"""
highspeedmovieanalysis.py

The purpose of this script is analyze high-speed movies acquired from LabVIEW.
A movie is analyzed for motion and centroid data of fish in wells. Fish are 
identified in each well using the rois text file written from the LabVIEW software. 

Arguments
---------
-r: The rois text file produced from the LabVIEW program Generate ROIS.vi
-m: The high speed movie to be analyzed
-e: The events file used in the LabVIEW program Data Acquisition.vi
-p: The pixel threshold to identify a larvae. Default is 3.
-f: The frame rate of the high speed movie. Default is 285.

Outputs
-------
Motion data file (.motion2)
Centroid data file (.centroid2)
Last frame PNG image
Last frame PNG image with ROI numbering
Tracked lines PNG image
Mode PNG image
 
"""

import numpy as np
import cv2
from matplotlib.pyplot import imread
import glob,argparse
from scipy.stats import mode
import math
from PIL import Image, ImageFont, ImageDraw
from pathlib import Path
import datetime
from datetime import datetime as dt
import re
import glob

parser = argparse.ArgumentParser(description='loading for fish behavior files')
parser.add_argument('-r', type=str, action="store", dest="roisfile")
parser.add_argument('-m', type=str, action="store", dest="moviefile")
parser.add_argument('-e', type=str, action="store", dest="eventsfile")
parser.add_argument('-p', type=str, action="store", dest="pixThreshold", default=3)
parser.add_argument('-f', type=str, action="store", dest="frameRate", default=285)

args = parser.parse_args()
roisfile = args.roisfile
videoStream = args.moviefile
eventsfile = args.eventsfile
pixThreshold = args.pixThreshold
frameRate = args.frameRate

def calc_mode(deq, nump_arr):
    """
    This function calculates the mode image of a set of frames from a list of movies.

    Parameters
    ---------
    deq: list
       Gaussian-blurred frames to be used for mode image calculation.
    nump_arr: ndarray
       Array of zeros matching the resolution of the 1088x660 movie.

    Returns
    -------
    nump_arr: ndarray
       Mode image

    """
    for j,k in enumerate(nump_arr[:,0]): 
        nump_arr[j,:] = mode(np.array([x[j,:] for x in deq]))[0]
    return nump_arr

def imageMode(movielist, modename):
    """
    This function uses the calcMode function to obtain the mode image. 
    It then writes the image as a PNG to be viewed later.

    Parameters 
    ---------
    movielist: list
        A list containing a set of ~30 movie numbers
    modename: string
        A string specifying the range of movies used to create mode image.
        For instance, 1to30 would be the name for movies 1 through 30.

    Outputs
    -------
    Mode PNG image for each range of 30 images

    """

    moviedeq = []
    i2 = 0    
    for filenumber in movielist:
        cap = cv2.VideoCapture(glob.glob("*_" + str(filenumber) + ".avi")[0])
        ret,frame = cap.read()
        storedFrame = grayBlur(frame)
        totalFrames = 0
        while(cap.isOpened()):
            ret,frame = cap.read()
            if ret == False:
                break
            currentFrame = grayBlur(frame)
            if totalFrames < 50:
                if totalFrames % 3 == 0:
                    moviedeq.append(currentFrame)
            totalFrames += 1
            storedFrame = currentFrame 
        i2 += 1
    testing = calc_mode(moviedeq, np.zeros([660,1088]))
    cv2.imwrite("mode_" + modename + ".png", testing)
    cap.release()
    cv2.destroyAllWindows()


def diffImage(storedFrame,currentFrame,pixThreshold):
    """
    This function finds the pixel difference between the current frame and
    the previous frame. It then normalizes the image pixel values by dividing by 255

    Parameters
    ---------
    storedFrame: ndarray
        Previous frame that has been blurred by the grayBlur function.
    currentFrame: ndarray
        The current frame; blurred using the grayBlur function.
    pixThreshold: int
        Value for the pixel threshold. Default is 3.

    Returns
    -------
    diff: ndarray
        output array that has the same size and shape as the frames
 
    """

    diff = cv2.absdiff(storedFrame,currentFrame)
    _,diff = cv2.threshold(diff,pixThreshold,255,cv2.THRESH_BINARY)
    diff = diff / 255
    return diff

def trackdiffImage(storedFrame, currentFrame, pixThreshold = 7):
    """
    This function finds the pixel difference between the current frame and
    the previous frame. Similar to the diffImage function, but uses a higher threshold
    and does not normalize the diff array.

    Parameters
    ---------
    storedFrame: ndarray
        Previous frame that has been blurred by the grayBlur function.
    currentFrame: ndarray
        The current frame; blurred using the grayBlur function.
    pixThreshold: int
        Value for the pixel threshold. Default is 7.

    Returns
    -------
    diff: ndarray
        Output array that has the same size and shape as the frames
 
    """
    diff = cv2.absdiff(storedFrame,currentFrame)
    _,diff = cv2.threshold(diff,7,255,cv2.THRESH_BINARY)
    return diff

def Blur(image):
    """
    This function blurs a frame using the GaussionBlur function from opencv.
    
    Parameters
    ---------
    image: ndarray
        Frame to be blurred

    Returns
    -------
    Gaussian blurred image

    """

    return cv2.GaussianBlur(image,(7,7),0) 

def grayBlur(image):
    """
    This function blurs a RGB frame turned to gray 
    using the GaussionBlur function from opencv.
    
    Parameters
    ---------
    image: ndarray
        Frame to be blurred

    Returns
    -------
    Gaussian blurred gray image

    """

    gray = cv2.cvtColor(image, cv2.COLOR_BGR2GRAY)
    return cv2.GaussianBlur(gray,(7,7),0) 

def convertMaskToWeights(mask):
    """
    This function takes a 1088x660 array of ROI locations and converts it to an
    array of weights for binning.

    Parameters
    ---------
    mask: ndarray
        1088x660 array with numbering of ROIs

    Returns
    -------
    w: ndarray
        1-dimensional array of unique values
 
    """
    vals = np.unique(mask)
    for i in range(len(vals)):
        mask[mask==vals[i]]=i
    mask = mask.astype(int)
    w = mask.ravel() 
    return w

def loadmodeImage(modefilename):
    """
    This function determines if a mode image file exists then loads it as an array.

    Parameters
    ---------
    modefilename: string
         Name of mode image

    Returns
    -------
    e: ndarray
         Mode image as an array of values

    """
    try:
        e = imread(modefilename)
    except:
        exit('Cannot open mode file')
    return(e)

def faststrptime(val):
    """
    This function splits a teensy string for time values and converts them as
    a datetime object.

    Parameters
    ---------
    val: string
         String containing values for time teensy command
    
    Returns
    -------
    A datetime object 
 
    """
    splits1 = val.split("/")
    splits2 = splits1[2].split(":")
    return datetime.datetime(
        int(splits1[2][0:4]), # %Y
        int(splits1[0]), # %m
        int(splits1[1]), # %d
        int(splits2[0][4:len(splits2[0])]), # %H
        int(splits2[1]), # %M
        int(splits2[2][0:2]), # %s
    )

def makenumROIsimage():
    """
    This function determines the last frame of the last movie, then writes the
    well number in the center of each well.
    
    Outputs
    -------
    A PNG of the last frame
    A PNG of the wells numbered

    """

    num = 0
    for i,line in enumerate(glob.glob(videoStream)): 
        movienum = int(re.split(' |_|.avi', line)[4])
        if movienum > num:
                num = movienum
                filename = line

    myFrameNumber = frameRate
    cap = cv2.VideoCapture(filename)
    totalFrames = cap.get(cv2.CAP_PROP_FRAME_COUNT)

    if myFrameNumber >= 0 & myFrameNumber <= totalFrames:
       cap.set(cv2.CAP_PROP_POS_FRAMES,myFrameNumber)

    ret, frame = cap.read()
    cv2.imwrite("lastframe.png", frame)
    image = Image.open('lastframe.png')
    draw = ImageDraw.Draw(image)
    font = ImageFont.load_default()

    f = open(roisfile, 'r')
    lines = f.readlines()
    i = 1
    for line in lines:
       try:
               print(int(line.split(' ')[0]))
       except ValueError:
               continue
       x1 = int(line.split(' ')[0])
       y1 = int(line.split(' ')[1])
       x2 = int(line.split(' ')[2])
       y2 = int(line.split(' ')[3])

       midx = math.ceil((x1 + x2)/2)
       midy = math.ceil((y1 + y2)/2)

       draw.text((midx,midy), str(i), font=font)
       i += 1
    image.save('NumberedROIsImage.png')    

def main(pixThreshold,frameRate,videoStream):
    """
    The main function of the analysis script. Analyzes an individual movie for movement in well.

    Parameters
    ---------
    pixThreshold: int
        The pixel threshold value. Default is 3.
    frameRate: int
        The value pertaining to the number of frames for the movie. Default is 285.
    videoStream: string
        The name of the movie to be analyzed.

    Outputs
    -------
    A data file pertaining to movement in each well, called <videoStream>.motion2
    A data file pertaining to the centroids of each fish, called <videoStream>.centroid2
    A PNG file showing the centroids of the fish during the entire movie, called <videoStream>_trackedimagewithlines.png.

    """
    saveFreq = 4500 
    filenumber = videoStream.split('.')[0].split('_')[len(videoStream.split('.')[0].split('_'))-1]

    f = open(eventsfile, 'r')
    lines = f.readlines()
    numcounter = 0
    counter = 0
    fullcounter = 0
    movielist = []
    movielists =[]
    timestamp_list = []
    filteredlist = [] 
    startdate = "2020-02-26"
    
    for line in lines:
        TAPES = line.split('\t')
        if int(TAPES[2]) == 1 or int(TAPES[2]) == 2:
            filteredlist.append(line)
    
    for newline in filteredlist:
        TAPES = newline.split('\t')
        fullcounter +=1
        if int(TAPES[2]) == 2:
             timestamp_list.append(0)
             continue
        startdate2 = startdate.split("-")[1] + "/" + startdate.split("-")[2] + "/" + startdate.split("-")[0]
        dateplustime = startdate2 + TAPES[0][0:len(TAPES[0])]
        thistime = faststrptime(dateplustime)
        unixtimestamp = dt.timestamp(thistime)
        timestamp_list.append(int(unixtimestamp))

    i = 0    
    for element in timestamp_list:

        if i < (len(timestamp_list)-1) and timestamp_list[i+(counter-i)]-timestamp_list[i] >= 3600:
           counter += 1
           i = counter
           movielist.append(counter)
           
           if len(movielist) <= 15:
                numcounter = 0
                j = 0
                for step in movielist:
                    movielists[len(movielists)-1].append(movielist[j])
                    j += 1
                movielist = []
                continue   
           else:
                movielists.append(movielist)
                movielist = []
                numcounter = 0
                continue

        if i < (len(timestamp_list)-1) and timestamp_list[i+1]-timestamp_list[i] >= 3600:
           counter += 1
           i = counter
           movielist.append(counter)

           if len(movielist) <= 15:
                numcounter = 0
                j = 0
                for step in movielist:
                    movielists[len(movielists)-1].append(movielist[j])
                    j += 1
                movielist = []
                continue
           else:
                movielists.append(movielist)
                movielist = []
                numcounter = 0
                continue

        counter += 1
        numcounter += 1
        if element != 0:
             movielist.append(counter)
             i += 1
        
        if numcounter == 30:
            numcounter = 0
            movielists.append(movielist)
            movielist = []
        
        if i > (len(timestamp_list)-1):
            movielists.append(movielist)
            movielist = []
            numcounter = 0
            
    numendlists = counter - fullcounter
    first = len(movielists)-numendlists
    last = len(movielists)
    del movielists[first:last]
    
    for x in movielists:
        for y in x:
            if int(filenumber) == y:
                movielist = x

    modename = str(movielist[0]) + "to" + str(movielist[len(movielist)-1])
    imageMode(movielist, modename)
    modefilename = "mode_" + modename + ".png"
    try:
       imread(modefilename)
    except:
       imageMode(movielist,modename)

    e = loadmodeImage(modefilename)
    
    cmaxxs = []
    cminxs = []
    cmaxys = []
    cminys = []
    roimask = np.zeros((660,1088))
    f = open(roisfile, 'r')
    lines = f.readlines()
    i = 1
    i2 = 0
    for line in lines:
       try:
               print(int(line.split(' ')[0]))
       except ValueError:
               i2 += 1
               continue
       minx = int(line.split(' ')[0])
       miny = int(line.split(' ')[1])
       maxx = int(line.split(' ')[2])
       maxy = int(line.split(' ')[3])
       roimask[int(miny):int(maxy),int(minx):int(maxx)] = i
       cmaxxs.append(maxx)
       cmaxys.append(maxy)
       cminxs.append(minx)
       cminys.append(miny)
       i += 1
    print(roimask)
    numberofwells = i-1
    numberofcols = int(i2/2)
    numberofrows = int(numberofwells/numberofcols)
    cmaxxs.sort()
    cmaxys.sort()
    cminxs.sort()
    cminys.sort()
    roimaskweights = convertMaskToWeights(roimask)

    cap = cv2.VideoCapture(videoStream)

    cap.set(3,roimask.shape[1])
    cap.set(4,roimask.shape[0])
    
    ret,frame = cap.read()
    storedImage = np.array(e * 255, dtype = np.uint8)
    storedMode = Blur(storedImage)
    storedFrame = grayBlur(frame)
    cenData = np.zeros([ saveFreq, len(np.unique(roimaskweights))*2 -2])
    print(len(np.unique(roimaskweights))*2 -2)
    pixData = np.zeros([ saveFreq, len(np.unique(roimaskweights))])
    i = 0;
    totalFrames = 0
    while(cap.isOpened()):
        ret,frame = cap.read()
        if ret == False:
            break
        currentFrame = grayBlur(frame)
        diffpix = diffImage(storedFrame,currentFrame,pixThreshold)
        diff = trackdiffImage(storedMode,currentFrame)
        diff.dtype = np.uint8
        contours,hierarchy = cv2.findContours(diff, cv2.RETR_TREE,cv2.CHAIN_APPROX_NONE)
        MIN_THRESH = 20.0
        MIN_THRESH_P = 20.0
        roi_dict = {}
        for r in range(0,numberofwells):
            roi_dict[r+1] = []
        for cs in range(0,len(contours)):
            if cv2.contourArea(contours[cs]) < 1.0:
                continue
            if cv2.arcLength(contours[cs],True) < 1.0:
                continue
            if cv2.contourArea(contours[cs]) > MIN_THRESH or cv2.arcLength(contours[cs],True) > MIN_THRESH_P:
                M = cv2.moments(contours[cs])
                cX = int(M["m10"] / M["m00"])
                cY = int(M["m01"] / M["m00"])
                area = cv2.contourArea(contours[cs])
                perim = cv2.arcLength(contours[cs],True)
                if int(roimask[cY,cX]) == 0:
                   continue
                if not roi_dict[int(roimask[cY,cX])]:
                    roi_dict[int(roimask[cY,cX])].append((area*perim,cX,cY))
                else:
                    if roi_dict[int(roimask[cY,cX])][0][0] < area*perim:
                        roi_dict[int(roimask[cY,cX])][0] = (area*perim,cX,cY)

        pixcounts = []
        pixcounts = np.bincount(roimaskweights, weights=diffpix.ravel())
        pixData[i,:] = np.hstack((pixcounts))
        counts = []
        keys = roi_dict.keys()
        keys = sorted(keys)
        for k in keys:
            x = -10000
            y = -10000
            if roi_dict[k]:
                x = roi_dict[k][0][1]
                y = roi_dict[k][0][2]
            counts.append(x)
            counts.append(y)
            cv2.line(storedImage,(x,y),(x,y),(255,255,255),2)
        if i == 284:
            cv2.imwrite(videoStream + '_trackedimagewithlines_' + str(i) + ".png", storedImage)
        cenData[i,:] = np.asarray(counts)
        totalFrames += 1
        storedFrame = currentFrame
        i += 1

    file = open(videoStream + ".centroid2",'w')
    for x in range(0,frameRate):
        for y in range(0,numberofwells*2):
            file.write(str(int(cenData[x,:][y])) + '\n')
    pixData = pixData[:i,:]
    pixData = pixData[:,1:] 
    file = open(videoStream + ".motion2",'w')
    for x in range(0,frameRate):
        for y in range(0,numberofwells):
            file.write(str(int(pixData[x,:][y])) + '\n')

    cap.release()
    cv2.destroyAllWindows()
    try:
        image = Image.open('lastframe.png')
    except:
        makenumROIsimage()

main(pixThreshold,frameRate,videoStream)
```
