## Supplemental Software for "A Customizable Low-Cost System for Massively Parallel Zebrafish Behavior Phenotyping": makeOMRgrating.py

```
"""
makeOMRgrating.py

This script creates a series of PNG files that afterwards
need to be fused into a movie (e.g. with ImageJ)

Outputs
-------
PNG images of gratings for movie creation
 
"""

import numpy as np
import png

frameRate = 24 # This needs to be set to the same value later on movie creation!

framesForward = 0*frameRate # For how many frames should the grating move forward
framesHold = 0*frameRate # In-between stationary period if desired
framesBackwards = 20*framesForward # Backwards movement

# How many pixels wide is a black+light stripe - should be adjusted such that
# The period of the grating will be 1cm on screen - that creates maximal response from fish
gratingPeriod = 100
#movie resolution
gratingWidth = 800
gratingHeight = 600


timeForward = np.arange(framesForward)
timeHold = np.ones(framesHold) * timeForward.max()
timeBackwards = timeForward.max() - np.arange(framesBackwards)

time = np.hstack(timeForward)


def MakeName(index,basename='grating'):
    """
    This function assumes the existence of a sub-directory named 'grating'
    """
    return basename+'_'+str(index).zfill(3)+'.png'


def OMRGrating(t,period,vDark=150,vBright=250,width=640,height=480):
    """
    Creates a square grating frame. In general larger contrast = larger response, but too strong edges sometimes cause problems

    Parameters
    ----------
    t: int
      Time point
    period: int
      Space between each grating stripe
    vDark: int
      Pixel value of black stripe
    vBright: int
      Pixel value of white stripe
    width: int 
      Width of the image
    height: int
      Height of the image

    Returns
    -------
    grating: ndarray
      Array with size widthxheight to create grating image

    """

    # Set movement per time-bin such that the grating will move one
    # Period with frameRate number of frames
    global frameRate
    move = period*t//frameRate
    grating = np.zeros((height,width))
    for i in range(height):
        j = (i+move) % height
        j = j % period
        if j < period//2:
            grating[i,:] = vDark
        else:
            grating[i,:] = vBright
    return grating

# Make the movie
w = png.Writer(gratingWidth,gratingHeight,greyscale=True)
for i,t in enumerate(time):
    grat = OMRGrating(t,gratingPeriod,width=gratingWidth,height=gratingHeight)
    try:
        f = open(MakeName(i),'wb')
    except:
        print('File error')
        break
    else:
        w.write(f,grat)
        f.close()
```
