## Supplemental Software for "A Customizable Low-Cost System for Massively Parallel Zebrafish Behavior Phenotyping": makeslurmfiles.py

```
#!/usr/bin/python

import os,sys,glob,argparse,shutil

# Input arguments
parser = argparse.ArgumentParser(description='options for slurm file generation')
parser.add_argument('-module', type=str, action="store", dest="module", default="Anaconda3")
parser.add_argument('-partition', type=str, action="store", dest="partition", default="short")
parser.add_argument('-statsfile', type=str, action="store", dest="statsfile", default="linearmodel") # prefix for this file
parser.add_argument('-gfile', type=str, action="store", dest="gfile", default="genotyping")
parser.add_argument('-efile', type=str, action="store", dest="efile", default="../fulltestrun_final_01_27_2020")
parser.add_argument('-prefix', type=str, action="store", dest="prefix", default="../testlog") # slow-speed data prefix
parser.add_argument('-hprefix', type=str, action="store", dest="hprefix", default="../hsmovie") # high-speed data prefix
parser.add_argument('-pfile', type=str, action="store", dest="pfile", default="../PlotParameters")
parser.add_argument('-esfile', type=str, action="store", dest="esfile", default="../sectionsfile")
parser.add_argument('-scriptpath', type=str, action="store", dest="scriptpath", default="/data/project/thymelab/Newest_Behavior_Scripts/versionforROI/") # path for the analysis suite
parser.add_argument('-other', type=str, action="store", dest="other", default="") # args to add at the end. YOU CANNOT INCLUDE THE INITIAL DASH FOR A NEW ARG WITH ARGPARSER, WILL BE ADDED BY THIS SCRIPT"

args = parser.parse_args()
module = args.module
partition = args.partition
statsfile = args.statsfile
gfile = args.gfile
efile = args.efile
prefix = args.prefix
hprefix = args.hprefix
pfile = args.pfile
esfile = args.esfile
scriptpath = args.scriptpath
other = args.other

# This logic determins what is a control and what is the "mutant" comparison. Add identifiers as needed.
logic = {				 "wt":0,
					"wtwt":0,
					"hetwt":2,
					"wthet":1,
					"wtandhet":1,
					"hetandwt":1,
					"hethet":3,
					"het":3,
					"homwt":5,
					"wthom":4,
					"hethom":6,
					"homhet":7,
					"hom":7,
					"homhom":8,
					"drug":1,
					"dmso":0
				}

def make_slurm_file(fd1, fd2, date, hfile):
	ffile = open('submission_script_' + fd1 + "_vs_" + fd2 + '.slurm', 'w')
	ffile.write("#!/bin/bash\n")
	ffile.write("#SBATCH -p " + partition + " # Partition to submit to\n")
	ffile.write("#SBATCH -n 1 # Number of cores requested\n")
	ffile.write("#SBATCH -N 1 # Ensure that all cores are on one machine\n")
	ffile.write("#SBATCH -t 600 # Runtime in minutes\n")
	ffile.write("#SBATCH --mem=64000 # Memory per cpu in MB (see also --mem-per-cpu)\n")
	ffile.write("#SBATCH -o " + statsfile + "_" + fd1 + "_vs_" + fd2 + "_%A_%a.out # Standard out goes to this file\n")
	ffile.write("#SBATCH -e hostname_%A_%a.err # Standard err goes to this filehostname\n")
	ffile.write("module load " + module + "\n")
	ffile.write("cd outputfulldata_" + fd1 + "_vs_" + fd2 + '\n')
	if os.path.exists("outputfulldata_" + fd1 + "_vs_" + fd2):
		print("Output directories already generated.")
	else:
		os.mkdir("outputfulldata_" + fd1 + "_vs_" + fd2)
	ffile.write("python " + scriptpath + "processmotiondata.py -t ")
	ffile.write("\"" + prefix + ".timestamp1." + date)
	ffile.write("\" -e \"" + efile + "\" -c \"" + prefix + ".centroid1." + date)
	ffile.write("\" -d \"" + prefix + ".motion1." + date)
	ffile.write("\" -m \"" + hprefix + date)
	ffile.write("_\" -g \../"" + hfile)
	ffile.write("\" -s \"" + esfile)
	ffile.write("\" -j \"" + pfile + "\"")
	if other != "":
		ffile.write(" -" + other)
	return 'submission_script_' + fd1 + "_vs_" + fd2 + '.slurm'

genofile = open(gfile, 'r')
lines = genofile.readlines()
dirnames = []
iddict = {}
for line in lines:
	if line.startswith('*'):
		destdir = line.strip().split(':')[0][1:]
		itype = destdir.split('_')[1]
		dirnames.append(destdir)
		ids = line.strip().split(':')[1].strip()
		iddict[itype] = ids
for file1 in glob.glob('*timestamp1*'):
	date = file1.split('.')[2]
sfile = open("jobsubmission.sh", 'w')
sfile.write("#!/bin/bash\n")
for fd1 in dirnames:
	for fd2 in dirnames:
		if fd1.split('_')[0] == fd2.split('_')[0]: # Check if same gene
			if fd1 != fd2:
				testfile = 'submission_script_' + fd1 + "_vs_" + fd2 + '.slurm'
				testfileo = 'submission_script_' + fd2 + "_vs_" + fd1 + '.slurm'
				if os.path.exists(testfile) or os.path.exists(testfileo):
					continue
				type1 = fd1.split('_')[1]
				type2 = fd2.split('_')[1]
				if logic[type1] > logic[type2]:
					hfile = fd2 + "_vs_" + fd1 + "_scripted_inputgenotypeids"
					ifile = open(hfile, 'w')
					ifile.write("controlgroup_" + type2 + ":" + iddict[type2])
					ifile.write("\ntestgroup_" + type1 + ":" + iddict[type1] + "\n")
					fname2r = make_slurm_file(fd2, fd1, date, hfile)
				else:
					hfile = fd1 + "_vs_" + fd2 + "_scripted_inputgenotypeids"
					ifile = open(hfile, 'w')
					ifile.write("controlgroup_" + type1 + ":" + iddict[type1])
					ifile.write("\ntestgroup_" + type2 + ":" + iddict[type2] + "\n")
					fname2r = make_slurm_file(fd1, fd2, date, hfile)
				sfile.write("sbatch ")
				sfile.write(fname2r)
				sfile.write("\nsleep 3\n")

os.system("chmod +x jobsubmission.sh")
```
