## Supplemental Software for "A Customizable Low-Cost System for Massively Parallel Zebrafish Behavior Phenotyping": png.py

```
#!/usr/bin/env python

from __future__ import print_function

# png.py - PNG encoder/decoder in pure Python
#
# Copyright (C) 2006 Johann C. Rocholl <>
# Portions Copyright (C) 2009 David Jones <>
# And probably portions Copyright (C) 2006 Nicko van Someren <>
#
# Original concept by Johann C. Rocholl.
#
# LICENCE (MIT)
#
# Permission is hereby granted, free of charge, to any person
# obtaining a copy of this software and associated documentation files
# (the "Software"), to deal in the Software without restriction,
# including without limitation the rights to use, copy, modify, merge,
# publish, distribute, sublicense, and/or sell copies of the Software,
# and to permit persons to whom the Software is furnished to do so,
# subject to the following conditions:
#
# The above copyright notice and this permission notice shall be
# included in all copies or substantial portions of the Software.
#
# THE SOFTWARE IS PROVIDED "AS IS", WITHOUT WARRANTY OF ANY KIND,
# EXPRESS OR IMPLIED, INCLUDING BUT NOT LIMITED TO THE WARRANTIES OF
# MERCHANTABILITY, FITNESS FOR A PARTICULAR PURPOSE AND
# NONINFRINGEMENT. IN NO EVENT SHALL THE AUTHORS OR COPYRIGHT HOLDERS
# BE LIABLE FOR ANY CLAIM, DAMAGES OR OTHER LIABILITY, WHETHER IN AN
# ACTION OF CONTRACT, TORT OR OTHERWISE, ARISING FROM, OUT OF OR IN
# CONNECTION WITH THE SOFTWARE OR THE USE OR OTHER DEALINGS IN THE
# SOFTWARE.

"""
Pure Python PNG Reader/Writer

This Python module implements support for PNG images (see PNG
specification at http://www.w3.org/TR/2003/REC-PNG-20031110/ ). It reads
and writes PNG files with all allowable bit depths
(1/2/4/8/16/24/32/48/64 bits per pixel) and colour combinations:
greyscale (1/2/4/8/16 bit); RGB, RGBA, LA (greyscale with alpha) with
8/16 bits per channel; colour mapped images (1/2/4/8 bit).
Adam7 interlacing is supported for reading and
writing.  A number of optional chunks can be specified (when writing)
and understood (when reading): ``tRNS``, ``bKGD``, ``gAMA``.

For help, type ``import png; help(png)`` in your python interpreter.

A good place to start is the :class:`Reader` and :class:`Writer`
classes.

Requires Python 2.3.  Limited support is available for Python 2.2, but
not everything works.  Best with Python 2.4 and higher.  Installation is
trivial, but see the ``README.txt`` file (with the source distribution)
for details.

This file can also be used as a command-line utility to convert
`Netpbm <http://netpbm.sourceforge.net/>`_ PNM files to PNG, and the
reverse conversion from PNG to PNM. The interface is similar to that
of the ``pnmtopng`` program from Netpbm.  Type ``python png.py --help``
at the shell prompt for usage and a list of options.

A note on spelling and terminology
----------------------------------

Generally British English spelling is used in the documentation.  So
that's "greyscale" and "colour".  This not only matches the author's
native language, it's also used by the PNG specification.

The major colour models supported by PNG (and hence by PyPNG) are:
greyscale, RGB, greyscale--alpha, RGB--alpha.  These are sometimes
referred to using the abbreviations: L, RGB, LA, RGBA.  In this case
each letter abbreviates a single channel: *L* is for Luminance or Luma
or Lightness which is the channel used in greyscale images; *R*, *G*,
*B* stand for Red, Green, Blue, the components of a colour image; *A*
stands for Alpha, the opacity channel (used for transparency effects,
but higher values are more opaque, so it makes sense to call it 
opacity).

A note on formats
-----------------

When getting pixel data out of this module (reading) and presenting
data to this module (writing) there are a number of ways the data could
be represented as a Python value.  Generally this module uses one of
three formats called "flat row flat pixel", "boxed row flat pixel", and
"boxed row boxed pixel".  Basically the concern is whether each pixel
and each row comes in its own little tuple (box), or not.

Consider an image that is 3 pixels wide by 2 pixels high, and each pixel
has RGB components:

Boxed row flat pixel::

  list([R,G,B, R,G,B, R,G,B],
       [R,G,B, R,G,B, R,G,B])

Each row appears as its own list, but the pixels are flattened so
that three values for one pixel simply follow the three values for
the previous pixel.  This is the most common format used, because it
provides a good compromise between space and convenience.  PyPNG regards
itself as at liberty to replace any sequence type with any sufficiently
compatible other sequence type; in practice each row is an array (from
the array module), and the outer list is sometimes an iterator rather
than an explicit list (so that streaming is possible).

Flat row flat pixel::

  [R,G,B, R,G,B, R,G,B,
   R,G,B, R,G,B, R,G,B]

The entire image is one single giant sequence of colour values.
Generally an array will be used (to save space), not a list.

Boxed row boxed pixel::

  list([ (R,G,B), (R,G,B), (R,G,B) ],
       [ (R,G,B), (R,G,B), (R,G,B) ])

Each row appears in its own list, but each pixel also appears in its own
tuple.  A serious memory burn in Python.

In all cases the top row comes first, and for each row the pixels are
ordered from left-to-right.  Within a pixel the values appear in the
order, R-G-B-A (or L-A for greyscale--alpha).

There is a fourth format, mentioned because it is used internally,
is close to what lies inside a PNG file itself, and has some support
from the public API.  This format is called packed.  When packed,
each row is a sequence of bytes (integers from 0 to 255), just as
it is before PNG scanline filtering is applied.  When the bit depth
is 8 this is essentially the same as boxed row flat pixel; when the
bit depth is less than 8, several pixels are packed into each byte;
when the bit depth is 16 (the only value more than 8 that is supported
by the PNG image format) each pixel value is decomposed into 2 bytes
(and `packed` is a misnomer).  This format is used by the
:meth:`Writer.write_packed` method.  It isn't usually a convenient
format, but may be just right if the source data for the PNG image
comes from something that uses a similar format (for example, 1-bit
BMPs, or another PNG file).

And now, my famous members
--------------------------
"""

__version__ = "0.0.18"

import itertools
import math
# http://www.python.org/doc/2.4.4/lib/module-operator.html
import operator
import struct
import sys
# http://www.python.org/doc/2.4.4/lib/module-warnings.html
import warnings
import zlib

from array import array
from functools import reduce

try:
    # `cpngfilters` is a Cython module: it must be compiled by
    # Cython for this import to work.
    # If this import does work, then it overrides pure-python
    # filtering functions defined later in this file (see `class
    # pngfilters`).
    import cpngfilters as pngfilters
except ImportError:
    pass


__all__ = ['Image', 'Reader', 'Writer', 'write_chunks', 'from_array']


# The PNG signature.
# http://www.w3.org/TR/PNG/#5PNG-file-signature
_signature = struct.pack('8B', 137, 80, 78, 71, 13, 10, 26, 10)

_adam7 = ((0, 0, 8, 8),
          (4, 0, 8, 8),
          (0, 4, 4, 8),
          (2, 0, 4, 4),
          (0, 2, 2, 4),
          (1, 0, 2, 2),
          (0, 1, 1, 2))

def group(s, n):
    # See http://www.python.org/doc/2.6/library/functions.html#zip
    return list(zip(*[iter(s)]*n))

def isarray(x):
    return isinstance(x, array)

def tostring(row):
    return row.tostring()

def interleave_planes(ipixels, apixels, ipsize, apsize):
    """
    Interleave (colour) planes, e.g. RGB + A = RGBA.

    Return an array of pixels consisting of the `ipsize` elements of
    data from each pixel in `ipixels` followed by the `apsize` elements
    of data from each pixel in `apixels`.  Conventionally `ipixels`
    and `apixels` are byte arrays so the sizes are bytes, but it
    actually works with any arrays of the same type.  The returned
    array is the same type as the input arrays which should be the
    same type as each other.
    """

    itotal = len(ipixels)
    atotal = len(apixels)
    newtotal = itotal + atotal
    newpsize = ipsize + apsize
    # Set up the output buffer
    # See http://www.python.org/doc/2.4.4/lib/module-array.html#l2h-1356
    out = array(ipixels.typecode)
    # It's annoying that there is no cheap way to set the array size :-(
    out.extend(ipixels)
    out.extend(apixels)
    # Interleave in the pixel data
    for i in range(ipsize):
        out[i:newtotal:newpsize] = ipixels[i:itotal:ipsize]
    for i in range(apsize):
        out[i+ipsize:newtotal:newpsize] = apixels[i:atotal:apsize]
    return out

def check_palette(palette):
    """Check a palette argument (to the :class:`Writer` class)
    for validity.  Returns the palette as a list if okay; raises an
    exception otherwise.
    """

    # None is the default and is allowed.
    if palette is None:
        return None

    p = list(palette)
    if not (0 < len(p) <= 256):
        raise ValueError("a palette must have between 1 and 256 entries")
    seen_triple = False
    for i,t in enumerate(p):
        if len(t) not in (3,4):
            raise ValueError(
              "palette entry %d: entries must be 3- or 4-tuples." % i)
        if len(t) == 3:
            seen_triple = True
        if seen_triple and len(t) == 4:
            raise ValueError(
              "palette entry %d: all 4-tuples must precede all 3-tuples" % i)
        for x in t:
            if int(x) != x or not(0 <= x <= 255):
                raise ValueError(
                  "palette entry %d: values must be integer: 0 <= x <= 255" % i)
    return p

def check_sizes(size, width, height):
    """Check that these arguments, in supplied, are consistent.
    Return a (width, height) pair.
    """

    if not size:
        return width, height

    if len(size) != 2:
        raise ValueError(
          "size argument should be a pair (width, height)")
    if width is not None and width != size[0]:
        raise ValueError(
          "size[0] (%r) and width (%r) should match when both are used."
            % (size[0], width))
    if height is not None and height != size[1]:
        raise ValueError(
          "size[1] (%r) and height (%r) should match when both are used."
            % (size[1], height))
    return size

def check_color(c, greyscale, which):
    """Checks that a colour argument for transparent or
    background options is the right form.  Returns the colour
    (which, if it's a bar integer, is "corrected" to a 1-tuple).
    """

    if c is None:
        return c
    if greyscale:
        try:
            len(c)
        except TypeError:
            c = (c,)
        if len(c) != 1:
            raise ValueError("%s for greyscale must be 1-tuple" %
                which)
        if not isinteger(c[0]):
            raise ValueError(
                "%s colour for greyscale must be integer" % which)
    else:
        if not (len(c) == 3 and
                isinteger(c[0]) and
                isinteger(c[1]) and
                isinteger(c[2])):
            raise ValueError(
                "%s colour must be a triple of integers" % which)
    return c

class Error(Exception):
    def __str__(self):
        return self.__class__.__name__ + ': ' + ' '.join(self.args)

class FormatError(Error):
    """Problem with input file format.  In other words, PNG file does
    not conform to the specification in some way and is invalid.
    """

class ChunkError(FormatError):
    pass


class Writer:
    """
    PNG encoder in pure Python.
    """

    def __init__(self, width=None, height=None,
                 size=None,
                 greyscale=False,
                 alpha=False,
                 bitdepth=8,
                 palette=None,
                 transparent=None,
                 background=None,
                 gamma=None,
                 compression=None,
                 interlace=False,
                 bytes_per_sample=None, # deprecated
                 planes=None,
                 colormap=None,
                 maxval=None,
                 chunk_limit=2**20,
                 x_pixels_per_unit = None,
                 y_pixels_per_unit = None,
                 unit_is_meter = False):
        """
        Create a PNG encoder object.

        Arguments:

        width, height
          Image size in pixels, as two separate arguments.
        size
          Image size (w,h) in pixels, as single argument.
        greyscale
          Input data is greyscale, not RGB.
        alpha
          Input data has alpha channel (RGBA or LA).
        bitdepth
          Bit depth: from 1 to 16.
        palette
          Create a palette for a colour mapped image (colour type 3).
        transparent
          Specify a transparent colour (create a ``tRNS`` chunk).
        background
          Specify a default background colour (create a ``bKGD`` chunk).
        gamma
          Specify a gamma value (create a ``gAMA`` chunk).
        compression
          zlib compression level: 0 (none) to 9 (more compressed);
          default: -1 or None.
        interlace
          Create an interlaced image.
        chunk_limit
          Write multiple ``IDAT`` chunks to save memory.
        x_pixels_per_unit
          Number of pixels a unit along the x axis (write a
          `pHYs` chunk).
        y_pixels_per_unit
          Number of pixels a unit along the y axis (write a
          `pHYs` chunk). Along with `x_pixel_unit`, this gives
          the pixel size ratio.
        unit_is_meter
          `True` to indicate that the unit (for the `pHYs`
          chunk) is metre.

        The image size (in pixels) can be specified either by using the
        `width` and `height` arguments, or with the single `size`
        argument.  If `size` is used it should be a pair (*width*,
        *height*).

        `greyscale` and `alpha` are booleans that specify whether
        an image is greyscale (or colour), and whether it has an
        alpha channel (or not).

        `bitdepth` specifies the bit depth of the source pixel values.
        Each source pixel value must be an integer between 0 and
        ``2**bitdepth-1``.  For example, 8-bit images have values
        between 0 and 255.  PNG only stores images with bit depths of
        1,2,4,8, or 16.  When `bitdepth` is not one of these values,
        the next highest valid bit depth is selected, and an ``sBIT``
        (significant bits) chunk is generated that specifies the
        original precision of the source image.  In this case the
        supplied pixel values will be rescaled to fit the range of
        the selected bit depth.

        The details of which bit depth / colour model combinations the
        PNG file format supports directly, are somewhat arcane
        (refer to the PNG specification for full details).  Briefly:
        "small" bit depths (1,2,4) are only allowed with greyscale and
        colour mapped images; colour mapped images cannot have bit depth
        16.

        For colour mapped images (in other words, when the `palette`
        argument is specified) the `bitdepth` argument must match one of
        the valid PNG bit depths: 1, 2, 4, or 8.  (It is valid to have a
        PNG image with a palette and an ``sBIT`` chunk, but the meaning
        is slightly different; it would be awkward to press the
        `bitdepth` argument into service for this.)

        The `palette` option, when specified, causes a colour
        mapped image to be created: the PNG colour type is set to 3;
        `greyscale` must not be set; `alpha` must not be set;
        `transparent` must not be set; the bit depth must be 1,2,4,
        or 8.  When a colour mapped image is created, the pixel values
        are palette indexes and the `bitdepth` argument specifies the
        size of these indexes (not the size of the colour values in
        the palette).

        The palette argument value should be a sequence of 3- or
        4-tuples.  3-tuples specify RGB palette entries; 4-tuples
        specify RGBA palette entries.  If both 4-tuples and 3-tuples
        appear in the sequence then all the 4-tuples must come
        before all the 3-tuples.  A ``PLTE`` chunk is created; if there
        are 4-tuples then a ``tRNS`` chunk is created as well.  The
        ``PLTE`` chunk will contain all the RGB triples in the same
        sequence; the ``tRNS`` chunk will contain the alpha channel for
        all the 4-tuples, in the same sequence.  Palette entries
        are always 8-bit.

        If specified, the `transparent` and `background` parameters must
        be a tuple with three integer values for red, green, blue, or
        a simple integer (or singleton tuple) for a greyscale image.

        If specified, the `gamma` parameter must be a positive number
        (generally, a `float`).  A ``gAMA`` chunk will be created.
        Note that this will not change the values of the pixels as
        they appear in the PNG file, they are assumed to have already
        been converted appropriately for the gamma specified.

        The `compression` argument specifies the compression level to
        be used by the ``zlib`` module.  Values from 1 to 9 specify
        compression, with 9 being "more compressed" (usually smaller
        and slower, but it doesn't always work out that way).  0 means
        no compression.  -1 and ``None`` both mean that the default
        level of compession will be picked by the ``zlib`` module
        (which is generally acceptable).

        If `interlace` is true then an interlaced image is created
        (using PNG's so far only interace method, *Adam7*).  This does
        not affect how the pixels should be presented to the encoder,
        rather it changes how they are arranged into the PNG file.
        On slow connexions interlaced images can be partially decoded
        by the browser to give a rough view of the image that is
        successively refined as more image data appears.

        .. note ::

          Enabling the `interlace` option requires the entire image
          to be processed in working memory.

        `chunk_limit` is used to limit the amount of memory used whilst
        compressing the image.  In order to avoid using large amounts of
        memory, multiple ``IDAT`` chunks may be created.
        """

        # At the moment the `planes` argument is ignored;
        # its purpose is to act as a dummy so that
        # ``Writer(x, y, **info)`` works, where `info` is a dictionary
        # returned by Reader.read and friends.
        # Ditto for `colormap`.

        width, height = check_sizes(size, width, height)
        del size

        if width <= 0 or height <= 0:
            raise ValueError("width and height must be greater than zero")
        if not isinteger(width) or not isinteger(height):
            raise ValueError("width and height must be integers")
        # http://www.w3.org/TR/PNG/#7Integers-and-byte-order
        if width > 2**32-1 or height > 2**32-1:
            raise ValueError("width and height cannot exceed 2**32-1")

        if alpha and transparent is not None:
            raise ValueError(
                "transparent colour not allowed with alpha channel")

        if bytes_per_sample is not None:
            warnings.warn('please use bitdepth instead of bytes_per_sample',
                          DeprecationWarning)
            if bytes_per_sample not in (0.125, 0.25, 0.5, 1, 2):
                raise ValueError(
                    "bytes per sample must be .125, .25, .5, 1, or 2")
            bitdepth = int(8*bytes_per_sample)
        del bytes_per_sample
        if not isinteger(bitdepth) or bitdepth < 1 or 16 < bitdepth:
            raise ValueError("bitdepth (%r) must be a positive integer <= 16" %
              bitdepth)

        self.rescale = None
        palette = check_palette(palette)
        if palette:
            if bitdepth not in (1,2,4,8):
                raise ValueError("with palette, bitdepth must be 1, 2, 4, or 8")
            if transparent is not None:
                raise ValueError("transparent and palette not compatible")
            if alpha:
                raise ValueError("alpha and palette not compatible")
            if greyscale:
                raise ValueError("greyscale and palette not compatible")
        else:
            # No palette, check for sBIT chunk generation.
            if alpha or not greyscale:
                if bitdepth not in (8,16):
                    targetbitdepth = (8,16)[bitdepth > 8]
                    self.rescale = (bitdepth, targetbitdepth)
                    bitdepth = targetbitdepth
                    del targetbitdepth
            else:
                assert greyscale
                assert not alpha
                if bitdepth not in (1,2,4,8,16):
                    if bitdepth > 8:
                        targetbitdepth = 16
                    elif bitdepth == 3:
                        targetbitdepth = 4
                    else:
                        assert bitdepth in (5,6,7)
                        targetbitdepth = 8
                    self.rescale = (bitdepth, targetbitdepth)
                    bitdepth = targetbitdepth
                    del targetbitdepth

        if bitdepth < 8 and (alpha or not greyscale and not palette):
            raise ValueError(
              "bitdepth < 8 only permitted with greyscale or palette")
        if bitdepth > 8 and palette:
            raise ValueError(
                "bit depth must be 8 or less for images with palette")

        transparent = check_color(transparent, greyscale, 'transparent')
        background = check_color(background, greyscale, 'background')

        # It's important that the true boolean values (greyscale, alpha,
        # colormap, interlace) are converted to bool because Iverson's
        # convention is relied upon later on.
        self.width = width
        self.height = height
        self.transparent = transparent
        self.background = background
        self.gamma = gamma
        self.greyscale = bool(greyscale)
        self.alpha = bool(alpha)
        self.colormap = bool(palette)
        self.bitdepth = int(bitdepth)
        self.compression = compression
        self.chunk_limit = chunk_limit
        self.interlace = bool(interlace)
        self.palette = palette
        self.x_pixels_per_unit = x_pixels_per_unit
        self.y_pixels_per_unit = y_pixels_per_unit
        self.unit_is_meter = bool(unit_is_meter)

        self.color_type = 4*self.alpha + 2*(not greyscale) + 1*self.colormap
        assert self.color_type in (0,2,3,4,6)

        self.color_planes = (3,1)[self.greyscale or self.colormap]
        self.planes = self.color_planes + self.alpha
        # :todo: fix for bitdepth < 8
        self.psize = (self.bitdepth/8) * self.planes

    def make_palette(self):
        """Create the byte sequences for a ``PLTE`` and if necessary a
        ``tRNS`` chunk.  Returned as a pair (*p*, *t*).  *t* will be
        ``None`` if no ``tRNS`` chunk is necessary.
        """

        p = array('B')
        t = array('B')

        for x in self.palette:
            p.extend(x[0:3])
            if len(x) > 3:
                t.append(x[3])
        p = tostring(p)
        t = tostring(t)
        if t:
            return p,t
        return p,None

    def write(self, outfile, rows):
        """Write a PNG image to the output file.  `rows` should be
        an iterable that yields each row in boxed row flat pixel
        format.  The rows should be the rows of the original image,
        so there should be ``self.height`` rows of ``self.width *
        self.planes`` values.  If `interlace` is specified (when
        creating the instance), then an interlaced PNG file will
        be written.  Supply the rows in the normal image order;
        the interlacing is carried out internally.

        .. note ::

          Interlacing will require the entire image to be in working
          memory.
        """

        if self.interlace:
            fmt = 'BH'[self.bitdepth > 8]
            a = array(fmt, itertools.chain(*rows))
            return self.write_array(outfile, a)

        nrows = self.write_passes(outfile, rows)
        if nrows != self.height:
            raise ValueError(
              "rows supplied (%d) does not match height (%d)" %
              (nrows, self.height))

    def write_passes(self, outfile, rows, packed=False):
        """
        Write a PNG image to the output file.

        Most users are expected to find the :meth:`write` or
        :meth:`write_array` method more convenient.
        
        The rows should be given to this method in the order that
        they appear in the output file.  For straightlaced images,
        this is the usual top to bottom ordering, but for interlaced
        images the rows should have already been interlaced before
        passing them to this function.

        `rows` should be an iterable that yields each row.  When
        `packed` is ``False`` the rows should be in boxed row flat pixel
        format; when `packed` is ``True`` each row should be a packed
        sequence of bytes.
        """

        # http://www.w3.org/TR/PNG/#5PNG-file-signature
        outfile.write(_signature)

        # http://www.w3.org/TR/PNG/#11IHDR
        write_chunk(outfile, b'IHDR',
                    struct.pack("!2I5B", self.width, self.height,
                                self.bitdepth, self.color_type,
                                0, 0, self.interlace))

        # See :chunk:order
        # http://www.w3.org/TR/PNG/#11gAMA
        if self.gamma is not None:
            write_chunk(outfile, b'gAMA',
                        struct.pack("!L", int(round(self.gamma*1e5))))

        # See :chunk:order
        # http://www.w3.org/TR/PNG/#11sBIT
        if self.rescale:
            write_chunk(outfile, b'sBIT',
                struct.pack('%dB' % self.planes,
                            *[self.rescale[0]]*self.planes))
        
        # :chunk:order: Without a palette (PLTE chunk), ordering is
        # relatively relaxed.  With one, gAMA chunk must precede PLTE
        # chunk which must precede tRNS and bKGD.
        # See http://www.w3.org/TR/PNG/#5ChunkOrdering
        if self.palette:
            p,t = self.make_palette()
            write_chunk(outfile, b'PLTE', p)
            if t:
                # tRNS chunk is optional. Only needed if palette entries
                # have alpha.
                write_chunk(outfile, b'tRNS', t)

        # http://www.w3.org/TR/PNG/#11tRNS
        if self.transparent is not None:
            if self.greyscale:
                write_chunk(outfile, b'tRNS',
                            struct.pack("!1H", *self.transparent))
            else:
                write_chunk(outfile, b'tRNS',
                            struct.pack("!3H", *self.transparent))

        # http://www.w3.org/TR/PNG/#11bKGD
        if self.background is not None:
            if self.greyscale:
                write_chunk(outfile, b'bKGD',
                            struct.pack("!1H", *self.background))
            else:
                write_chunk(outfile, b'bKGD',
                            struct.pack("!3H", *self.background))

        # http://www.w3.org/TR/PNG/#11pHYs
        if self.x_pixels_per_unit is not None and self.y_pixels_per_unit is not None:
            tup = (self.x_pixels_per_unit, self.y_pixels_per_unit, int(self.unit_is_meter))
            write_chunk(outfile, b'pHYs', struct.pack("!LLB",*tup))

        # http://www.w3.org/TR/PNG/#11IDAT
        if self.compression is not None:
            compressor = zlib.compressobj(self.compression)
        else:
            compressor = zlib.compressobj()

        # Choose an extend function based on the bitdepth.  The extend
        # function packs/decomposes the pixel values into bytes and
        # stuffs them onto the data array.
        data = array('B')
        if self.bitdepth == 8 or packed:
            extend = data.extend
        elif self.bitdepth == 16:
            # Decompose into bytes
            def extend(sl):
                fmt = '!%dH' % len(sl)
                data.extend(array('B', struct.pack(fmt, *sl)))
        else:
            # Pack into bytes
            assert self.bitdepth < 8
            # samples per byte
            spb = int(8/self.bitdepth)
            def extend(sl):
                a = array('B', sl)
                # Adding padding bytes so we can group into a whole
                # number of spb-tuples.
                l = float(len(a))
                extra = math.ceil(l / float(spb))*spb - l
                a.extend([0]*int(extra))
                # Pack into bytes
                l = group(a, spb)
                l = [reduce(lambda x,y:
                                           (x << self.bitdepth) + y, e) for e in l]
                data.extend(l)
        if self.rescale:
            oldextend = extend
            factor = \
              float(2**self.rescale[1]-1) / float(2**self.rescale[0]-1)
            def extend(sl):
                oldextend([int(round(factor*x)) for x in sl])

        # Build the first row, testing mostly to see if we need to
        # changed the extend function to cope with NumPy integer types
        # (they cause our ordinary definition of extend to fail, so we
        # wrap it).  See
        # http://code.google.com/p/pypng/issues/detail?id=44
        enumrows = enumerate(rows)
        del rows

        # First row's filter type.
        data.append(0)
        # :todo: Certain exceptions in the call to ``.next()`` or the
        # following try would indicate no row data supplied.
        # Should catch.
        i,row = next(enumrows)
        try:
            # If this fails...
            extend(row)
        except:
            # ... try a version that converts the values to int first.
            # Not only does this work for the (slightly broken) NumPy
            # types, there are probably lots of other, unknown, "nearly"
            # int types it works for.
            def wrapmapint(f):
                return lambda sl: f([int(x) for x in sl])
            extend = wrapmapint(extend)
            del wrapmapint
            extend(row)

        for i,row in enumrows:
            # Add "None" filter type.  Currently, it's essential that
            # this filter type be used for every scanline as we do not
            # mark the first row of a reduced pass image; that means we
            # could accidentally compute the wrong filtered scanline if
            # we used "up", "average", or "paeth" on such a line.
            data.append(0)
            extend(row)
            if len(data) > self.chunk_limit:
                compressed = compressor.compress(tostring(data))
                if len(compressed):
                    write_chunk(outfile, b'IDAT', compressed)
                # Because of our very witty definition of ``extend``,
                # above, we must re-use the same ``data`` object.  Hence
                # we use ``del`` to empty this one, rather than create a
                # fresh one (which would be my natural FP instinct).
                del data[:]
        if len(data):
            compressed = compressor.compress(tostring(data))
        else:
            compressed = b''
        flushed = compressor.flush()
        if len(compressed) or len(flushed):
            write_chunk(outfile, b'IDAT', compressed + flushed)
        # http://www.w3.org/TR/PNG/#11IEND
        write_chunk(outfile, b'IEND')
        return i+1

    def write_array(self, outfile, pixels):
        """
        Write an array in flat row flat pixel format as a PNG file on
        the output file.  See also :meth:`write` method.
        """

        if self.interlace:
            self.write_passes(outfile, self.array_scanlines_interlace(pixels))
        else:
            self.write_passes(outfile, self.array_scanlines(pixels))

    def write_packed(self, outfile, rows):
        """
        Write PNG file to `outfile`.  The pixel data comes from `rows`
        which should be in boxed row packed format.  Each row should be
        a sequence of packed bytes.

        Technically, this method does work for interlaced images but it
        is best avoided.  For interlaced images, the rows should be
        presented in the order that they appear in the file.

        This method should not be used when the source image bit depth
        is not one naturally supported by PNG; the bit depth should be
        1, 2, 4, 8, or 16.
        """

        if self.rescale:
            raise Error("write_packed method not suitable for bit depth %d" %
              self.rescale[0])
        return self.write_passes(outfile, rows, packed=True)

    def convert_pnm(self, infile, outfile):
        """
        Convert a PNM file containing raw pixel data into a PNG file
        with the parameters set in the writer object.  Works for
        (binary) PGM, PPM, and PAM formats.
        """

        if self.interlace:
            pixels = array('B')
            pixels.fromfile(infile,
                            (self.bitdepth/8) * self.color_planes *
                            self.width * self.height)
            self.write_passes(outfile, self.array_scanlines_interlace(pixels))
        else:
            self.write_passes(outfile, self.file_scanlines(infile))

    def convert_ppm_and_pgm(self, ppmfile, pgmfile, outfile):
        """
        Convert a PPM and PGM file containing raw pixel data into a
        PNG outfile with the parameters set in the writer object.
        """
        pixels = array('B')
        pixels.fromfile(ppmfile,
                        (self.bitdepth/8) * self.color_planes *
                        self.width * self.height)
        apixels = array('B')
        apixels.fromfile(pgmfile,
                         (self.bitdepth/8) *
                         self.width * self.height)
        pixels = interleave_planes(pixels, apixels,
                                   (self.bitdepth/8) * self.color_planes,
                                   (self.bitdepth/8))
        if self.interlace:
            self.write_passes(outfile, self.array_scanlines_interlace(pixels))
        else:
            self.write_passes(outfile, self.array_scanlines(pixels))

    def file_scanlines(self, infile):
        """
        Generates boxed rows in flat pixel format, from the input file
        `infile`.  It assumes that the input file is in a "Netpbm-like"
        binary format, and is positioned at the beginning of the first
        pixel.  The number of pixels to read is taken from the image
        dimensions (`width`, `height`, `planes`) and the number of bytes
        per value is implied by the image `bitdepth`.
        """

        # Values per row
        vpr = self.width * self.planes
        row_bytes = vpr
        if self.bitdepth > 8:
            assert self.bitdepth == 16
            row_bytes *= 2
            fmt = '>%dH' % vpr
            def line():
                return array('H', struct.unpack(fmt, infile.read(row_bytes)))
        else:
            def line():
                scanline = array('B', infile.read(row_bytes))
                return scanline
        for y in range(self.height):
            yield line()

    def array_scanlines(self, pixels):
        """
        Generates boxed rows (flat pixels) from flat rows (flat pixels)
        in an array.
        """

        # Values per row
        vpr = self.width * self.planes
        stop = 0
        for y in range(self.height):
            start = stop
            stop = start + vpr
            yield pixels[start:stop]

    def array_scanlines_interlace(self, pixels):
        """
        Generator for interlaced scanlines from an array.  `pixels` is
        the full source image in flat row flat pixel format.  The
        generator yields each scanline of the reduced passes in turn, in
        boxed row flat pixel format.
        """

        # http://www.w3.org/TR/PNG/#8InterlaceMethods
        # Array type.
        fmt = 'BH'[self.bitdepth > 8]
        # Value per row
        vpr = self.width * self.planes
        for xstart, ystart, xstep, ystep in _adam7:
            if xstart >= self.width:
                continue
            # Pixels per row (of reduced image)
            ppr = int(math.ceil((self.width-xstart)/float(xstep)))
            # number of values in reduced image row.
            row_len = ppr*self.planes
            for y in range(ystart, self.height, ystep):
                if xstep == 1:
                    offset = y * vpr
                    yield pixels[offset:offset+vpr]
                else:
                    row = array(fmt)
                    # There's no easier way to set the length of an array
                    row.extend(pixels[0:row_len])
                    offset = y * vpr + xstart * self.planes
                    end_offset = (y+1) * vpr
                    skip = self.planes * xstep
                    for i in range(self.planes):
                        row[i::self.planes] = \
                            pixels[offset+i:end_offset:skip]
                    yield row

def write_chunk(outfile, tag, data=b''):
    """
    Write a PNG chunk to the output file, including length and
    checksum.
    """

    # http://www.w3.org/TR/PNG/#5Chunk-layout
    outfile.write(struct.pack("!I", len(data)))
    outfile.write(tag)
    outfile.write(data)
    checksum = zlib.crc32(tag)
    checksum = zlib.crc32(data, checksum)
    checksum &= 2**32-1
    outfile.write(struct.pack("!I", checksum))

def write_chunks(out, chunks):
    """Create a PNG file by writing out the chunks."""

    out.write(_signature)
    for chunk in chunks:
        write_chunk(out, *chunk)

def filter_scanline(type, line, fo, prev=None):
    """Apply a scanline filter to a scanline.  `type` specifies the
    filter type (0 to 4); `line` specifies the current (unfiltered)
    scanline as a sequence of bytes; `prev` specifies the previous
    (unfiltered) scanline as a sequence of bytes. `fo` specifies the
    filter offset; normally this is size of a pixel in bytes (the number
    of bytes per sample times the number of channels), but when this is
    < 1 (for bit depths < 8) then the filter offset is 1.
    """

    assert 0 <= type < 5

    # The output array.  Which, pathetically, we extend one-byte at a
    # time (fortunately this is linear).
    out = array('B', [type])

    def sub():
        ai = -fo
        for x in line:
            if ai >= 0:
                x = (x - line[ai]) & 0xff
            out.append(x)
            ai += 1
    def up():
        for i,x in enumerate(line):
            x = (x - prev[i]) & 0xff
            out.append(x)
    def average():
        ai = -fo
        for i,x in enumerate(line):
            if ai >= 0:
                x = (x - ((line[ai] + prev[i]) >> 1)) & 0xff
            else:
                x = (x - (prev[i] >> 1)) & 0xff
            out.append(x)
            ai += 1
    def paeth():
        # http://www.w3.org/TR/PNG/#9Filter-type-4-Paeth
        ai = -fo # also used for ci
        for i,x in enumerate(line):
            a = 0
            b = prev[i]
            c = 0

            if ai >= 0:
                a = line[ai]
                c = prev[ai]
            p = a + b - c
            pa = abs(p - a)
            pb = abs(p - b)
            pc = abs(p - c)
            if pa <= pb and pa <= pc:
                Pr = a
            elif pb <= pc:
                Pr = b
            else:
                Pr = c

            x = (x - Pr) & 0xff
            out.append(x)
            ai += 1

    if not prev:
        # We're on the first line.  Some of the filters can be reduced
        # to simpler cases which makes handling the line "off the top"
        # of the image simpler.  "up" becomes "none"; "paeth" becomes
        # "left" (non-trivial, but true). "average" needs to be handled
        # specially.
        if type == 2: # "up"
            type = 0
        elif type == 3:
            prev = [0]*len(line)
        elif type == 4: # "paeth"
            type = 1
    if type == 0:
        out.extend(line)
    elif type == 1:
        sub()
    elif type == 2:
        up()
    elif type == 3:
        average()
    else: # type == 4
        paeth()
    return out


def from_array(a, mode=None, info={}):
    """Create a PNG :class:`Image` object from a 2- or 3-dimensional
    array.  One application of this function is easy PIL-style saving:
    ``png.from_array(pixels, 'L').save('foo.png')``.

    Unless they are specified using the *info* parameter, the PNG's
    height and width are taken from the array size.  For a 3 dimensional
    array the first axis is the height; the second axis is the width;
    and the third axis is the channel number.  Thus an RGB image that is
    16 pixels high and 8 wide will use an array that is 16x8x3.  For 2
    dimensional arrays the first axis is the height, but the second axis
    is ``width*channels``, so an RGB image that is 16 pixels high and 8
    wide will use a 2-dimensional array that is 16x24 (each row will be
    8*3 = 24 sample values).

    *mode* is a string that specifies the image colour format in a
    PIL-style mode.  It can be:

    ``'L'``
      greyscale (1 channel)
    ``'LA'``
      greyscale with alpha (2 channel)
    ``'RGB'``
      colour image (3 channel)
    ``'RGBA'``
      colour image with alpha (4 channel)

    The mode string can also specify the bit depth (overriding how this
    function normally derives the bit depth, see below).  Appending
    ``';16'`` to the mode will cause the PNG to be 16 bits per channel;
    any decimal from 1 to 16 can be used to specify the bit depth.

    When a 2-dimensional array is used *mode* determines how many
    channels the image has, and so allows the width to be derived from
    the second array dimension.

    The array is expected to be a ``numpy`` array, but it can be any
    suitable Python sequence.  For example, a list of lists can be used:
    ``png.from_array([[0, 255, 0], [255, 0, 255]], 'L')``.  The exact
    rules are: ``len(a)`` gives the first dimension, height;
    ``len(a[0])`` gives the second dimension; ``len(a[0][0])`` gives the
    third dimension, unless an exception is raised in which case a
    2-dimensional array is assumed.  It's slightly more complicated than
    that because an iterator of rows can be used, and it all still
    works.  Using an iterator allows data to be streamed efficiently.

    The bit depth of the PNG is normally taken from the array element's
    datatype (but if *mode* specifies a bitdepth then that is used
    instead).  The array element's datatype is determined in a way which
    is supposed to work both for ``numpy`` arrays and for Python
    ``array.array`` objects.  A 1 byte datatype will give a bit depth of
    8, a 2 byte datatype will give a bit depth of 16.  If the datatype
    does not have an implicit size, for example it is a plain Python
    list of lists, as above, then a default of 8 is used.

    The *info* parameter is a dictionary that can be used to specify
    metadata (in the same style as the arguments to the
    :class:`png.Writer` class).  For this function the keys that are
    useful are:
    
    height
      overrides the height derived from the array dimensions and allows
      *a* to be an iterable.
    width
      overrides the width derived from the array dimensions.
    bitdepth
      overrides the bit depth derived from the element datatype (but
      must match *mode* if that also specifies a bit depth).

    Generally anything specified in the
    *info* dictionary will override any implicit choices that this
    function would otherwise make, but must match any explicit ones.
    For example, if the *info* dictionary has a ``greyscale`` key then
    this must be true when mode is ``'L'`` or ``'LA'`` and false when
    mode is ``'RGB'`` or ``'RGBA'``.
    """

    # We abuse the *info* parameter by modifying it.  Take a copy here.
    # (Also typechecks *info* to some extent).
    info = dict(info)

    # Syntax check mode string.
    bitdepth = None
    try:
        # Assign the 'L' or 'RGBA' part to `gotmode`.
        if mode.startswith('L'):
            gotmode = 'L'
            mode = mode[1:]
        elif mode.startswith('RGB'):
            gotmode = 'RGB'
            mode = mode[3:]
        else:
            raise Error()
        if mode.startswith('A'):
            gotmode += 'A'
            mode = mode[1:]

        # Skip any optional ';'
        while mode.startswith(';'):
            mode = mode[1:]

        # Parse optional bitdepth
        if mode:
            try:
                bitdepth = int(mode)
            except (TypeError, ValueError):
                raise Error()
    except Error:
        raise Error("mode string should be 'RGB' or 'L;16' or similar.")
    mode = gotmode

    # Get bitdepth from *mode* if possible.
    if bitdepth:
        if info.get('bitdepth') and bitdepth != info['bitdepth']:
            raise Error("mode bitdepth (%d) should match info bitdepth (%d)." %
              (bitdepth, info['bitdepth']))
        info['bitdepth'] = bitdepth

    # Fill in and/or check entries in *info*.
    # Dimensions.
    if 'size' in info:
        # Check width, height, size all match where used.
        for dimension,axis in [('width', 0), ('height', 1)]:
            if dimension in info:
                if info[dimension] != info['size'][axis]:
                    raise Error(
                      "info[%r] should match info['size'][%r]." %
                      (dimension, axis))
        info['width'],info['height'] = info['size']
    if 'height' not in info:
        try:
            l = len(a)
        except TypeError:
            raise Error(
              "len(a) does not work, supply info['height'] instead.")
        info['height'] = l
    # Colour format.
    if 'greyscale' in info:
        if bool(info['greyscale']) != ('L' in mode):
            raise Error("info['greyscale'] should match mode.")
    info['greyscale'] = 'L' in mode
    if 'alpha' in info:
        if bool(info['alpha']) != ('A' in mode):
            raise Error("info['alpha'] should match mode.")
    info['alpha'] = 'A' in mode

    planes = len(mode)
    if 'planes' in info:
        if info['planes'] != planes:
            raise Error("info['planes'] should match mode.")

    # In order to work out whether we the array is 2D or 3D we need its
    # first row, which requires that we take a copy of its iterator.
    # We may also need the first row to derive width and bitdepth.
    a,t = itertools.tee(a)
    row = next(t)
    del t
    try:
        row[0][0]
        threed = True
        testelement = row[0]
    except (IndexError, TypeError):
        threed = False
        testelement = row
    if 'width' not in info:
        if threed:
            width = len(row)
        else:
            width = len(row) // planes
        info['width'] = width

    if threed:
        # Flatten the threed rows
        a = (itertools.chain.from_iterable(x) for x in a)

    if 'bitdepth' not in info:
        try:
            dtype = testelement.dtype
            # goto the "else:" clause.  Sorry.
        except AttributeError:
            try:
                # Try a Python array.array.
                bitdepth = 8 * testelement.itemsize
            except AttributeError:
                # We can't determine it from the array element's
                # datatype, use a default of 8.
                bitdepth = 8
        else:
            # If we got here without exception, we now assume that
            # the array is a numpy array.
            if dtype.kind == 'b':
                bitdepth = 1
            else:
                bitdepth = 8 * dtype.itemsize
        info['bitdepth'] = bitdepth

    for thing in 'width height bitdepth greyscale alpha'.split():
        assert thing in info
    return Image(a, info)

# So that refugee's from PIL feel more at home.  Not documented.
fromarray = from_array

class Image:
    """A PNG image.  You can create an :class:`Image` object from
    an array of pixels by calling :meth:`png.from_array`.  It can be
    saved to disk with the :meth:`save` method.
    """

    def __init__(self, rows, info):
        """
        .. note ::
        
          The constructor is not public.  Please do not call it.
        """
        
        self.rows = rows
        self.info = info

    def save(self, file):
        """Save the image to *file*.  If *file* looks like an open file
        descriptor then it is used, otherwise it is treated as a
        filename and a fresh file is opened.

        In general, you can only call this method once; after it has
        been called the first time and the PNG image has been saved, the
        source data will have been streamed, and cannot be streamed
        again.
        """

        w = Writer(**self.info)

        try:
            file.write
            def close(): pass
        except AttributeError:
            file = open(file, 'wb')
            def close(): file.close()

        try:
            w.write(file, self.rows)
        finally:
            close()

class _readable:
    """
    A simple file-like interface for strings and arrays.
    """

    def __init__(self, buf):
        self.buf = buf
        self.offset = 0

    def read(self, n):
        r = self.buf[self.offset:self.offset+n]
        if isarray(r):
            r = r.tostring()
        self.offset += n
        return r

try:
    str(b'dummy', 'ascii')
except TypeError:
    as_str = str
else:
    def as_str(x):
        return str(x, 'ascii')

class Reader:
    """
    PNG decoder in pure Python.
    """

    def __init__(self, _guess=None, **kw):
        """
        Create a PNG decoder object.

        The constructor expects exactly one keyword argument. If you
        supply a positional argument instead, it will guess the input
        type. You can choose among the following keyword arguments:

        filename
          Name of input file (a PNG file).
        file
          A file-like object (object with a read() method).
        bytes
          ``array`` or ``string`` with PNG data.

        """
        if ((_guess is not None and len(kw) != 0) or
            (_guess is None and len(kw) != 1)):
            raise TypeError("Reader() takes exactly 1 argument")

        # Will be the first 8 bytes, later on.  See validate_signature.
        self.signature = None
        self.transparent = None
        # A pair of (len,type) if a chunk has been read but its data and
        # checksum have not (in other words the file position is just
        # past the 4 bytes that specify the chunk type).  See preamble
        # method for how this is used.
        self.atchunk = None

        if _guess is not None:
            if isarray(_guess):
                kw["bytes"] = _guess
            elif isinstance(_guess, str):
                kw["filename"] = _guess
            elif hasattr(_guess, 'read'):
                kw["file"] = _guess

        if "filename" in kw:
            self.file = open(kw["filename"], "rb")
        elif "file" in kw:
            self.file = kw["file"]
        elif "bytes" in kw:
            self.file = _readable(kw["bytes"])
        else:
            raise TypeError("expecting filename, file or bytes array")


    def chunk(self, seek=None, lenient=False):
        """
        Read the next PNG chunk from the input file; returns a
        (*type*, *data*) tuple.  *type* is the chunk's type as a
        byte string (all PNG chunk types are 4 bytes long).
        *data* is the chunk's data content, as a byte string.

        If the optional `seek` argument is
        specified then it will keep reading chunks until it either runs
        out of file or finds the type specified by the argument.  Note
        that in general the order of chunks in PNGs is unspecified, so
        using `seek` can cause you to miss chunks.

        If the optional `lenient` argument evaluates to `True`,
        checksum failures will raise warnings rather than exceptions.
        """

        self.validate_signature()

        while True:
            # http://www.w3.org/TR/PNG/#5Chunk-layout
            if not self.atchunk:
                self.atchunk = self.chunklentype()
            length, type = self.atchunk
            self.atchunk = None
            data = self.file.read(length)
            if len(data) != length:
                raise ChunkError('Chunk %s too short for required %i octets.'
                  % (type, length))
            checksum = self.file.read(4)
            if len(checksum) != 4:
                raise ChunkError('Chunk %s too short for checksum.' % type)
            if seek and type != seek:
                continue
            verify = zlib.crc32(type)
            verify = zlib.crc32(data, verify)
            # Whether the output from zlib.crc32 is signed or not varies
            # according to hideous implementation details, see
            # http://bugs.python.org/issue1202 .
            # We coerce it to be positive here (in a way which works on
            # Python 2.3 and older).
            verify &= 2**32 - 1
            verify = struct.pack('!I', verify)
            if checksum != verify:
                (a, ) = struct.unpack('!I', checksum)
                (b, ) = struct.unpack('!I', verify)
                message = "Checksum error in %s chunk: 0x%08X != 0x%08X." % (type, a, b)
                if lenient:
                    warnings.warn(message, RuntimeWarning)
                else:
                    raise ChunkError(message)
            return type, data

    def chunks(self):
        """Return an iterator that will yield each chunk as a
        (*chunktype*, *content*) pair.
        """

        while True:
            t,v = self.chunk()
            yield t,v
            if t == b'IEND':
                break

    def undo_filter(self, filter_type, scanline, previous):
        """Undo the filter for a scanline.  `scanline` is a sequence of
        bytes that does not include the initial filter type byte.
        `previous` is decoded previous scanline (for straightlaced
        images this is the previous pixel row, but for interlaced
        images, it is the previous scanline in the reduced image, which
        in general is not the previous pixel row in the final image).
        When there is no previous scanline (the first row of a
        straightlaced image, or the first row in one of the passes in an
        interlaced image), then this argument should be ``None``.

        The scanline will have the effects of filtering removed, and the
        result will be returned as a fresh sequence of bytes.
        """

        # :todo: Would it be better to update scanline in place?
        # Yes, with the Cython extension making the undo_filter fast,
        # updating scanline inplace makes the code 3 times faster
        # (reading 50 images of 800x800 went from 40s to 16s)
        result = scanline

        if filter_type == 0:
            return result

        if filter_type not in (1,2,3,4):
            raise FormatError('Invalid PNG Filter Type.'
              '  See http://www.w3.org/TR/2003/REC-PNG-20031110/#9Filters .')

        # Filter unit.  The stride from one pixel to the corresponding
        # byte from the previous pixel.  Normally this is the pixel
        # size in bytes, but when this is smaller than 1, the previous
        # byte is used instead.
        fu = max(1, self.psize)

        # For the first line of a pass, synthesize a dummy previous
        # line.  An alternative approach would be to observe that on the
        # first line 'up' is the same as 'null', 'paeth' is the same
        # as 'sub', with only 'average' requiring any special case.
        if not previous:
            previous = array('B', [0]*len(scanline))

        def sub():
            """Undo sub filter."""

            ai = 0
            # Loop starts at index fu.  Observe that the initial part
            # of the result is already filled in correctly with
            # scanline.
            for i in range(fu, len(result)):
                x = scanline[i]
                a = result[ai]
                result[i] = (x + a) & 0xff
                ai += 1

        def up():
            """Undo up filter."""

            for i in range(len(result)):
                x = scanline[i]
                b = previous[i]
                result[i] = (x + b) & 0xff

        def average():
            """Undo average filter."""

            ai = -fu
            for i in range(len(result)):
                x = scanline[i]
                if ai < 0:
                    a = 0
                else:
                    a = result[ai]
                b = previous[i]
                result[i] = (x + ((a + b) >> 1)) & 0xff
                ai += 1

        def paeth():
            """Undo Paeth filter."""

            # Also used for ci.
            ai = -fu
            for i in range(len(result)):
                x = scanline[i]
                if ai < 0:
                    a = c = 0
                else:
                    a = result[ai]
                    c = previous[ai]
                b = previous[i]
                p = a + b - c
                pa = abs(p - a)
                pb = abs(p - b)
                pc = abs(p - c)
                if pa <= pb and pa <= pc:
                    pr = a
                elif pb <= pc:
                    pr = b
                else:
                    pr = c
                result[i] = (x + pr) & 0xff
                ai += 1

        # Call appropriate filter algorithm.  Note that 0 has already
        # been dealt with.
        (None,
         pngfilters.undo_filter_sub,
         pngfilters.undo_filter_up,
         pngfilters.undo_filter_average,
         pngfilters.undo_filter_paeth)[filter_type](fu, scanline, previous, result)
        return result

    def deinterlace(self, raw):
        """
        Read raw pixel data, undo filters, deinterlace, and flatten.
        Return in flat row flat pixel format.
        """

        # Values per row (of the target image)
        vpr = self.width * self.planes

        # Make a result array, and make it big enough.  Interleaving
        # writes to the output array randomly (well, not quite), so the
        # entire output array must be in memory.
        fmt = 'BH'[self.bitdepth > 8]
        a = array(fmt, [0]*vpr*self.height)
        source_offset = 0

        for xstart, ystart, xstep, ystep in _adam7:
            if xstart >= self.width:
                continue
            # The previous (reconstructed) scanline.  None at the
            # beginning of a pass to indicate that there is no previous
            # line.
            recon = None
            # Pixels per row (reduced pass image)
            ppr = int(math.ceil((self.width-xstart)/float(xstep)))
            # Row size in bytes for this pass.
            row_size = int(math.ceil(self.psize * ppr))
            for y in range(ystart, self.height, ystep):
                filter_type = raw[source_offset]
                source_offset += 1
                scanline = raw[source_offset:source_offset+row_size]
                source_offset += row_size
                recon = self.undo_filter(filter_type, scanline, recon)
                # Convert so that there is one element per pixel value
                flat = self.serialtoflat(recon, ppr)
                if xstep == 1:
                    assert xstart == 0
                    offset = y * vpr
                    a[offset:offset+vpr] = flat
                else:
                    offset = y * vpr + xstart * self.planes
                    end_offset = (y+1) * vpr
                    skip = self.planes * xstep
                    for i in range(self.planes):
                        a[offset+i:end_offset:skip] = \
                            flat[i::self.planes]
        return a

    def iterboxed(self, rows):
        """Iterator that yields each scanline in boxed row flat pixel
        format.  `rows` should be an iterator that yields the bytes of
        each row in turn.
        """

        def asvalues(raw):
            """Convert a row of raw bytes into a flat row.  Result will
            be a freshly allocated object, not shared with
            argument.
            """

            if self.bitdepth == 8:
                return array('B', raw)
            if self.bitdepth == 16:
                raw = tostring(raw)
                return array('H', struct.unpack('!%dH' % (len(raw)//2), raw))
            assert self.bitdepth < 8
            width = self.width
            # Samples per byte
            spb = 8//self.bitdepth
            out = array('B')
            mask = 2**self.bitdepth - 1
            shifts = [self.bitdepth * i
                for i in reversed(list(range(spb)))]
            for o in raw:
                out.extend([mask&(o>>i) for i in shifts])
            return out[:width]

        return map(asvalues, rows)

    def serialtoflat(self, bytes, width=None):
        """Convert serial format (byte stream) pixel data to flat row
        flat pixel.
        """

        if self.bitdepth == 8:
            return bytes
        if self.bitdepth == 16:
            bytes = tostring(bytes)
            return array('H',
              struct.unpack('!%dH' % (len(bytes)//2), bytes))
        assert self.bitdepth < 8
        if width is None:
            width = self.width
        # Samples per byte
        spb = 8//self.bitdepth
        out = array('B')
        mask = 2**self.bitdepth - 1
        shifts = list(map(self.bitdepth.__mul__, reversed(list(range(spb)))))
        l = width
        for o in bytes:
            out.extend([(mask&(o>>s)) for s in shifts][:l])
            l -= spb
            if l <= 0:
                l = width
        return out

    def iterstraight(self, raw):
        """Iterator that undoes the effect of filtering, and yields
        each row in serialised format (as a sequence of bytes).
        Assumes input is straightlaced.  `raw` should be an iterable
        that yields the raw bytes in chunks of arbitrary size.
        """

        # length of row, in bytes
        rb = self.row_bytes
        a = array('B')
        # The previous (reconstructed) scanline.  None indicates first
        # line of image.
        recon = None
        for some in raw:
            a.extend(some)
            while len(a) >= rb + 1:
                filter_type = a[0]
                scanline = a[1:rb+1]
                del a[:rb+1]
                recon = self.undo_filter(filter_type, scanline, recon)
                yield recon
        if len(a) != 0:
            # :file:format We get here with a file format error:
            # when the available bytes (after decompressing) do not
            # pack into exact rows.
            raise FormatError(
              'Wrong size for decompressed IDAT chunk.')
        assert len(a) == 0

    def validate_signature(self):
        """If signature (header) has not been read then read and
        validate it; otherwise do nothing.
        """

        if self.signature:
            return
        self.signature = self.file.read(8)
        if self.signature != _signature:
            raise FormatError("PNG file has invalid signature.")

    def preamble(self, lenient=False):
        """
        Extract the image metadata by reading the initial part of
        the PNG file up to the start of the ``IDAT`` chunk.  All the
        chunks that precede the ``IDAT`` chunk are read and either
        processed for metadata or discarded.

        If the optional `lenient` argument evaluates to `True`, checksum
        failures will raise warnings rather than exceptions.
        """

        self.validate_signature()

        while True:
            if not self.atchunk:
                self.atchunk = self.chunklentype()
                if self.atchunk is None:
                    raise FormatError(
                      'This PNG file has no IDAT chunks.')
            if self.atchunk[1] == b'IDAT':
                return
            self.process_chunk(lenient=lenient)

    def chunklentype(self):
        """Reads just enough of the input to determine the next
        chunk's length and type, returned as a (*length*, *type*) pair
        where *type* is a string.  If there are no more chunks, ``None``
        is returned.
        """

        x = self.file.read(8)
        if not x:
            return None
        if len(x) != 8:
            raise FormatError(
              'End of file whilst reading chunk length and type.')
        length,type = struct.unpack('!I4s', x)
        if length > 2**31-1:
            raise FormatError('Chunk %s is too large: %d.' % (type,length))
        return length,type

    def process_chunk(self, lenient=False):
        """Process the next chunk and its data.  This only processes the
        following chunk types, all others are ignored: ``IHDR``,
        ``PLTE``, ``bKGD``, ``tRNS``, ``gAMA``, ``sBIT``, ``pHYs``.

        If the optional `lenient` argument evaluates to `True`,
        checksum failures will raise warnings rather than exceptions.
        """

        type, data = self.chunk(lenient=lenient)
        method = '_process_' + as_str(type)
        m = getattr(self, method, None)
        if m:
            m(data)

    def _process_IHDR(self, data):
        # http://www.w3.org/TR/PNG/#11IHDR
        if len(data) != 13:
            raise FormatError('IHDR chunk has incorrect length.')
        (self.width, self.height, self.bitdepth, self.color_type,
         self.compression, self.filter,
         self.interlace) = struct.unpack("!2I5B", data)

        check_bitdepth_colortype(self.bitdepth, self.color_type)

        if self.compression != 0:
            raise Error("unknown compression method %d" % self.compression)
        if self.filter != 0:
            raise FormatError("Unknown filter method %d,"
              " see http://www.w3.org/TR/2003/REC-PNG-20031110/#9Filters ."
              % self.filter)
        if self.interlace not in (0,1):
            raise FormatError("Unknown interlace method %d,"
              " see http://www.w3.org/TR/2003/REC-PNG-20031110/#8InterlaceMethods ."
              % self.interlace)

        # Derived values
        # http://www.w3.org/TR/PNG/#6Colour-values
        colormap =  bool(self.color_type & 1)
        greyscale = not (self.color_type & 2)
        alpha = bool(self.color_type & 4)
        color_planes = (3,1)[greyscale or colormap]
        planes = color_planes + alpha

        self.colormap = colormap
        self.greyscale = greyscale
        self.alpha = alpha
        self.color_planes = color_planes
        self.planes = planes
        self.psize = float(self.bitdepth)/float(8) * planes
        if int(self.psize) == self.psize:
            self.psize = int(self.psize)
        self.row_bytes = int(math.ceil(self.width * self.psize))
        # Stores PLTE chunk if present, and is used to check
        # chunk ordering constraints.
        self.plte = None
        # Stores tRNS chunk if present, and is used to check chunk
        # ordering constraints.
        self.trns = None
        # Stores sbit chunk if present.
        self.sbit = None

    def _process_PLTE(self, data):
        # http://www.w3.org/TR/PNG/#11PLTE
        if self.plte:
            warnings.warn("Multiple PLTE chunks present.")
        self.plte = data
        if len(data) % 3 != 0:
            raise FormatError(
              "PLTE chunk's length should be a multiple of 3.")
        if len(data) > (2**self.bitdepth)*3:
            raise FormatError("PLTE chunk is too long.")
        if len(data) == 0:
            raise FormatError("Empty PLTE is not allowed.")

    def _process_bKGD(self, data):
        try:
            if self.colormap:
                if not self.plte:
                    warnings.warn(
                      "PLTE chunk is required before bKGD chunk.")
                self.background = struct.unpack('B', data)
            else:
                self.background = struct.unpack("!%dH" % self.color_planes,
                  data)
        except struct.error:
            raise FormatError("bKGD chunk has incorrect length.")

    def _process_tRNS(self, data):
        # http://www.w3.org/TR/PNG/#11tRNS
        self.trns = data
        if self.colormap:
            if not self.plte:
                warnings.warn("PLTE chunk is required before tRNS chunk.")
            else:
                if len(data) > len(self.plte)/3:
                    # Was warning, but promoted to Error as it
                    # would otherwise cause pain later on.
                    raise FormatError("tRNS chunk is too long.")
        else:
            if self.alpha:
                raise FormatError(
                  "tRNS chunk is not valid with colour type %d." %
                  self.color_type)
            try:
                self.transparent = \
                    struct.unpack("!%dH" % self.color_planes, data)
            except struct.error:
                raise FormatError("tRNS chunk has incorrect length.")

    def _process_gAMA(self, data):
        try:
            self.gamma = struct.unpack("!L", data)[0] / 100000.0
        except struct.error:
            raise FormatError("gAMA chunk has incorrect length.")

    def _process_sBIT(self, data):
        self.sbit = data
        if (self.colormap and len(data) != 3 or
            not self.colormap and len(data) != self.planes):
            raise FormatError("sBIT chunk has incorrect length.")

    def _process_pHYs(self, data):
        # http://www.w3.org/TR/PNG/#11pHYs
        self.phys = data
        fmt = "!LLB"
        if len(data) != struct.calcsize(fmt):
            raise FormatError("pHYs chunk has incorrect length.")
        self.x_pixels_per_unit, self.y_pixels_per_unit, unit = struct.unpack(fmt,data)
        self.unit_is_meter = bool(unit)

    def read(self, lenient=False):
        """
        Read the PNG file and decode it.  Returns (`width`, `height`,
        `pixels`, `metadata`).

        May use excessive memory.

        `pixels` are returned in boxed row flat pixel format.

        If the optional `lenient` argument evaluates to True,
        checksum failures will raise warnings rather than exceptions.
        """

        def iteridat():
            """Iterator that yields all the ``IDAT`` chunks as strings."""
            while True:
                try:
                    type, data = self.chunk(lenient=lenient)
                except ValueError as e:
                    raise ChunkError(e.args[0])
                if type == b'IEND':
                    # http://www.w3.org/TR/PNG/#11IEND
                    break
                if type != b'IDAT':
                    continue
                # type == b'IDAT'
                # http://www.w3.org/TR/PNG/#11IDAT
                if self.colormap and not self.plte:
                    warnings.warn("PLTE chunk is required before IDAT chunk")
                yield data

        def iterdecomp(idat):
            """Iterator that yields decompressed strings.  `idat` should
            be an iterator that yields the ``IDAT`` chunk data.
            """

            # Currently, with no max_length parameter to decompress,
            # this routine will do one yield per IDAT chunk: Not very
            # incremental.
            d = zlib.decompressobj()
            # Each IDAT chunk is passed to the decompressor, then any
            # remaining state is decompressed out.
            for data in idat:
                # :todo: add a max_length argument here to limit output
                # size.
                yield array('B', d.decompress(data))
            yield array('B', d.flush())

        self.preamble(lenient=lenient)
        raw = iterdecomp(iteridat())

        if self.interlace:
            raw = array('B', itertools.chain(*raw))
            arraycode = 'BH'[self.bitdepth>8]
            # Like :meth:`group` but producing an array.array object for
            # each row.
            pixels = map(lambda *row: array(arraycode, row),
                       *[iter(self.deinterlace(raw))]*self.width*self.planes)
        else:
            pixels = self.iterboxed(self.iterstraight(raw))
        meta = dict()
        for attr in 'greyscale alpha planes bitdepth interlace'.split():
            meta[attr] = getattr(self, attr)
        meta['size'] = (self.width, self.height)
        for attr in 'gamma transparent background'.split():
            a = getattr(self, attr, None)
            if a is not None:
                meta[attr] = a
        if self.plte:
            meta['palette'] = self.palette()
        return self.width, self.height, pixels, meta


    def read_flat(self):
        """
        Read a PNG file and decode it into flat row flat pixel format.
        Returns (*width*, *height*, *pixels*, *metadata*).

        May use excessive memory.

        `pixels` are returned in flat row flat pixel format.

        See also the :meth:`read` method which returns pixels in the
        more stream-friendly boxed row flat pixel format.
        """

        x, y, pixel, meta = self.read()
        arraycode = 'BH'[meta['bitdepth']>8]
        pixel = array(arraycode, itertools.chain(*pixel))
        return x, y, pixel, meta

    def palette(self, alpha='natural'):
        """Returns a palette that is a sequence of 3-tuples or 4-tuples,
        synthesizing it from the ``PLTE`` and ``tRNS`` chunks.  These
        chunks should have already been processed (for example, by
        calling the :meth:`preamble` method).  All the tuples are the
        same size: 3-tuples if there is no ``tRNS`` chunk, 4-tuples when
        there is a ``tRNS`` chunk.  Assumes that the image is colour type
        3 and therefore a ``PLTE`` chunk is required.

        If the `alpha` argument is ``'force'`` then an alpha channel is
        always added, forcing the result to be a sequence of 4-tuples.
        """

        if not self.plte:
            raise FormatError(
                "Required PLTE chunk is missing in colour type 3 image.")
        plte = group(array('B', self.plte), 3)
        if self.trns or alpha == 'force':
            trns = array('B', self.trns or '')
            trns.extend([255]*(len(plte)-len(trns)))
            plte = list(map(operator.add, plte, group(trns, 1)))
        return plte

    def asDirect(self):
        """Returns the image data as a direct representation of an
        ``x * y * planes`` array.  This method is intended to remove the
        need for callers to deal with palettes and transparency
        themselves.  Images with a palette (colour type 3)
        are converted to RGB or RGBA; images with transparency (a
        ``tRNS`` chunk) are converted to LA or RGBA as appropriate.
        When returned in this format the pixel values represent the
        colour value directly without needing to refer to palettes or
        transparency information.

        Like the :meth:`read` method this method returns a 4-tuple:

        (*width*, *height*, *pixels*, *meta*)

        This method normally returns pixel values with the bit depth
        they have in the source image, but when the source PNG has an
        ``sBIT`` chunk it is inspected and can reduce the bit depth of
        the result pixels; pixel values will be reduced according to
        the bit depth specified in the ``sBIT`` chunk (PNG nerds should
        note a single result bit depth is used for all channels; the
        maximum of the ones specified in the ``sBIT`` chunk.  An RGB565
        image will be rescaled to 6-bit RGB666).

        The *meta* dictionary that is returned reflects the `direct`
        format and not the original source image.  For example, an RGB
        source image with a ``tRNS`` chunk to represent a transparent
        colour, will have ``planes=3`` and ``alpha=False`` for the
        source image, but the *meta* dictionary returned by this method
        will have ``planes=4`` and ``alpha=True`` because an alpha
        channel is synthesized and added.

        *pixels* is the pixel data in boxed row flat pixel format (just
        like the :meth:`read` method).

        All the other aspects of the image data are not changed.
        """

        self.preamble()

        # Simple case, no conversion necessary.
        if not self.colormap and not self.trns and not self.sbit:
            return self.read()

        x,y,pixels,meta = self.read()

        if self.colormap:
            meta['colormap'] = False
            meta['alpha'] = bool(self.trns)
            meta['bitdepth'] = 8
            meta['planes'] = 3 + bool(self.trns)
            plte = self.palette()
            def iterpal(pixels):
                for row in pixels:
                    row = [plte[x] for x in row]
                    yield array('B', itertools.chain(*row))
            pixels = iterpal(pixels)
        elif self.trns:
            # It would be nice if there was some reasonable way
            # of doing this without generating a whole load of
            # intermediate tuples.  But tuples does seem like the
            # easiest way, with no other way clearly much simpler or
            # much faster.  (Actually, the L to LA conversion could
            # perhaps go faster (all those 1-tuples!), but I still
            # wonder whether the code proliferation is worth it)
            it = self.transparent
            maxval = 2**meta['bitdepth']-1
            planes = meta['planes']
            meta['alpha'] = True
            meta['planes'] += 1
            typecode = 'BH'[meta['bitdepth']>8]
            def itertrns(pixels):
                for row in pixels:
                    # For each row we group it into pixels, then form a
                    # characterisation vector that says whether each
                    # pixel is opaque or not.  Then we convert
                    # True/False to 0/maxval (by multiplication),
                    # and add it as the extra channel.
                    row = group(row, planes)
                    opa = map(it.__ne__, row)
                    opa = map(maxval.__mul__, opa)
                    opa = list(zip(opa)) # convert to 1-tuples
                    yield array(typecode,
                      itertools.chain(*map(operator.add, row, opa)))
            pixels = itertrns(pixels)
        targetbitdepth = None
        if self.sbit:
            sbit = struct.unpack('%dB' % len(self.sbit), self.sbit)
            targetbitdepth = max(sbit)
            if targetbitdepth > meta['bitdepth']:
                raise Error('sBIT chunk %r exceeds bitdepth %d' %
                    (sbit,self.bitdepth))
            if min(sbit) <= 0:
                raise Error('sBIT chunk %r has a 0-entry' % sbit)
            if targetbitdepth == meta['bitdepth']:
                targetbitdepth = None
        if targetbitdepth:
            shift = meta['bitdepth'] - targetbitdepth
            meta['bitdepth'] = targetbitdepth
            def itershift(pixels):
                for row in pixels:
                    yield [p >> shift for p in row]
            pixels = itershift(pixels)
        return x,y,pixels,meta

    def asFloat(self, maxval=1.0):
        """Return image pixels as per :meth:`asDirect` method, but scale
        all pixel values to be floating point values between 0.0 and
        *maxval*.
        """

        x,y,pixels,info = self.asDirect()
        sourcemaxval = 2**info['bitdepth']-1
        del info['bitdepth']
        info['maxval'] = float(maxval)
        factor = float(maxval)/float(sourcemaxval)
        def iterfloat():
            for row in pixels:
                yield [factor * p for p in row]
        return x,y,iterfloat(),info

    def _as_rescale(self, get, targetbitdepth):
        """Helper used by :meth:`asRGB8` and :meth:`asRGBA8`."""

        width,height,pixels,meta = get()
        maxval = 2**meta['bitdepth'] - 1
        targetmaxval = 2**targetbitdepth - 1
        factor = float(targetmaxval) / float(maxval)
        meta['bitdepth'] = targetbitdepth
        def iterscale():
            for row in pixels:
                yield [int(round(x*factor)) for x in row]
        if maxval == targetmaxval:
            return width, height, pixels, meta
        else:
            return width, height, iterscale(), meta

    def asRGB8(self):
        """Return the image data as an RGB pixels with 8-bits per
        sample.  This is like the :meth:`asRGB` method except that
        this method additionally rescales the values so that they
        are all between 0 and 255 (8-bit).  In the case where the
        source image has a bit depth < 8 the transformation preserves
        all the information; where the source image has bit depth
        > 8, then rescaling to 8-bit values loses precision.  No
        dithering is performed.  Like :meth:`asRGB`, an alpha channel
        in the source image will raise an exception.

        This function returns a 4-tuple:
        (*width*, *height*, *pixels*, *metadata*).
        *width*, *height*, *metadata* are as per the
        :meth:`read` method.
        
        *pixels* is the pixel data in boxed row flat pixel format.
        """

        return self._as_rescale(self.asRGB, 8)

    def asRGBA8(self):
        """Return the image data as RGBA pixels with 8-bits per
        sample.  This method is similar to :meth:`asRGB8` and
        :meth:`asRGBA`:  The result pixels have an alpha channel, *and*
        values are rescaled to the range 0 to 255.  The alpha channel is
        synthesized if necessary (with a small speed penalty).
        """

        return self._as_rescale(self.asRGBA, 8)

    def asRGB(self):
        """Return image as RGB pixels.  RGB colour images are passed
        through unchanged; greyscales are expanded into RGB
        triplets (there is a small speed overhead for doing this).

        An alpha channel in the source image will raise an
        exception.

        The return values are as for the :meth:`read` method
        except that the *metadata* reflect the returned pixels, not the
        source image.  In particular, for this method
        ``metadata['greyscale']`` will be ``False``.
        """

        width,height,pixels,meta = self.asDirect()
        if meta['alpha']:
            raise Error("will not convert image with alpha channel to RGB")
        if not meta['greyscale']:
            return width,height,pixels,meta
        meta['greyscale'] = False
        typecode = 'BH'[meta['bitdepth'] > 8]
        def iterrgb():
            for row in pixels:
                a = array(typecode, [0]) * 3 * width
                for i in range(3):
                    a[i::3] = row
                yield a
        return width,height,iterrgb(),meta

    def asRGBA(self):
        """Return image as RGBA pixels.  Greyscales are expanded into
        RGB triplets; an alpha channel is synthesized if necessary.
        The return values are as for the :meth:`read` method
        except that the *metadata* reflect the returned pixels, not the
        source image.  In particular, for this method
        ``metadata['greyscale']`` will be ``False``, and
        ``metadata['alpha']`` will be ``True``.
        """

        width,height,pixels,meta = self.asDirect()
        if meta['alpha'] and not meta['greyscale']:
            return width,height,pixels,meta
        typecode = 'BH'[meta['bitdepth'] > 8]
        maxval = 2**meta['bitdepth'] - 1
        maxbuffer = struct.pack('=' + typecode, maxval) * 4 * width
        def newarray():
            return array(typecode, maxbuffer)

        if meta['alpha'] and meta['greyscale']:
            # LA to RGBA
            def convert():
                for row in pixels:
                    # Create a fresh target row, then copy L channel
                    # into first three target channels, and A channel
                    # into fourth channel.
                    a = newarray()
                    pngfilters.convert_la_to_rgba(row, a)
                    yield a
        elif meta['greyscale']:
            # L to RGBA
            def convert():
                for row in pixels:
                    a = newarray()
                    pngfilters.convert_l_to_rgba(row, a)
                    yield a
        else:
            assert not meta['alpha'] and not meta['greyscale']
            # RGB to RGBA
            def convert():
                for row in pixels:
                    a = newarray()
                    pngfilters.convert_rgb_to_rgba(row, a)
                    yield a
        meta['alpha'] = True
        meta['greyscale'] = False
        return width,height,convert(),meta

def check_bitdepth_colortype(bitdepth, colortype):
    """Check that `bitdepth` and `colortype` are both valid,
    and specified in a valid combination. Returns if valid,
    raise an Exception if not valid.
    """

    if bitdepth not in (1,2,4,8,16):
        raise FormatError("invalid bit depth %d" % bitdepth)
    if colortype not in (0,2,3,4,6):
        raise FormatError("invalid colour type %d" % colortype)
    # Check indexed (palettized) images have 8 or fewer bits
    # per pixel; check only indexed or greyscale images have
    # fewer than 8 bits per pixel.
    if colortype & 1 and bitdepth > 8:
        raise FormatError(
          "Indexed images (colour type %d) cannot"
          " have bitdepth > 8 (bit depth %d)."
          " See http://www.w3.org/TR/2003/REC-PNG-20031110/#table111 ."
          % (bitdepth, colortype))
    if bitdepth < 8 and colortype not in (0,3):
        raise FormatError("Illegal combination of bit depth (%d)"
          " and colour type (%d)."
          " See http://www.w3.org/TR/2003/REC-PNG-20031110/#table111 ."
          % (bitdepth, colortype))

def isinteger(x):
    try:
        return int(x) == x
    except (TypeError, ValueError):
        return False


# === Support for users without Cython ===

try:
    pngfilters
except NameError:
    class pngfilters(object):
        def undo_filter_sub(filter_unit, scanline, previous, result):
            """Undo sub filter."""

            ai = 0
            # Loops starts at index fu.  Observe that the initial part
            # of the result is already filled in correctly with
            # scanline.
            for i in range(filter_unit, len(result)):
                x = scanline[i]
                a = result[ai]
                result[i] = (x + a) & 0xff
                ai += 1
        undo_filter_sub = staticmethod(undo_filter_sub)

        def undo_filter_up(filter_unit, scanline, previous, result):
            """Undo up filter."""

            for i in range(len(result)):
                x = scanline[i]
                b = previous[i]
                result[i] = (x + b) & 0xff
        undo_filter_up = staticmethod(undo_filter_up)

        def undo_filter_average(filter_unit, scanline, previous, result):
            """Undo up filter."""

            ai = -filter_unit
            for i in range(len(result)):
                x = scanline[i]
                if ai < 0:
                    a = 0
                else:
                    a = result[ai]
                b = previous[i]
                result[i] = (x + ((a + b) >> 1)) & 0xff
                ai += 1
        undo_filter_average = staticmethod(undo_filter_average)

        def undo_filter_paeth(filter_unit, scanline, previous, result):
            """Undo Paeth filter."""

            # Also used for ci.
            ai = -filter_unit
            for i in range(len(result)):
                x = scanline[i]
                if ai < 0:
                    a = c = 0
                else:
                    a = result[ai]
                    c = previous[ai]
                b = previous[i]
                p = a + b - c
                pa = abs(p - a)
                pb = abs(p - b)
                pc = abs(p - c)
                if pa <= pb and pa <= pc:
                    pr = a
                elif pb <= pc:
                    pr = b
                else:
                    pr = c
                result[i] = (x + pr) & 0xff
                ai += 1
        undo_filter_paeth = staticmethod(undo_filter_paeth)

        def convert_la_to_rgba(row, result):
            for i in range(3):
                result[i::4] = row[0::2]
            result[3::4] = row[1::2]
        convert_la_to_rgba = staticmethod(convert_la_to_rgba)

        def convert_l_to_rgba(row, result):
            """Convert a grayscale image to RGBA. This method assumes
            the alpha channel in result is already correctly
            initialized.
            """
            for i in range(3):
                result[i::4] = row
        convert_l_to_rgba = staticmethod(convert_l_to_rgba)

        def convert_rgb_to_rgba(row, result):
            """Convert an RGB image to RGBA. This method assumes the
            alpha channel in result is already correctly initialized.
            """
            for i in range(3):
                result[i::4] = row[i::3]
        convert_rgb_to_rgba = staticmethod(convert_rgb_to_rgba)


# === Command Line Support ===

def read_pam_header(infile):
    """
    Read (the rest of a) PAM header.  `infile` should be positioned
    immediately after the initial 'P7' line (at the beginning of the
    second line).  Returns are as for `read_pnm_header`.
    """
    
    # Unlike PBM, PGM, and PPM, we can read the header a line at a time.
    header = dict()
    while True:
        l = infile.readline().strip()
        if l == b'ENDHDR':
            break
        if not l:
            raise EOFError('PAM ended prematurely')
        if l[0] == b'#':
            continue
        l = l.split(None, 1)
        if l[0] not in header:
            header[l[0]] = l[1]
        else:
            header[l[0]] += b' ' + l[1]

    required = [b'WIDTH', b'HEIGHT', b'DEPTH', b'MAXVAL']
    WIDTH,HEIGHT,DEPTH,MAXVAL = required
    present = [x for x in required if x in header]
    if len(present) != len(required):
        raise Error('PAM file must specify WIDTH, HEIGHT, DEPTH, and MAXVAL')
    width = int(header[WIDTH])
    height = int(header[HEIGHT])
    depth = int(header[DEPTH])
    maxval = int(header[MAXVAL])
    if (width <= 0 or
        height <= 0 or
        depth <= 0 or
        maxval <= 0):
        raise Error(
          'WIDTH, HEIGHT, DEPTH, MAXVAL must all be positive integers')
    return 'P7', width, height, depth, maxval

def read_pnm_header(infile, supported=(b'P5', b'P6')):
    """
    Read a PNM header, returning (format,width,height,depth,maxval).
    `width` and `height` are in pixels.  `depth` is the number of
    channels in the image; for PBM and PGM it is synthesized as 1, for
    PPM as 3; for PAM images it is read from the header.  `maxval` is
    synthesized (as 1) for PBM images.
    """

    # Generally, see http://netpbm.sourceforge.net/doc/ppm.html
    # and http://netpbm.sourceforge.net/doc/pam.html

    # Technically 'P7' must be followed by a newline, so by using
    # rstrip() we are being liberal in what we accept.  I think this
    # is acceptable.
    type = infile.read(3).rstrip()
    if type not in supported:
        raise NotImplementedError('file format %s not supported' % type)
    if type == b'P7':
        # PAM header parsing is completely different.
        return read_pam_header(infile)
    # Expected number of tokens in header (3 for P4, 4 for P6)
    expected = 4
    pbm = (b'P1', b'P4')
    if type in pbm:
        expected = 3
    header = [type]

    # We have to read the rest of the header byte by byte because the
    # final whitespace character (immediately following the MAXVAL in
    # the case of P6) may not be a newline.  Of course all PNM files in
    # the wild use a newline at this point, so it's tempting to use
    # readline; but it would be wrong.
    def getc():
        c = infile.read(1)
        if not c:
            raise Error('premature EOF reading PNM header')
        return c

    c = getc()
    while True:
        # Skip whitespace that precedes a token.
        while c.isspace():
            c = getc()
        # Skip comments.
        while c == '#':
            while c not in b'\n\r':
                c = getc()
        if not c.isdigit():
            raise Error('unexpected character %s found in header' % c)
        # According to the specification it is legal to have comments
        # that appear in the middle of a token.
        # This is bonkers; I've never seen it; and it's a bit awkward to
        # code good lexers in Python (no goto).  So we break on such
        # cases.
        token = b''
        while c.isdigit():
            token += c
            c = getc()
        # Slight hack.  All "tokens" are decimal integers, so convert
        # them here.
        header.append(int(token))
        if len(header) == expected:
            break
    # Skip comments (again)
    while c == '#':
        while c not in '\n\r':
            c = getc()
    if not c.isspace():
        raise Error('expected header to end with whitespace, not %s' % c)

    if type in pbm:
        # synthesize a MAXVAL
        header.append(1)
    depth = (1,3)[type == b'P6']
    return header[0], header[1], header[2], depth, header[3]

def write_pnm(file, width, height, pixels, meta):
    """Write a Netpbm PNM/PAM file.
    """

    bitdepth = meta['bitdepth']
    maxval = 2**bitdepth - 1
    # Rudely, the number of image planes can be used to determine
    # whether we are L (PGM), LA (PAM), RGB (PPM), or RGBA (PAM).
    planes = meta['planes']
    # Can be an assert as long as we assume that pixels and meta came
    # from a PNG file.
    assert planes in (1,2,3,4)
    if planes in (1,3):
        if 1 == planes:
            # PGM
            # Could generate PBM if maxval is 1, but we don't (for one
            # thing, we'd have to convert the data, not just blat it
            # out).
            fmt = 'P5'
        else:
            # PPM
            fmt = 'P6'
        header = '%s %d %d %d\n' % (fmt, width, height, maxval)
    if planes in (2,4):
        # PAM
        # See http://netpbm.sourceforge.net/doc/pam.html
        if 2 == planes:
            tupltype = 'GRAYSCALE_ALPHA'
        else:
            tupltype = 'RGB_ALPHA'
        header = ('P7\nWIDTH %d\nHEIGHT %d\nDEPTH %d\nMAXVAL %d\n'
                  'TUPLTYPE %s\nENDHDR\n' %
                  (width, height, planes, maxval, tupltype))
    file.write(header.encode('ascii'))
    # Values per row
    vpr = planes * width
    # struct format
    fmt = '>%d' % vpr
    if maxval > 0xff:
        fmt = fmt + 'H'
    else:
        fmt = fmt + 'B'
    for row in pixels:
        file.write(struct.pack(fmt, *row))
    file.flush()

def color_triple(color):
    """
    Convert a command line colour value to a RGB triple of integers.
    FIXME: Somewhere we need support for greyscale backgrounds etc.
    """
    if color.startswith('#') and len(color) == 4:
        return (int(color[1], 16),
                int(color[2], 16),
                int(color[3], 16))
    if color.startswith('#') and len(color) == 7:
        return (int(color[1:3], 16),
                int(color[3:5], 16),
                int(color[5:7], 16))
    elif color.startswith('#') and len(color) == 13:
        return (int(color[1:5], 16),
                int(color[5:9], 16),
                int(color[9:13], 16))

def _add_common_options(parser):
    """Call *parser.add_option* for each of the options that are
    common between this PNG--PNM conversion tool and the gen
    tool.
    """
    parser.add_option("-i", "--interlace",
                      default=False, action="store_true",
                      help="create an interlaced PNG file (Adam7)")
    parser.add_option("-t", "--transparent",
                      action="store", type="string", metavar="#RRGGBB",
                      help="mark the specified colour as transparent")
    parser.add_option("-b", "--background",
                      action="store", type="string", metavar="#RRGGBB",
                      help="save the specified background colour")
    parser.add_option("-g", "--gamma",
                      action="store", type="float", metavar="value",
                      help="save the specified gamma value")
    parser.add_option("-c", "--compression",
                      action="store", type="int", metavar="level",
                      help="zlib compression level (0-9)")
    return parser

def _main(argv):
    """
    Run the PNG encoder with options from the command line.
    """

    # Parse command line arguments
    from optparse import OptionParser
    version = '%prog ' + __version__
    parser = OptionParser(version=version)
    parser.set_usage("%prog [options] [imagefile]")
    parser.add_option('-r', '--read-png', default=False,
                      action='store_true',
                      help='Read PNG, write PNM')
    parser.add_option("-a", "--alpha",
                      action="store", type="string", metavar="pgmfile",
                      help="alpha channel transparency (RGBA)")
    _add_common_options(parser)

    (options, args) = parser.parse_args(args=argv[1:])

    # Convert options
    if options.transparent is not None:
        options.transparent = color_triple(options.transparent)
    if options.background is not None:
        options.background = color_triple(options.background)

    # Prepare input and output files
    if len(args) == 0:
        infilename = '-'
        infile = sys.stdin
    elif len(args) == 1:
        infilename = args[0]
        infile = open(infilename, 'rb')
    else:
        parser.error("more than one input file")
    outfile = sys.stdout
    if sys.platform == "win32":
        import msvcrt, os
        msvcrt.setmode(sys.stdout.fileno(), os.O_BINARY)

    if options.read_png:
        # Encode PNG to PPM
        png = Reader(file=infile)
        width,height,pixels,meta = png.asDirect()
        write_pnm(outfile, width, height, pixels, meta) 
    else:
        # Encode PNM to PNG
        format, width, height, depth, maxval = \
          read_pnm_header(infile, (b'P5',b'P6',b'P7'))
        # When it comes to the variety of input formats, we do something
        # rather rude.  Observe that L, LA, RGB, RGBA are the 4 colour
        # types supported by PNG and that they correspond to 1, 2, 3, 4
        # channels respectively.  So we use the number of channels in
        # the source image to determine which one we have.  We do not
        # care about TUPLTYPE.
        greyscale = depth <= 2
        pamalpha = depth in (2,4)
        supported = [2**x-1 for x in range(1,17)]
        try:
            mi = supported.index(maxval)
        except ValueError:
            raise NotImplementedError(
              'your maxval (%s) not in supported list %s' %
              (maxval, str(supported)))
        bitdepth = mi+1
        writer = Writer(width, height,
                        greyscale=greyscale,
                        bitdepth=bitdepth,
                        interlace=options.interlace,
                        transparent=options.transparent,
                        background=options.background,
                        alpha=bool(pamalpha or options.alpha),
                        gamma=options.gamma,
                        compression=options.compression)
        if options.alpha:
            pgmfile = open(options.alpha, 'rb')
            format, awidth, aheight, adepth, amaxval = \
              read_pnm_header(pgmfile, 'P5')
            if amaxval != '255':
                raise NotImplementedError(
                  'maxval %s not supported for alpha channel' % amaxval)
            if (awidth, aheight) != (width, height):
                raise ValueError("alpha channel image size mismatch"
                                 " (%s has %sx%s but %s has %sx%s)"
                                 % (infilename, width, height,
                                    options.alpha, awidth, aheight))
            writer.convert_ppm_and_pgm(infile, pgmfile, outfile)
        else:
            writer.convert_pnm(infile, outfile)


if __name__ == '__main__':
    try:
        _main(sys.argv)
    except Error as e:
        print(e, file=sys.stderr)
```
