## Supplemental Software for "A Customizable Low-Cost System for Massively Parallel Zebrafish Behavior Phenotyping": EventSection.py

```
#!/usr/bin/python
import datetime
import time
import Fish
import fileloading # for the constant values and access to options
from datetime import timedelta

class EventSection:

	def __init__(self, namestring, startdate):
		self.starttime = datetime.datetime.strptime(startdate + " " + namestring.split('=')[1].split('-')[0].split('_')[1], "%m/%d/%Y %H:%M:%S") + datetime.timedelta(days=int(namestring.split('=')[1].split('-')[0].split('_')[0])) # datetime #df_timestart = datetime.datetime.strptime(startdate + " 13:00:15", "%Y-%m-%d %H:%M:%S") #lf_timestart = lf_timestart + datetime.timedelta(days=1)
		self.endtime = datetime.datetime.strptime(startdate + " " + namestring.split('=')[1].split('-')[1].split('_')[1], "%m/%d/%Y %H:%M:%S") + datetime.timedelta(days=int(namestring.split('=')[1].split('-')[1].split('_')[0])) # datetime #df_timestart = datetime.datetime.strptime(startdate + " 13:00:15", "%Y-%m-%d %H:%M:%S") #lf_timestart = lf_timestart + datetime.timedelta(days=1)
		self.name = namestring.split('=')[0].split('_')[1] # string #habituation_daytaps=1_15:25:30-1_16:59:00, e.g. daytaps
		self.type = namestring.split('=')[0].split('_')[0] # string #habituation_daytaps=1_15:25:30-1_16:59:00, e.g. "habituation"
		self.events = {}
		self.indexdict = {}

	def check_endtime(self, endDateTime):
		if self.starttime > endDateTime:
			print("Error, the sections file starttime is after the end of the run . . . this seems unlikely to ever happen")
		if self.endtime > endDateTime:
			print("Error, the end time listed in the sections file is after the real end of the file, replacing the end time - 1 min, but check your sections file")
			self.endtime = endDateTime - datetime.timedelta(minutes = 1)
		# Have gotten strange bugs in last second that don't seem to be an issue with first second, so not worrying about it for start time check
		if self.endtime == endDateTime:
			print("Warning, the end time listed in the sections file is the same as the real end of the file, which is risky because that final second is often incomplete, replacing the end time - 1 min")
			self.endtime = endDateTime - datetime.timedelta(minutes = 1)

	def check_starttime(self, startDateTime):
		if self.starttime < startDateTime:
			print("Error, the start time listed in the sections file is before the real beginning of the file, replacing the start time + 1 min, but check your sections file because it coud indicate a big problem")
			self.starttime = startDateTime + datetime.timedelta(minutes = 1)
	
	def add_indices(self, k, v1, v2): # binned time: (indexstart, indexend)
		self.indexdict[str(k)] = (v1,v2)

	#4:26:00 PM      1       v120 D40 a0.01 f500 d5 p D500 a1 f500 d5 p D455 v200;        
	def process_teensy_str(self, teensystr):
		annotations = []
		annotationsfinal = []
		teensy = teensystr.split(" ")
		# get all indices that start with D
		indices = [i for i, x in enumerate(teensy) if x.startswith("D")]
		#print(indices)
		for t in range(0, len(teensy)):
			if teensy[t].startswith('a') or teensy[t].startswith('v'):
				annotate = teensy[t]
				if len(indices) != 0:
					for i2 in range(0,len(indices)):
						if t > indices[i2]:
							annotate = annotate + "%" + teensy[indices[i2]]
				annotations.append(annotate)
					
			else:
				continue
		for a in range(0,len(annotations)):
			Dlist = annotations[a].split("%")
			totalD = 0
			for d in range(1, len(Dlist)):
				totalD = totalD + int(Dlist[d][1:])
			# If it's a signal at the very end of the string, such as light going off
			if totalD > 980:
				continue
			# NEED THE START AND END FRAMES HERE TO BE THE WINDOW FOR FINDING THE START OF MOVEMENT, NOT STOPPING BY THE END (CAN FINISH CURRENT MOTION)
			# Frames of a strong tap are 4-26 ideally
			# Initiates around frame 6-7 for strong non-habituated tap and frames 8-10 for slower
			# So call initiation 4-16 (could maybe get away with smaller window even for starting) and stopping point of 30
			# For strong dark flash, rare starts at 27 (some very rare even as early as 20-25!!) and most start in the 30s
			# Rule will be that fish can't have moved in the previous 10 frames to be counted for next event or for any event
			# For MSI with the 50 msec delay, the main tap response is around frame 24, which is a bit risky for those early dark flashes
			# 50 / 3.5 is 14, so 14 + 9-10 basically. We need to use the same windows whether MSI or plain event inside MSI block . . .
			# with MSI, the window could be 4-12, which transfers over to 18-26 starting, so we are overlapping. With 40 msec delay instead of 50 we are at 16-24 . .  (not a ton better)
			# plan, make dark flahses with v0 have earlier start and have the ones with v120 and v140 start at 26
			# need to divide totalD so that it's in frames, not in milliseconds
			if annotations[a][0] == 'a':
				startframe = int(totalD / fileloading.msecperframe) + 4			
				endframe = startframe + 8
			elif annotations[a][0] == 'v':
				if 100 < int(str(annotations[a][1:]).strip(";")) < 200: # NEED TO CHANGE THIS FOR ALL THE Vs SPECIFICALLY BETWEEN 100 and 200, OTHERWISE THEY WILL GO WITH THE ELSE STATEMENT (LIGHT AND OTHER DARKS)
					startframe = int(totalD / fileloading.msecperframe) + 26
					endframe = 266
				else:
					startframe = int(totalD / fileloading.msecperframe) + 18
					endframe = 266
			else:
				print("Error, the events are not acoustic or visual (a or v), or there is a bug")
			annotationsfinal.append(annotations[a].split("%")[0] + "%" + str(startframe) + "%" + str(endframe))
		#print(annotationsfinal)
		return annotationsfinal

	def add_event(self, eventtype, eventvoltage, eventtime):
		# This assumes the sections file just has slow-speed data labeled as "time" sections and high-speed data with various labels
		if self.type != "time":
			stringinput = str(''.join(eventvoltage.split("."))).strip()
			annotatedeventlist = self.process_teensy_str(stringinput)
			for annotatedevent in annotatedeventlist:
				if eventtype + "_" + str(''.join(  str(''.join(eventvoltage.split("."))).split(" ") )).strip().strip(";") + "_" + annotatedevent in self.events.keys():
					self.events[eventtype + "_" + str(''.join(  str(''.join(eventvoltage.split("."))).split(" ") )).strip().strip(";") + "_" + annotatedevent].append(eventtime)
				else:
					self.events[eventtype + "_" + str(''.join(  str(''.join(eventvoltage.split("."))).split(" ") )).strip().strip(";") + "_" + annotatedevent] = [eventtime]
```
