## Supplemental Software for "A Customizable Low-Cost System for Massively Parallel Zebrafish Behavior Phenotyping": fileloading.py

```
#!/usr/bin/python

import os,sys,glob,re,argparse
import numpy as np
import scipy
import datetime
import time
from datetime import timedelta
import os.path
from os import path
from Fish import Fish # fish object
from EventSection import EventSection # event section object


# Input arguments
parser = argparse.ArgumentParser(description='loading for fish behavior files')
parser.add_argument('-graphonly', action="store_true", dest="graphonly", default=False)
parser.add_argument('-j', type=str, action="store", dest="graphparameters", default="PlotParameters")
parser.add_argument('-t', type=str, action="store", dest="tstampfile")
parser.add_argument('-e', type=str, action="store", dest="eventsfile")
parser.add_argument('-c', type=str, action="store", dest="centroidfile")
parser.add_argument('-d', type=str, action="store", dest="dpixfile")
parser.add_argument('-m', type=str, action="store", dest="movieprefix")
parser.add_argument('-g', type=str, action="store", dest="genotypefile")
parser.add_argument('-s', type=str, action="store", dest="sectionsfile", default="sectionsfile")
parser.add_argument('-n', type=int, action="store", dest="numberofwells", default=96)
parser.add_argument('-i', type=float, action="store", dest="msecperframe", default=3.508772)
parser.add_argument('-v', type=str, action="store", dest="thresholdvalues", default="0.5,3.0") # List of thresholds for the distance and dpix calculations, which typically should not be touched. First is for distance, which is less robust than dpix, and after is the dpix value. This is the threshold value to be counted for a bout if greater than or equal to.
parser.add_argument('-w', type=str, action="store", dest="hsthresholdvalues", default="0.9,3.0") # List of high-speed thresholds for the distance and dpix calculations, which typically should not be touched. First is for distance, which is less robust than dpix, and after is the dpix value. This is the threshold value to be counted for a bout if greater than or equal to.
parser.add_argument('-f', type=str, action="store", dest="thresholdframes", default="1,3") # Same as above but for number of frames.
parser.add_argument('-x', type=str, action="store", dest="hsthresholdframes", default="2,3") # Same as above but for number of frames and high-speed data.
# The classic Schier (and now Prober) sleep plots are inactive min / hour (sleep) and active sec / hour (with sleep bouts not counted)
parser.add_argument('-a', type=str, action="store", dest="activitytimes", default="1/60,60/600,60/3600,1/3600") # list of comparisons for activity data in seconds
parser.add_argument('-y', type=str, action="store", dest="activitytimesthresholds", default="1,10") # thresholds for activity data, first distance and second dpix (differs from bout thresholds because we don't have frame considerations)
parser.add_argument('-b', type=str, action="store", dest="boutbins", default="60,600,3600") # list of times bins for the bout data (ie, ave bout speed / minute) in seconds
parser.add_argument('-z', type=str, action="store", dest="seizurefilters", default="4.0,300,1300,70") # ((boutrev > 4.0) and (300 < (boutspeed) < 1300) and (boutdistance > 70)):
parser.add_argument('-l', type=int, action="store", dest="lightbaseline", default=200) # baseline level of light, used to determine what is a dark flash for filtering O-bend
parser.add_argument('-o', type=str, action="store", dest="obendfilter", default="60,10") # two measures that are intersected (both must be true), and responses with greater magnitude than both are considered true O-bends. The two measures are "responsetime" and "sumabsha" (sum of absolute value of heading angles). To change the type of measure, code in graphsstatsandfilter.py must be changed
parser.add_argument('-p', type=str, action="store", dest="cbendfilter", default="0.2,1500") # two measures that are intersected (both must be true), and responses with greater magnitude than both are considered true C-bends. The two measures are "responsevelocity" and "responsecumdpix". To change the type of measure, code in graphsstatsandfilter.py must be changed

args = parser.parse_args()
graphonly = args.graphonly
graphparameters = args.graphparameters
if(not graphonly):
	tstampfile = args.tstampfile
	eventsfile = args.eventsfile
	centroidfile = args.centroidfile
	dpixfile = args.dpixfile
	movieprefix = args.movieprefix
	genotypefile = args.genotypefile
	sectionsfile = args.sectionsfile
thresholdvalues = list(map(float, args.thresholdvalues.split(',')))
hsthresholdvalues = list(map(float, args.hsthresholdvalues.split(',')))
thresholdframes = list(map(int, args.thresholdframes.split(',')))
hsthresholdframes = list(map(int, args.hsthresholdframes.split(',')))
numberofwells = args.numberofwells
msecperframe = args.msecperframe
activitytimes = list(map(int, re.split(',|, |/|/', args.activitytimes)))
activitytimesthresholds = list(map(float, args.activitytimesthresholds.split(',')))
boutbins = list(map(int, args.boutbins.split(','))) # these bins and activity bins are not going to be less than a second (that doesn't work in code well), so it's fine to use int instead of float
seizurefilters = list(map(float, args.seizurefilters.split(',')))
lightbaseline = args.lightbaseline
obendfilter = list(map(float, args.obendfilter.split(',')))
cbendfilter = list(map(float, args.cbendfilter.split(',')))

# Helper functions, cart2pol and faststrptime

# Convert cartesian coordinates to polar
def cart2pol(x, y):
	rho = np.sqrt(x**2 + y**2)
	theta = np.arctan2(y, x)
	return(rho, theta)


# A fast function for reading time data
# This depends on have the below format as output, which it is from our LabView program
def faststrptime(val):
	#Example
	#6/18/201612:59:34 PM
	splits1 = val.split("/")
	splits2 = splits1[2].split(":")
	return datetime.datetime(
		int(splits1[2][0:4]), # %Year
		int(splits1[0]), # %Month
		int(splits1[1]), # %Day
		int(splits2[0][4:len(splits2[0])]), # %Hour
		int(splits2[1]), # %Minute
		int(splits2[2][0:2]), # %Second
	)

# End Helper functions


# Loading of the centroid position data for the high-speed movies
def load_movie_motion_pos(hs_pos, thistime, counter):
	# At this point I actually don't care if the data is centered
	# All I want it for is distances
	moviename = movieprefix + str(counter) + ".avi.centroid2"
	cenfile = open(moviename, 'r')
	hscen_data_list = []
	lines = cenfile.readlines()
	for line in lines:
		hscen_data_list.append(int(line))
	hscen_data_array = np.array(hscen_data_list)
	hscen_data_array = hscen_data_array.reshape(hscen_data_array.size // (numberofwells*2), (numberofwells*2))
	hs_pos[thistime] = hscen_data_array


# Loading of the delta pixel motion data for the high-speed movies
def load_movie_motion(hs_dpix, thistime, counter):
	moviename = movieprefix + str(counter) + ".avi.motion2"
	if not path.exists(moviename):
		print("Not all high-speed movies are tracked. Please check. Exiting.")
		exit()
	firstdpix = open(moviename, 'r')
	hsdp_data_list = []
	lines = firstdpix.readlines()
	for line in lines:
		hsdp_data_list.append(int(line))
	hsdp_data_array = np.array(hsdp_data_list)
	hsdp_data_array = hsdp_data_array.reshape(hsdp_data_array.size // numberofwells, numberofwells)
	hs_dpix[thistime] = hsdp_data_array


# Loading in the high-speed movies and the sections we will analyze (sectionsfile)
def load_event_data(startdate, endDT, startDT):
	# Load in the original events file
	# Tab separation vs spacing is KEY in the events file (LabView will fail if not right, and so will this code)
	lastAMorPM0 = None
	lastAMorPMcounter0 = 0
	hs_dpix = {}
	hs_pos = {}
	events = []
	#hs_dpix is a dictionary of numpy array for each high-speed movie event
	s = open(sectionsfile, 'r')
	slines = s.readlines()
	for sline in slines:
		# The EventSection object contains all of the high-speed events that are linked to each other
		# For example, the prepulse test has both strong tap, weak tap, and prepulse tap events in that section
		# Sections are designated by the user in the sections file and have a specific naming convention
		try:
			eventsection = EventSection(sline.strip(), startdate)
		except:
			print("The format of the input of your sections files is wrong. Every line should be as follows: habituation_daytaps=1_15:25:30-1_16:59:00")
			print("Exiting")
			exit()
		eventsection.check_endtime(endDT) # Makes sure that the last section is not after the true end of the data
		eventsection.check_starttime(startDT) # Makes sure that the last section is not after the true end of the data
		events.append(eventsection)
	f = open(eventsfile, 'r')
	lines = f.readlines()
	hscounter = 0
	#1:06:24\tPM\t0\t2\n'
	# Getting list of all the ID numbers associated with just the high-speed short movies
	highspeedlist = []
	for file in glob.glob(movieprefix + '*motion2'):
		highspeedlist.append(int(file.split('.avi.')[0].split('_')[-1]))
	highspeedlist.sort()
	for line in lines:
		TAPES = line.split('	')
		# Need to be splitting on tab only for the new format of the input file with the spaces in last column
		# Converting date to the format that we were using for the 
		#6/18/2016_12:59:34_PM
		dateplustime = startdate + "_" + TAPES[0][0:len(TAPES[0])] + "_" + TAPES[1]
		thistime = datetime.datetime.strptime(dateplustime, '%m/%d/%Y_%I:%M:%S_%p')
		#thistime = faststrptime(dateplustime)
		thisAMorPM0 = TAPES[1]
		if lastAMorPM0 != None:
			# If we transition either AM->PM or PM->AM
			# The only time we will ever have issues with this is if the events file doesn't correspond well with beginning timestamp input
			# Or there could be issues with start after midnight
			# These situations should not happen
			# Basically things become a mess if the program does not read the events file in as it should have and it skips starting at beginning
			if thisAMorPM0 != lastAMorPM0:
				if thisAMorPM0 == "AM":
					lastAMorPMcounter0 = lastAMorPMcounter0 + 1
		lastAMorPM0 = thisAMorPM0
		thistime = thistime + datetime.timedelta(days=(lastAMorPMcounter0))
		for eventsection in events:
			if eventsection.starttime <= thistime <= eventsection.endtime:
				# Debug
				# print(TAPES[2],TAPES[3].strip(';'),  thistime, e.starttime, e.endtime)
				eventsection.add_event(TAPES[2], TAPES[3].strip(';'), thistime)
		lasttime = thistime
		highspeedlist.sort()
		if int(TAPES[2]) != 0:
			# Load in only the high-speed movie data marked '1' and with the high-speed movie prefix
			# This would need to be modified if a labview code other than '1' was used to make high-speed movies that should analyzed in this way
			# Would also need to modify code in processmotiondata.py, specifically the CalculateEventProperties function relies on this "1" to trigger analysis
			if int(TAPES[2]) == 1:
				load_movie_motion(hs_dpix, thistime, highspeedlist[hscounter])
				load_movie_motion_pos(hs_pos, thistime, highspeedlist[hscounter])
				hscounter = hscounter + 1
	if hscounter != len(highspeedlist):
		print("ERROR, the number of high-speed movies does not correspond to the number expected based on the event file")
		print("Please make sure that the event file is accurate and run was fully completed. Modify events file to reflect reality of data output if needed")
		print("Exiting")
		exit()
	# Debug to make sure eveything is added correctly
	#for e in events:
	#	print("e: ", e.name, e.type, e.events)
	return (hs_dpix, hs_pos, events)


# The milliseconds are estimated based on the number of frames per that second
# It is clearly not accurate if a chunk of frames were lost in just the middle, for example
# However it is useful for downstream analyses to have milliseconds, in particular if one has to analyze responses in the slow speed data
def convert_to_ms_time(timestamp_data_array, timestamp_data_dict):
	mstimestamp_data_array = timestamp_data_array
	startt = timestamp_data_array[0]
	mseccounter = 0
	for position in range(0, len(timestamp_data_array)):
		if (position + 1) != len(timestamp_data_array):
			if timestamp_data_array[position] == timestamp_data_array[position + 1]:
				mseccounter = mseccounter + 1
			else:
				startpos = position - mseccounter
				msec = 1000.0 / ((position - startpos) + 1)
				for i in range(startpos, position + 1):
					newsec = timestamp_data_array[i] + datetime.timedelta(milliseconds = msec*(i-startpos))
					mstimestamp_data_array[i] = newsec
				mseccounter = 0
		else:
			mseccounter = mseccounter + 1
			startpos = position - mseccounter
			msec = 1000.0 / ((position - startpos) + 1)
			for i2 in range(startpos, position + 1):
				newsec = timestamp_data_array[i2] + datetime.timedelta(milliseconds = msec*(i2-startpos))
				mstimestamp_data_array[i2] = newsec
			mseccounter = 0
			break
	return mstimestamp_data_array


# The sections file is in 24 hour time, as is Python code
def load_timestamp_file():
	# Loading in the timestamp file data
	# Used to have to get rid of the ^M character that is between the times, but not with updated Anaconda
	timestamp_data_array = []
	dropped_seconds = []
	f = open(tstampfile, 'r')
	lines = f.readlines()
	f.close()
	for line in lines:
		timestamp_data_array.append(line.strip())
	n = 0
	timestamp_data_dict = {}
	lasttime = None
	starttime = None
	for t in timestamp_data_array:
		thistime = faststrptime(t)
		# I can't include the AM/PM in the faststrptime loading for speed reasons
		# So input times are assumed to just be AM (other than 12, assumed PM), and this adding/subtracting converts to 24 hour
		thisAMorPM0 = t.split()[len(t.split())-1]
		if thistime.hour == 12:
			if thisAMorPM0 == "AM":
				thistime = thistime + datetime.timedelta(hours = -12)
		elif thisAMorPM0 == "PM":
			thistime = thistime + datetime.timedelta(hours = 12)
		timestamp_data_array[n] = thistime
		timestamp_data_dict[thistime] = n
		# Account for situation at beginning, so we don't enter code below and try to subtract
		if n == 0:
			n = n +1
			lasttime = thistime
			starttime = thistime
			continue
		# This step is important later for the fast slicing of the data
		# Missing seconds need to be accounted for so we know not to expect it
		# Ideally we do not lose that amount of frames, but it can happen
		tdelta1 = thistime - lasttime
		testtime = thistime - datetime.timedelta(seconds = tdelta1.total_seconds())
		if thistime != lasttime:
			timestamp_data_dict[thistime] = n
			if tdelta1.total_seconds() > 1:
				for x in range(0,int(tdelta1.total_seconds()-1)):
					print("DROPPED A SECOND: ", thistime, lasttime, testtime, testtime + datetime.timedelta(seconds=1), timestamp_data_array[n], n-1, timestamp_data_array[n-1])
					dropped_seconds.append(testtime + datetime.timedelta(seconds=1))
					testtime = testtime + datetime.timedelta(seconds=1)
			lasttime = thistime
		n = n + 1
	mstimestamp_data_array = convert_to_ms_time(timestamp_data_array, timestamp_data_dict)
	return (mstimestamp_data_array, timestamp_data_dict, dropped_seconds, thistime, starttime)


# Find max and min value for each fish in order to identify well edges
# This is actually not really the well edges, but the edges of the fish's radius of movement
# I prefer this approach to reading in the originally designated ROI, just in case ROI isn't accurate (ie, includes extra plastic of well edge)
def max_min(cen_data_array):
	maxxys = []
	minxys = []
	for n in range (0, numberofwells*2,2):
		maxtest = np.amax(cen_data_array[:,n])
		mintest = np.amin(cen_data_array[:,n])
		# This is the check for if nothing ever moves (the well is empty)
		if maxtest == mintest and maxtest == 0:
			maxxys.append(0)
			maxxys.append(0)
			minxys.append(0)
			minxys.append(0)
		else:
			maxrealx = maxtest
			minrealx = np.amin(cen_data_array[:,n][np.nonzero(cen_data_array[:,n])])
			maxrealy = np.amax(cen_data_array[:,n+1])
			minrealy = np.amin(cen_data_array[:,n+1][np.nonzero(cen_data_array[:,n+1])])
			maxxys.append(maxrealx)
			maxxys.append(maxrealy)
			minxys.append(minrealx)
			minxys.append(minrealy)
	maxxysnp = np.array(maxxys)
	minxysnp = np.array(minxys)
	return( maxxysnp, minxysnp)


# Polar coordinates are essential for easy calculation of well edge/center preferences
def convert_to_polar(cen_data_array):
	(maxxysnp, minxysnp) = max_min(cen_data_array)
	midcoords = (maxxysnp + minxysnp) / 2
	midcoords = midcoords.astype(np.int16)
	cen_data_array = cen_data_array.astype(np.int16)
	cen_data_array[cen_data_array == 0] = -10000 # just setting to very low value to make it easier to skip later
	# subtract middle coordinate to get everything centered about 0
	zerodcoords = np.zeros(np.shape(cen_data_array))
	for i in range(0, numberofwells*2):
		zerodcoords[:,i] = cen_data_array[:,i] - midcoords[i]
	zerodcoords[zerodcoords < -5000 ] = 0
	# zerodcoords currently contains negative numbers, which I think mean that the fish hasn't moved yet
	thetadata = np.zeros((len(cen_data_array), numberofwells))
	rhodata = np.zeros((len(cen_data_array), numberofwells))
	xzerod = np.zeros((len(cen_data_array), numberofwells))
	yzerod = np.zeros((len(cen_data_array), numberofwells))
	for i in range(0, numberofwells):
		(rhodata[:,i], thetadata[:,i]) = cart2pol(zerodcoords[:,2*i], zerodcoords[:,2*i+1])
		xzerod[:,i] = zerodcoords[:,2*i]
		yzerod[:,i] = zerodcoords[:,2*i+1]
	return (rhodata, thetadata, xzerod, yzerod)


# The Fish object carries all the data around for each fish, including their genotype and ID number
# Later (after processmotiondata.py) the data inside this Fish object is analyzed (bouts counted, binned, responses counted) and the AnalyzedFish object carries that data
# This analysis code only compares two groups: a control group and a test group
# Or it can analyze a single group, but no statistics will be done
def generate_fish_objects(dp_data_array, rho_array, theta_array, x_array, y_array, hs_dpix, hs_pos, genotypefile):
	f = open(genotypefile, 'r')
	lines = f.readlines()
	f.close()
	genotype_list = {}
	for line in lines:
		# The file must use "controlgroup_geno: #,#,#" and "testgroup_geno: #,#,#,#" to emphasize to users that only two are allowed
		genogroup = line.split(':')[0].split('_')[0]
		if(len(line.split(':')[0].split('_')[0]) > 1):
			realgenotype = line.split(':')[0].split('_')[1]
		else:
			if((line.split(':')[0].split('_')[0]) == "controlgroup"):
				realgenotype = "control"
			elif((line.split(':')[0].split('_')[0]) == "testgroup"):
				realgenotype = "test"
			else:
				print("Not using correct nomenclature of controlgroup_genotype: and testgroup_genotype:")
				realgenotype = line.split(':')[0].split('_')[0]
		fishidslist = line.split(':')[1].strip().split(',')
		inputids = []
		for id in fishidslist:
			# Have to subtract 1 so that we can start from 0
			# Keeping it like this, but then adding 1 back to the ID that is saved
			inputids.append(int(id)-1)
		genotype_list[genogroup + "_" + realgenotype] = inputids
	fish_list = []
	for n in range(0, numberofwells):
		split_hs_dpix = {}
		split_hs_pos_x = {}
		split_hs_pos_y = {}
		for d in hs_dpix.keys():
			split_hs_dpix[d] = hs_dpix[d][:,n]
			split_hs_pos_x[d] = hs_pos[d][:,2*n]
			split_hs_pos_y[d] = hs_pos[d][:,2*n+1]
		for x in genotype_list.keys():
			if n in genotype_list[x]:
				# Adding 1 back onto the fish.idnumber, because all we use it for later is to connect to original input and we want it to match
				newfish = Fish(n + 1, x.split('_')[0], x.split('_')[1], dp_data_array[:,n], rho_array[:,n], theta_array[:,n], x_array[:,n], y_array[:,n], split_hs_dpix, split_hs_pos_x, split_hs_pos_y)
				fish_list.append(newfish)
	return fish_list


# Start here
def loading_procedures():
	tuple_timestamps = load_timestamp_file()
	print("Done loading timestamp file")
	
	with open(dpixfile, 'rb') as fid:
		dp_data_array = np.fromfile(fid, dtype = '>u2')
	dp_data_array = dp_data_array.reshape(dp_data_array.size // numberofwells, numberofwells)
	print("Done loading dpix")

	with open(centroidfile, 'rb') as fid:
		cen_data_array = np.fromfile(fid, '>u2')
	cen_data_array = cen_data_array.reshape(cen_data_array.size // (numberofwells*2), (numberofwells*2))
	cen_data_array[cen_data_array == 65535] = 0 # just setting to zero to make it easier to ignore
	# This is because they are all values of -16 and the initial type of the array is unsigned int, but it should be clear that it means the fish hasn't moved yet
	# converting them to zero for now so that it makes it easier to deal with the downstream max/min tests
	tuple_rho_theta = convert_to_polar(cen_data_array)
	print("Done converting to polar coordinates")

	# This is when the run started, calculated from the timestamp file and it is useful for putting event data on correct day
	# Day "0" in the sections file corresponds to this start date from the very beginning of the timestamp file
	startdate = str(tuple_timestamps[0][0].month) + "/" + str(tuple_timestamps[0][0].day) + "/" + str(tuple_timestamps[0][0].year)
	endDT = tuple_timestamps[3]
	startDT = tuple_timestamps[4]
	global_tuple_events = load_event_data(startdate, endDT, startDT)
	print("Done loading events")
	
	fish_list = generate_fish_objects(dp_data_array, tuple_rho_theta[0], tuple_rho_theta[1], tuple_rho_theta[2], tuple_rho_theta[3], global_tuple_events[0], global_tuple_events[1], genotypefile)
	return (fish_list, tuple_timestamps[0], tuple_timestamps[1], tuple_timestamps[2], global_tuple_events[2])

def initialize_args():
	print("Initializing arguments")
```
