## Supplemental Software for "A Customizable Low-Cost System for Massively Parallel Zebrafish Behavior Phenotyping": Fish.py

```
#!/usr/bin/python

# This is the fish, which starts out unanalyzed and then gains analyzed data
# It carries around genotype info and basic info about position in each frame of the slow- and high-speed data
class Fish:

	def __init__(self, idnumber, genogroup, realgenotype, dpix, rho_array, theta_array, x_array, y_array, hs_dict, hs_pos_x, hs_pos_y):

		self.idnumber = idnumber # string
		self.genogroup = genogroup # string
		self.realgenotype = realgenotype # string
		self.dpix = dpix # numpy array
		self.rho_array = rho_array # numpy array
		self.theta_array = theta_array # numpy array
		self.x_array = x_array # numpy array
		self.y_array = y_array # numpy array
		self.hs_dict = hs_dict # numpy array
		self.hs_pos_x = hs_pos_x # numpy array
		self.hs_pos_y = hs_pos_y # numpy array
		self.binned_data = []

	def add_binned_data(self, binned_data):
		for d in binned_data:
			self.binned_data.append(d)
```
