## Supplemental Software for "A Customizable Low-Cost System for Massively Parallel Zebrafish Behavior Phenotyping": graphsstatsandfilter.py

```
#!/usr/bin/python
import os,sys,glob,re
import numpy as np
import scipy
from scipy import stats
import datetime
import time
from datetime import timedelta
import matplotlib
matplotlib.use('Agg')
import matplotlib.pyplot as plt
from matplotlib import colors as c
from matplotlib  import cm
from scipy.stats.kde import gaussian_kde
from numpy import linspace
from scipy.stats import kruskal
#from scipy.stats import nanmean
#from scipy.stats import nanmedian
import pandas as pd
import statsmodels.api as sm
from scipy.stats import mstats
from matplotlib.ticker import FormatStrFormatter
from mpl_toolkits.axes_grid1 import make_axes_locatable
import ast
import math

# Helper

def fix_time(time):
	timetype = " sec"
	if 0.016 < float(time) / 60.0 < 1: # more than 1 sec, less than one min
		timetype = " / " + str(time) + " sec"
	elif 1 <= float(time) / 60.0 < 60:
		time = str(float(time) / 60.0) # time in minutes, not seconds
		if int(float(time)) == 1:
			timetype = " / min"
		else:
			timetype = " / " + str(int(float(time))) + " min"
	elif 60 <= float(time) / 60.0:
		time = str(float(time) / 3600) # time in hours, not seconds
		if int(float(time)) == 1:
			timetype = " / hour"
		else:
			timetype = " / " + str(int(float(time))) + " hour"
	return timetype

def fix_time_outer(intime):
	if "over" in intime:
		timenum = intime.split("over")[0]
		timedenom = intime.split("over")[1]
		return " " + fix_time(int(timenum)).split('/')[1].strip() + " / " + fix_time(timedenom).split('/')[1].strip()
	else:
		return fix_time(int(intime))

# Graphing

def hm_plot(intensity, type, ids, xaxis, yaxis, catarray):
	heatgraphname = "heatgraph_" + type +".png"
	x = range(0,len(intensity[0])+1)
	y = range(0,len(intensity)+1)
	x,y = np.meshgrid(x,y)
	intensity = np.array(intensity)
	fig = plt.figure()
	ax1 = fig.add_subplot(121)
	if "responsefull" in heatgraphname:
		xaxis = "Frames"
	ax1.set_xlabel(xaxis)
	ax1.set_ylabel("Fish ID Numbers")
	ax1.set_yticks(np.arange(len(ids)) + 0.5)
	ax1.set_yticklabels(reversed(ids))
	#ax1.set_yticklabels(reversed(ids), fontsize = 14)
	im = ax1.pcolormesh(x,y,intensity,cmap='hot',vmin=np.nanmin(catarray),vmax=np.nanmax(catarray))
	divider = make_axes_locatable(ax1)
	cax = divider.append_axes('bottom', size='10%', pad=0.6)
	plt.colorbar(im, cax=cax, orientation='horizontal')
	cax.set_xlabel(yaxis)
	plt.savefig(heatgraphname, transparent=True, format ="png")
	plt.close()


def box_plot(arrays, type, ylabel, genos):
	data = []
	# just want the "het" or "hom" or whatever label
	justgenos = []
	for g2 in genos:
		justgenos.append(g2.split('-')[1])
	for a in arrays:
		data.append(np.asarray(a))
	boxgraphname = "boxgraph_" + type + ".png"
	dictdata = {}
	for l in range(0, len(data)):
		if data[l].ndim > 1:
			mu1 = np.nanmean(data[l], axis=1)
		else:
			mu1 = data[l]
		dictdata[str(l)] = mu1
	df = pd.DataFrame(dict([ (k,pd.Series(v)) for k,v in dictdata.items() ]))
	df.columns = justgenos
	df.to_csv(boxgraphname + "_data.csv", sep='\t')
	plt.clf()
	plt.cla()
	fig = plt.figure()
	ax1 = fig.add_subplot(121)
	ax1.set_ylabel(ylabel)
	ax1.set_xticklabels(justgenos)
	ax1.set_axisbelow(True)
	plot = df.boxplot(ax=ax1)
	plt.savefig(boxgraphname, transparent=True, format="png")
	plt.close()


def ribbon_plot(arrays, type, ylabel, xlabel, t = None):
	colors = ['black','red','blue','yellow','purple','orange']
	ribgraphname = type + ".png"
	fig = plt.figure()
	ax1 = fig.add_subplot(121)
	if t == None:
		t = np.arange(np.shape(arrays[0])[1])
		if "responsefull" in ribgraphname:
			t = t * 3.5
			xlabel = "Time (msec)"
	for a in range(0,len(arrays)):
		arr = np.asarray(arrays[a])
		#array1 = np.asarray(arrays[0])
		#array2 = np.asarray(arrays[1])
		mu = np.nanmean(arr, axis=0)
		#mu1 = np.nanmean(array1, axis=0)
		sigma = stats.sem(arr, axis=0, nan_policy='omit')
		#sigma1 = stats.sem(array1, axis=0, nan_policy='omit')
		#mu2 = np.nanmean(array2, axis=0)
		#sigma2 = stats.sem(array2, axis=0, nan_policy='omit')
		ax1.plot(t, mu, lw=1, color = colors[a])
		#ax1.plot(t, mu1, lw=1, label = "mean wt", color = 'black')
		#ax1.plot(t, mu2, lw=1, label = "mean mut", color = 'red')
		ax1.fill_between(t, mu+sigma, mu-sigma, facecolor=colors[a], alpha=0.3)
		#ax1.fill_between(t, mu1+sigma1, mu1-sigma1, facecolor='black', alpha=0.3)
		#ax1.fill_between(t, mu2+sigma2, mu2-sigma2, facecolor='red', alpha=0.3)
	ax1.set_xlabel(xlabel)
	ax1.set_ylabel(ylabel)
	ax1.grid()
	fig.savefig(ribgraphname, transparent=True, format="png")
	plt.close()

# End Graphing


# Statistics

def linear_model_array(ribgraphname, array1, array2):
	fw = open(ribgraphname + "_newdata.csv", 'w')
	fw.write("time,movement,id,mutornot\n")
	for n in range(0, array1.shape[0]):
		t = 0
		for d in array1[n,:]:
			fw.write(str(t))
			fw.write(",")
			fw.write(str(d))
			fw.write(",")
			fw.write(str(n))
			fw.write(",wt")
			fw.write("\n")
			t = t+1
	for n2 in range(0, array2.shape[0]):
		t2 = 0
		for d2 in array2[n2,:]:
			fw.write(str(t2))
			fw.write(",")
			fw.write(str(d2))
			# just adding 100 to the id number, so that it's different from wt ids, since real ids are gone by now
			fw.write(",")
			fw.write(str(int(n2) + 100))
			fw.write(",mut")
			fw.write("\n")
			t2 = t2+1
	fw.close()
	data = pd.read_csv(ribgraphname + "_newdata.csv")
	data = data[data.movement.notnull()]
	model = sm.MixedLM.from_formula("movement ~ mutornot + time + mutornot * time", data, groups=data["id"])
	result = model.fit()
	print(ribgraphname)
	print(result.summary())

#ssmd = (meanwt - meanmut) / (math.sqrt(varwt + varmut)))
def anova(dataname, nparray1, nparray2):
	if nparray1.ndim > 1:
		nanmean1 = np.nanmean(np.nanmean(nparray1, axis=1))
		nanvar1 = np.nanvar(np.nanmean(nparray1, axis=1))
		nanmean2 = np.nanmean(np.nanmean(nparray2, axis=1))
		nanvar2 = np.nanvar(np.nanmean(nparray2, axis=1))
		H, pval = mstats.kruskalwallis(np.nanmean(nparray1, axis=1), np.nanmean(nparray2, axis=1))
		print("anova: ", dataname, ': N of control, test, Mean of array control, test, Variance of array control, test, SSMD, H-stat, P-value: ', len(np.nanmean(nparray1, axis=1)),len(np.nanmean(nparray2, axis=1)), str(nanmean1), str(nanmean2), str(nanvar1), str(nanvar2), str((nanmean1 - nanmean2) / math.sqrt(nanvar1+nanvar2)), str(H), str(pval))
	else:
		nanmean1 = np.nanmean(np.nanmean(nparray1))
		nanvar1 = np.nanvar(nparray1)
		nanmean2 = np.nanmean(np.nanmean(nparray2))
		nanvar2 = np.nanvar(nparray1)
		H, pval = mstats.kruskalwallis(nparray1, nparray2)
		print("anova: ", dataname, ': N of control, test, Mean of array control, test, Variance of array control, test, SSMD, H-stat, P-value: ', len(np.nanmean(nparray1)), len(np.nanmean(nparray2)), str(nanmean1), str(nanmean2), str(nanvar1), str(nanvar2), str((nanmean1 - nanmean2) / math.sqrt(nanvar1+nanvar2)), str(H), str(pval))

# End Statistics


def filter_obend(array, ribgraphname, header, filters):
	newnamev = []
	newnamel = []
	for r in ribgraphname.split("_"):
		if r.startswith("response"):
			newnamev.append("responsetime")
			newnamel.append("responsesumabsheadingangle")
		else:
			newnamev.append(r)
			newnamel.append(r)
	velarray = np.loadtxt("_".join(newnamev), delimiter = ',')
	latarray = np.loadtxt("_".join(newnamel), delimiter = ',')
	boolvel = velarray > filters[0]
	# Used to be latency/velocity, but now we filter these anyway based on the frames considered, so using new measure
	boollat = latarray > filters[1]
	boolmask = np.logical_and(boolvel, boollat)
	invboolmask = np.logical_not(np.logical_and(boolvel, boollat))
	mxw = np.ma.masked_array(array, mask=boolmask)
	imxw = np.ma.masked_array(array, mask=invboolmask)
	if "_responsefrequency_" in ribgraphname:
		imxw = np.ma.filled(imxw, 0)
		mxw = np.ma.filled(mxw, 0)
		imxw[np.isnan(mxw)] = np.nan
	else:
		imxw = np.ma.filled(imxw, np.nan)
		mxw = np.ma.filled(mxw, np.nan)
	bigname = "realescapes" + ribgraphname
	smallname = "notescapes" + ribgraphname
	np.savetxt(bigname, np.array(imxw,dtype=np.float64), delimiter = ',', header=header)
	np.savetxt(smallname, np.array(mxw,dtype=np.float64), delimiter = ',', header=header)


def filter_cbend(array, ribgraphname, header, filters):
	newnamev = []
	newnamel = []
	for r in ribgraphname.split("_"):
		if r.startswith("response"):
			newnamev.append("responsevelocity")
			newnamel.append("responsepeakdpix")
		else:
			newnamev.append(r)
			newnamel.append(r)
	velarray = np.loadtxt("_".join(newnamev), delimiter = ',')
	latarray = np.loadtxt("_".join(newnamel), delimiter = ',')
	boolvel = velarray > filters[0]
	boollat = latarray > filters[1]
	boolmask = np.logical_and(boolvel, boollat)
	invboolmask = np.logical_not(np.logical_and(boolvel, boollat))
	mxw = np.ma.masked_array(array, mask=boolmask)
	imxw = np.ma.masked_array(array, mask=invboolmask)
	if "_responsefrequency_" in ribgraphname:
		imxw = np.ma.filled(imxw, 0)
		mxw = np.ma.filled(mxw, 0)
		imxw[np.isnan(mxw)] = np.nan
	else:
		imxw = np.ma.filled(imxw, np.nan)
	mxw = np.ma.filled(mxw, np.nan)
	bigname = "realescapes" + ribgraphname
	smallname = "notescapes" + ribgraphname
	np.savetxt(bigname, np.array(imxw,dtype=np.float64), delimiter = ',', header=header)
	np.savetxt(smallname, np.array(mxw,dtype=np.float64), delimiter = ',', header=header)


def calc_stats(graphname, listofarrays,slowdata=False):
	tuplist = []
	# Normal situation is 2
	if(len(listofarrays) == 2):
		array1 = listofarrays[0]
		array2 = listofarrays[1]
		tuplist.append((array1, array2))
	elif(len(listofarrays) == 1):
		print("Cannot do statistics with a single dataset - make sure you have a control and a test group")
	elif(len(listofarrays) > 2):
		for l in range(0, len(listofarrays)):
			for l2 in range(0, len(listofarrays)):
				if l2 > l:
					tuplist.append(listofarrays[l], listofarrays[l2])
	for tup in tuplist:
		try:
			anova(graphname, tup[0],tup[1])
		except:
			print("anova failed: ", graphname)
		if(slowdata):
			if(len(listofarrays) == 2):
				try:
					linear_model_array(graphname, tup[0], tup[1])
				except:
					print("linear model failed: ", graphname)

def main(graphparametersfile, baselinelight, obendfilters, cbendfilters):
	# Create a dictionary out of the graphing parameters file
	#ribgraph_mean_time_day2nightppi_boutvelocity_600_het.data
	#ribgraph_mean_voltthreshold_day6dpfppinight_responsevelocity_1_a0f1400d5pD300a1f1400d5p_a1%89%97_hom.data
	genos = set()
	plotdict = {}
	with open(graphparametersfile) as f:
		for line in f:
			(key, val) = line.split(':')
			plotdict[key.strip()] = val.strip()
	for file1 in glob.glob("ribgraph*.data"): # Loop to get genotypes
		filename = file1.split(".")[0] 
		geno = filename.split("_")[-1]
		genos.add(geno)
	genos = list(genos)
	genos.sort()
	for file2 in glob.glob("ribgraph*.data"): # Loop to filter the big moves (cbends and obends) from less strong responses
		array = np.loadtxt(file2, delimiter=',')
		f0 = open(file2)
		header0 = f0.readline().strip()
		ids = header0.split(" ")[1].split("-")
		header = '-'.join(ids)
		if ("response" in file2) and ("responsefull" not in file2):
			stim = file2.split(".")[0].split("_")[-2]
			if stim.startswith("a"): # Acoustic stimulus
				filter_cbend(array, file2, header, cbendfilters)
			elif stim.startswith("b"): # Visual stimulus
				intensity = int(stim.split("%")[0].split("b")[1]) # The voltage of the light
				# Only calculating for reduction of light
				if intensity < baselinelight:
					filter_obend(array, file2, header, obendfilters)
			elif stim.startswith("v"): # Visual stimulus
				intensity = int(stim.split("%")[0].split("v")[1]) # The voltage of the light
				# Only calculating for reduction of light
				if intensity < baselinelight:
					filter_obend(array, file2, header, obendfilters)
	for file3 in glob.glob("*ribgraph*"+ genos[0] + ".data"): # Loop through (just one genotype) to make graphs and do statistical analyses
		# Make a list of the number of genotypes, if it's more than two give warning, if it's just one also give warning and don't do stats	
		arraylist = []
		idlist = []
		namelist = []
		# Get the header for the file that has the fish IDs in order
		f = open(file3)
		header = f.readline()
		ids = header.split(" ")[1].split("-")
		idlist.append(ids)
		namelist.append(file3.split('.')[0])
		array0 = np.loadtxt(file3, delimiter = ',')
		if np.size(array0) == 0: # if this is an empty dataset
			print("Empty dataset ", file3)
			continue
		arraylist.append(array0)
		# NEED TO DEAL WITH THIS SO THE CONTROL GROUP IS ONE THAT IS FIRST
		# It will be because "C" comes before "T" (control vs test), but should include a warning for people
		for g in range(1,len(genos)):
			f2 = open(file3.replace(genos[0],genos[g]))
			header2 = f2.readline()
			ids2 = header2.split(" ")[1].split("-")
			namelist.append(file3.replace(genos[0],genos[g]).split('.')[0])
			idlist.append(ids2)
			arraylist.append(np.loadtxt(file3.replace(genos[0], genos[g]), delimiter = ','))
		# Example names
		#ribgraph_mean_voltthreshold_day7dpfmsdf_responsevelocity_1_v120a001f1000d5pD995v200_a001%4%12_controlgroup-het.data
		#ribgraph_mean_time_day3heatshock_dpix_bouttime_600_testgroup-hom.png
		#ribgraph_mean_time_day3nightall_active_60over3600_testgroup-hom
		nolabel = True
		# Combine all data to generate a scale for heatmaps that goes across both datasets
		catarray = np.concatenate(arraylist)
		# Looking to see if shows up more than once, in which case we overwrite
		foundaxis = False
		similaraxes = []
		for plottype0 in plotdict.keys():
			if plottype0 in file3:
				if foundaxis == True:
					print("Found related axis more than once, going to overwrite axes and plot info with longest one")
					similaraxes.append(plottype0)
				foundaxis = True
		longlen = -1
		longtype = ""
		for xtype in similaraxes:
			if len(xtype) > longlen:
				longlen = len(xtype)
				longtype = xtype
		# Right now we are simply getting the yaxis from the input info, but in theory could get more than this if we expand the PlotParameters
		# As we get a lot of stuff, we may want fancier format and a function to get everything
		if longtype != "":
			yaxis = plotdict[longtype]
		else:
			foundaxis = False
			for plottype in plotdict.keys():
				if plottype in file3:
					yaxis = plotdict[plottype]
					foundaxis = True
			if foundaxis == False:
				print("Y-axis is not in the PlotParameters file for the following file: ", file3)
				yaxis = "No y-axis specified"
		if "_time_" in file3:
			if "_dpix_" in file3:
				yaxis = yaxis + " [dpix] "
			timetype = fix_time_outer(file3.split('.')[0].split('_')[-2])
			yaxis = yaxis + timetype
			for a in range(0, len(arraylist)):
				if(arraylist[a].ndim <2):
					arraylist[a] = np.reshape(arraylist[a], (-1, 1))
				hm_plot(arraylist[a], namelist[a], idlist[a], timetype.split(" / ")[-1].strip(), yaxis, catarray)
			box_plot(arraylist, '_'.join(file3.split('.')[0].split('_')[:-1]), yaxis, genos)
			ribbon_plot(arraylist, '_'.join(file3.split('.')[0].split('_')[:-1]), yaxis, "Time (" + timetype.split(" / ")[-1].strip() + ")")
			#ribgraph_mean_time_day2dfall_numberofbouts_3600_controlgroup-het.data
			calc_stats('_'.join(file3.split('.')[0].split('_')[:-1]), arraylist, True)
			nolabel = False
		else: # response graphs
			for a in range(0, len(arraylist)):
				if(arraylist[a].ndim <2):
					arraylist[a] = np.reshape(arraylist[a], (-1, 1))
				hm_plot(arraylist[a], namelist[a], idlist[a], "Events", yaxis, catarray)
			box_plot(arraylist, '_'.join(file3.split('.')[0].split('_')[:-1]), yaxis, genos)
			ribbon_plot(arraylist, '_'.join(file3.split('.')[0].split('_')[:-1]), yaxis, "Events")
			calc_stats('_'.join(file3.split('.')[0].split('_')[:-1]), arraylist, False)
			nolabel = False
		if(nolabel):
			print("Label for plot is not in the input label file! ", file3)
			box_plot(arraylist, '_'.join(file3.split('.')[0].split('_')[:-1]), "No label")
			ribbon_plot(arraylist, '_'.join(file3.split('.')[0].split('_')[:-1]), "No label")
```
