## Supplemental Software for "A Customizable Low-Cost System for Massively Parallel Zebrafish Behavior Phenotyping": ProcessedData.py

```
#!/usr/bin/python

# Processed data object, with access to binned_time((tuple that can be 2 or 1)), data itself, name, and then will add the graphing parameters to it (x-axis, y-axis), high-speed or slow-speed (default)
class ProcessedData:

	def __init__(self, name, binned_data, time_bin, event_type=""):

		self.name = name # string
		self.binned_data = binned_data # list
		self.time_bin = time_bin # tuple with one or two elements
		self.event_type = event_type
		self.slow_speed = (event_type == "") # Should give True if this is a high-speed event
```
