## Supplemental Software for "A Customizable Low-Cost System for Massively Parallel Zebrafish Behavior Phenotyping": processmotiondata.py

```
#!/usr/bin/python

import os,sys,glob,re
import numpy as np
import scipy
import datetime
import time
from datetime import timedelta
from ProcessedData import ProcessedData # ProcessedData object
from scipy.stats import norm
import copy
import math
import matplotlib
import cProfile

import fileloading # prepares all the files
import setupgraphsandsavedata
import graphsstatsandfilter

# Helper functions

# these frames need to be optimized for the multi-sensory integration paradigm in particular
# in the end these frame splittings might need to be associated with the type of event (ie, there could be different for MSI, at least the one with the 50 msec start delay on tap)
# These frames will be set for the window of starting to look for an event. Then a started event can continue through until 14 more frames
# When deciding if a fish just moved and is therefore not valid, would want to look back a certain number of frames, maybe even 300 msec, so that would automatically remove all prepulse responses unless we increase the distance between them. Maybe it's best to eliminate all fish that happened to move in the window starting
def setup_frames(ekinput):
	start = ekinput.split('%')[1]
	end = ekinput.split('%')[2]
	return range(int(start), int(end))	


# Formuala is square root of r1^2 + r2^2 - 2r1r2cos(theta1-theta2)
# Don't really need this now that I'm saving the x,y coordinates also, but will just keep anyway
def polar_to_distance(rho1, rho2, theta1, theta2):
	# Large values of rho means that the fish has not moved yet
	if rho1 > 5000 or rho2 > 5000:
		dist = 0.0
	elif rho1 != rho2 or theta1 != theta2:
		dist = math.sqrt(rho1**2 + rho2**2 - 2*rho1*rho2*np.cos(theta1 - theta2))
		#dist = np.sqrt(rho1**2 + rho2**2 - 2*rho1*rho2*np.cos(theta1 - theta2))
	else:
		dist = 0.0
	return dist


def cart_to_distance(x1, x2, y1, y2):
	if x1 < -4000 or x2 < -5000:
		dist = 0.0
	elif x1 != x2 or y1 != y2:
		dist = math.sqrt((x2-x1)**2 + (y2 - y1)**2)
	else:
		dist = 0.0
	return abs(dist)

# End Helper functions


def PolyArea(xarr,yarr): # x and y are the coordinate lists for bout start to bout end
	pA = 0.5*np.abs(np.dot(xarr,np.roll(yarr,1))-np.dot(yarr,np.roll(xarr,1)))
	if pA < 300:
		# Get crazy high values sometimes, maybe not a complete polygon shape with all needed points
		return 0.5*np.abs(np.dot(xarr,np.roll(yarr,1))-np.dot(yarr,np.roll(xarr,1)))
	else:
		return np.nan

# For getting the start and end index of each event section, depending on how it was binned
def find_indices(timestart, timeend, timestamp_data_dict, startindices):
	startindex = timestamp_data_dict[timestart]
	endindex = timestamp_data_dict[timeend]
	shortstartindex = np.searchsorted(startindices[::2], startindex) - 1
	shortendindex = np.searchsorted(startindices[::2], endindex) - 1
	return shortstartindex, shortendindex

# Calculate heading angle
def HeadingAngle(xarr,yarr,bout_start,bout_end, hs=False):
	angles = []
	angvels = []
	initdir = 1
	for i in range(bout_start, (bout_end-1)):
		a = np.array([xarr[i],yarr[i]])
		b = np.array([xarr[i+1],yarr[i+1]])
		c = np.array([xarr[i+2],yarr[i+2]])
		ba = b - a
		bc = b - c
		bc_atan = np.arctan2(np.cross(ba,bc), np.dot(ba,bc))
		if np.abs(bc_atan) != 0:
			turnangle = (np.pi - np.abs(bc_atan)) * np.sign(bc_atan)
		else:
			turnangle = 0
		angles.append(turnangle)
		# Direction of initial turn
		# Negative signed turns are counterclockwise, positive are clockwise
		if i == bout_start:
			if np.abs(bc_atan) != 0:
				initdir = initdir * np.sign(bc_atan)
			else:
				initdir = 0
	return np.sum(angles), np.sum(np.abs(angles)), initdir


# Calculate the revolutions in the well for the bout - important for potential seizures
# This is different from heading angle because it is based on the center of the well (looking for animals that circle well edges)
#  if ((fish_rev > 4.0) and (300 < fish_speed < 1300) and (fish_dist > 70)) = potential seizure
def calculate_boutrev(bout_start, bout_end, unsignedthetas, rhos):
	thetasum = 0
	for r2 in range(bout_start, bout_end):
		currenttheta = unsignedthetas[r2]
		futuretheta = unsignedthetas[r2 + 1]
		thetasub = futuretheta - currenttheta
		if thetasub > np.pi:
			thetasub = thetasub - (2 * np.pi)
		if thetasub < -np.pi:
			thetasub = thetasub + (2 * np.pi)
		thetasum = thetasum + thetasub
	return(abs(thetasum))


# Calculate bout angular velocity - this is interesting for potential seizures, but less so on a summarized fish level
# This is different from heading angle because it is based on the center of the well (looking for animals that circle well edges)
def calculate_boutanglevel(bout_start, bout_end, thetas):
	anglevelframe = []
	for r3 in thetas[bout_start:(bout_end+1)]:
		anglevelframe.append(abs(float(r3) / float(fileloading.msecperframe)))
	aveanglevel = (float(sum(anglevelframe)) / float(len(anglevelframe)))
	peakanglevel = max(anglevelframe)
	return aveanglevel,peakanglevel


# Calculate the preferences for the center vs edge for bouts and interbouts
def calculate_centerfrac(bout_start, bout_end, interbout_start, interbout_end, rhos, rhomax, halfpt):
	cmoments = 0
	intercmoments = 0
	tmoments = 0
	intertmoments = 0
	centerfrac = 0.0
	intercenterfrac = 0.0
	averhofrac = 0.0
	interaverhofrac = 0.0
	totalrho = 0.0
	intertotalrho = 0.0
	for r3 in rhos[bout_start:(bout_end+1)]:
		totalrho = r3 + totalrho
		if r3 < halfpt:
			cmoments = cmoments + 1
		tmoments = tmoments + 1
	centerfrac = float(cmoments) / float(tmoments)
	averhofrac = (float(totalrho) / float(tmoments)) / float(rhomax)
	for r2 in rhos[interbout_start:(interbout_end+1)]:
		intertotalrho = r2 + intertotalrho
		if r2 < halfpt:
			intercmoments = intercmoments + 1
		intertmoments = intertmoments + 1
	# Only comes up if there is a bout at end of movie, because then the interbout_end and _start are the same and don't really exist
	# None of this data will end up being analyzed though, because we ignore the last seconds usually anyway
	if intertmoments != 0.0:
		intercenterfrac = float(intercmoments) / float(intertmoments)
		interaverhofrac = (float(intertotalrho) / float(intertmoments)) / float(rhomax)
	else:
		intercenterfrac = 0.0
		interaverhofrac = 0.0
	return centerfrac, averhofrac, intercenterfrac, interaverhofrac


# Calculate distances moved in every frame of high-speed movie data
def hs_calculate_distance(fish):
	hs_dist = {}
	for timekey in fish.hs_pos_x:
		xlist = []
		ylist = []
		for x in range(0, len(fish.hs_pos_x[timekey])):
			xlist.append(fish.hs_pos_x[timekey][x])
			ylist.append(fish.hs_pos_y[timekey][x])
		hs_dist[timekey] = np.zeros(np.shape(np.asarray(xlist)), dtype=float)
		hs_dist[timekey][0] = 0.0
		for d in range(0, (len(xlist)-1)):
			if (xlist[d] - xlist[d+1] == 0.0) and (ylist[d] - ylist[d+1] == 0.0):
				hs_dist[timekey][d+1] = 0.0
			else:
				hs_dist[timekey][d+1] = cart_to_distance(xlist[d], xlist[d+1], ylist[d], ylist[d+1])
	return hs_dist


# Calculate distances moved in every frame of slow-speed data
def calculate_distance(fish):
	# distance unit is in pixels
	rhos = fish.rho_array
	thetas = fish.theta_array
	distances = np.zeros(np.shape(rhos), dtype=float)
	# at first the index the distance traveled is 0,0
	distances[0] = 0.0
	for r in range(0, (len(rhos)-1)):
		if (rhos[r] - rhos[r+1] == 0.0) and (thetas[r] - thetas[r+1] == 0.0):
			distances[r+1] = 0.0
		else:
			distances[r+1] = polar_to_distance(rhos[r], rhos[r+1], thetas[r], thetas[r+1])
	return distances


# This function is where you would add in additional measures, if you want something new for slow-speed data
def CalculateBoutProperties(fish_distancesordpix, timestamp_data_array, boutstarts, boutends, rhos, thetas, xarray, yarray, prefix = "", rhothetadone = False):
	# Constants needed
	rhomax = np.amax(rhos)
	halfpt = rhomax * 0.45 # this is what we are calling the halfpt
	unsignedthetas = thetas + np.pi
	
	centerfracs = []
	intercenterfracs = []
	averhofracs = []
	interaverhofracs = []
	
	bouttimes = []
	boutdistancesordpix = []
	boutdisplacements = []
	interbouttimes = []
	
	sleepbouts = [] # list of tuples for filtering waking activity

	boutpropertieswithpre = {}
	boutproperties = {
			"bouttime": [],
			"interbouttime": [],
			"numberofboutsSLEEP": [], # interbouts of > 60 seconds
			"fracboutoverinterbout": [],
			"boutspeed": [],
			"boutcumulativemovement": [],
			"boutdisplacement": [],
			"boutvelocity": [],
			"boutaveangvelocity": [],
			"boutpeakangvelocity": [],
			"boutrevolutions": [],
			"boutseizurecount": [],
			"boutsumheadingangle": [], # Doesn't make a lot of sense, but it comes along with calculating the unsigned total
			"boutsumabsheadingangle": [],
			"fracconsistentinitdirectionheadingangle": [], # In this function we aren't appending the %, only the actual clockwise or counterclockwise (1 or -1), but later we calculate it and don't want to mess with name change later
			"boutrhofraction": [],
			"interboutrhofraction": [],
			"boutcenterfraction": [],
			"interboutcenterfraction": [],
			"numberofbouts": []
			}

	for b in range(0, len(boutstarts)):
		bout_start = boutstarts[b]
		bout_end = boutends[b]
		boutproperties["numberofbouts"].append(1)
		bouttime = (timestamp_data_array[boutends[b]] - timestamp_data_array[boutstarts[b]]).total_seconds() * 1000
		boutproperties["bouttime"].append(bouttime)
		boutdistanceordpix = 0
		for c in range(boutstarts[b], boutends[b]):
			boutdistanceordpix = boutdistanceordpix + fish_distancesordpix[c]
		boutproperties["boutcumulativemovement"].append(boutdistanceordpix)
		# Speeds can be applicable to dpix data (vs. displacement is not) because cumdpix / time could be interesting
		if (bouttime != 0):
			boutproperties["boutspeed"].append(float(boutdistanceordpix) / float(bouttime))
		else:
			print("Bout time appears to be 0 for some reason, boutstart, boutend: ", boutstarts[b], timestamp_data_array[boutends[b]], boutends[b], timestamp_data_array[boutstarts[b]], len(timestamp_data_array))
			boutproperties["boutspeed"].append(0.0)
		interbout_start = boutends[b] + 1
		if b != (len(boutstarts)-1):
			interbout_end = boutstarts[b+1] - 1
		else:
			interbout_end = np.shape(rhos)[0]
		# Each bout also gets an interbout that follows it (even last one, goes until end of data)
		# So the number of items in every list passed back should be exactly the same
		IBtime = (timestamp_data_array[interbout_end] - timestamp_data_array[interbout_start]).total_seconds() * 1000
		boutproperties["interbouttime"].append(IBtime)
		if (IBtime != 0):
			boutproperties["fracboutoverinterbout"].append(float(bouttime) / float(IBtime))
		else:
			boutproperties["fracboutoverinterbout"].append(0.0)
		if IBtime > 60000:
			boutproperties["numberofboutsSLEEP"].append(1)
			sleepbouts.extend(list(range(interbout_start,interbout_end)))
		else:
			boutproperties["numberofboutsSLEEP"].append(0)
		# The call of this function that is done second (the dpix one) will already have completed this and the lists will be empty
		if( rhothetadone == False):
			boutdisplacement = polar_to_distance(rhos[bout_start], rhos[bout_end], thetas[bout_start], thetas[bout_end])
			boutproperties["boutdisplacement"].append(boutdisplacement)
			boutproperties["boutvelocity"].append(float(boutdisplacement) / float(bouttime))
			aveanglevel,peakanglevel = calculate_boutanglevel(bout_start, bout_end, thetas)
			boutproperties["boutaveangvelocity"].append(aveanglevel)
			boutproperties["boutpeakangvelocity"].append(peakanglevel)
			boutrev = calculate_boutrev(bout_start, bout_end, unsignedthetas, rhos)
			boutproperties["boutrevolutions"].append(boutrev)
			
			#if ((boutrev > 4.0) and (300 < (bouttime / boutdistanceordpix) < 1300) and (boutdistanceordpix > 70)):
			if ((boutrev > fileloading.seizurefilters[0]) and (fileloading.seizurefilters[1] < (float(bouttime) / float(boutdistanceordpix)) < fileloading.seizurefilters[2]) and (fileloading.seizurefilters[3] > 70)):
				boutproperties["boutseizurecount"].append(1)
			else:
				boutproperties["boutseizurecount"].append(0)
			sumheading,sumabsheading,initialdirection = HeadingAngle(xarray,yarray,bout_start,bout_end)
			boutproperties["boutsumheadingangle"].append(sumheading)
			boutproperties["boutsumabsheadingangle"].append(sumabsheading)
			boutproperties["fracconsistentinitdirectionheadingangle"].append(initialdirection)
			centerfrac,averhofrac,intercenterfrac,interaverhofrac = calculate_centerfrac(bout_start, bout_end, interbout_start, interbout_end, rhos, rhomax, halfpt)
			boutproperties["boutrhofraction"].append(averhofrac)
			boutproperties["interboutrhofraction"].append(interaverhofrac)
			boutproperties["boutcenterfraction"].append(centerfrac)
			boutproperties["interboutcenterfraction"].append(intercenterfrac)
	for k in boutproperties.keys():
		boutpropertieswithpre[prefix + k] = boutproperties[k]
	return boutpropertieswithpre, sleepbouts


# Identify bouts, including not counting if there is noise of a single frame below threshold and filtering by number of frames in bout
def findBoutIndices(fish_distances, threshold, frames):
	boutstarts = []
	boutstarts2 = []
	realboutstarts = []
	boutends = []
	realboutends = []
	laststart = -1
	finallaststart = -1
	for d in range(0, len(fish_distances)-1):
		# Is value more than threshold and the last bout was determined finished
		# Could be the start of a bout, but it might not be if it doesn't pass frame cutoff
		if (fish_distances[d] >= threshold) and (laststart == -1):
			laststart = d
			finallaststart = d
		elif(laststart != -1):
			if fish_distances[d] < threshold and fish_distances[d+1] < threshold:
				if ((d-1) - laststart) >= frames:
					boutstarts.append(laststart)
					boutends.append(d-1)
				laststart = -1
	if fish_distances[len(fish_distances)-1] >=threshold:
		# If the bout was still happening when we stopped using camera
		boutstarts.append(finallaststart)
		boutends.append(len(fish_distances)-1)
	return boutstarts,boutends


# COULD UPDATE THIS FXN TO BE BETTER, LIKE THE SLOW-SPEED ONE, BUT NOT RIGHT NOW	
# IT WORKS FINE, JUST COULD BE MORE CLEANLY WRITTEN AND EASIER TO UNDERSTAND
def identify_event_bout(dpix_movement, approximate_frames, threshold, frame_threshold, highspeed=True):
	bout_start_fr = 0
	bout_end_fr = 0
	earlybout = False
	if approximate_frames[0] > 0:
		for b0 in range(0, approximate_frames[0] - 3): # Assumes that there is a buffer at the beginning of at least three frames before the expected motion
			if dpix_movement[b0] > threshold and dpix_movement[b0+1] > threshold and dpix_movement[b0+2] > threshold: # has to be real motion and not a flicker 
				earlybout = True
	# If fish is moving before they are supposed to based on the frame
	# In theory this could be wrong and lose data if you have a dramatically shortened response latency or have not optimized frame choices vs frame rate of camera
	if earlybout == True:
		# FUTURE: could also rework to find ALL "bouts" in the movies and then align with the start in the frames of interest (could also fuse if there are issues with broken bouts)
		# Let's keep it like this, but still keep the frame and intensity threshold for identifying bouts so we aren't tossing out fish who just shifted a tiny bit
		return (np.nan, np.nan)
	else:
		for d in range(0, len(approximate_frames)):
			if dpix_movement[approximate_frames[d]] >= threshold:
				bout_start_fr = approximate_frames[d]
				break
		if highspeed == True:
			for b in range(bout_start_fr, len(dpix_movement) - 4): # -4 seems arbitrary, this is just for any issues at end of movie or changing intensities?
				if b == 0:
					break
				if dpix_movement[b] < threshold and dpix_movement[b+1] < threshold and dpix_movement[b+2] < threshold: # really sure that bout is over (the visual ones tend to fail)
					bout_end_fr = b - 1
					break
				bout_end_fr = b - 1 # NEED THIS IN CASE THE BOUT STARTS AND THEN WE DONT HAVE ENOUGH FRAMES TO FINISH IT, SO YOU DONT END UP WITH END BEING BEFORE START
			if bout_end_fr - bout_start_fr>frame_threshold:
				return (bout_start_fr, bout_end_fr)
			else:
				return (0, 0)
		else: # NOT SURE HOW WELL THE HIGH-SPEED WORKS FOR CHECKING THE MOVING FRAMES . . . SEEMS LIKE NOT WELL, NON-HIGHSPEED DATA IS NOT REALLY WORTH LOOKING AT
		# This is legacy code, in the off-chance people want to look at responses on data that is a low FPS . . . not advisable
			for b2 in range(bout_start_fr, approximate_frames[-1] + len(approximate_frames)):
				if b2 == 0:
					break
				if dpix_movement[b2] < threshold:
					bout_end_fr = b2 - 1
					break
				bout_end_fr = b2 - 1 # NEED THIS IN CASE THE BOUT STARTS AND THEN WE DONT HAVE ENOUGH FRAMES TO FINISH IT, SO YOU DONT END UP WITH END BEING BEFORE START
			#bout_start_fr = bout_start_fr - 1 # Need this to help keep distance bouts that are only a single frame, no longer need it though if basing all on dpix - legacy code
			return (bout_start_fr, bout_end_fr)


def CalculateEventProperties(name, eventsection, hs_dict, hs_distances, threshold, frame_threshold, hs_pos_x, hs_pos_y):
	binnedlist = []
	# The keys for the event defines the kind of event (i.e. the info in teensy string)
	for event in eventsection.keys():
		responseproperties = {
		"responsefrequency": [],
		"responsedisplacement": [],
		"responselatency": [],
		"responsecumulativedistance": [],
		"responsecumulativedpix": [],
		"responsevelocity": [],
		"responsespeed": [],
		"responsetime": [],
		"responsepolygonarea": [],
		"responsesumheadingangle": [],
		"responsesumabsheadingangle": [],
		"responseinitdirectionheadingangle":[],
		"responsepeakspeed": [],
		"responsepeakdpix": [],
		"responsefulldata": [],
		"responsefulldpixdata": []
		}
		approximate_frames = setup_frames(event.split("_")[2])
		startframe = approximate_frames[0]
		endframe = approximate_frames[-1] + 14 # THIS IS PROBABLY SILLY, COULD JUST DO THE ENTIRE 285 FRAMES AND VIEW THAT
		if (event.split("_")[0] != "1"): # This is the code for the teensy to say high-speed movie
			continue
		for eventtime in eventsection[event]: # time key used in hs_distances
			# Bout are identified only on dpix data, not on distance data, but then the frames of the dpix responses are used to calculate the distance information
			# COULD UPDATE THIS FXN TO BE BETTER, LIKE THE SLOW-SPEED ONE, BUT NOT RIGHT NOW	
			(dpix_bout_start, dpix_bout_end) = identify_event_bout(hs_dict[eventtime], approximate_frames, threshold, frame_threshold)
			if (np.isnan(dpix_bout_end) or np.isnan(dpix_bout_start)):
				for k,vlist in responseproperties.items():
					if("responsefull" in k):
						vlist.append(np.full((endframe-startframe), np.nan))
					else:
						vlist.append(np.nan)
			elif (dpix_bout_start == 0 and dpix_bout_end == 0):
				for k2,vlist2 in responseproperties.items():
					if("responsefull" in k2):
						vlist2.append(np.full((endframe-startframe), np.nan))
					elif("frequency" in k2):
						vlist2.append(0)
					else:
						vlist2.append(np.nan)
			else:
				bout_time = (dpix_bout_end - dpix_bout_start) * fileloading.msecperframe
				responseproperties["responselatency"].append((dpix_bout_start-startframe) * fileloading.msecperframe)
				responseproperties["responsetime"].append(bout_time)
				responseproperties["responsefrequency"].append(1)
				bout_disp = cart_to_distance(hs_pos_x[eventtime][dpix_bout_start], hs_pos_x[eventtime][dpix_bout_end], hs_pos_y[eventtime][dpix_bout_start], hs_pos_y[eventtime][dpix_bout_end])
				responseproperties["responsedisplacement"].append(bout_disp)
				responseproperties["responsepolygonarea"].append(PolyArea(hs_pos_x[eventtime][dpix_bout_start:dpix_bout_end], hs_pos_y[eventtime][dpix_bout_start:dpix_bout_end]))
				sumHA,sumabsHA,initdirHA = HeadingAngle(hs_pos_x[eventtime], hs_pos_y[eventtime], dpix_bout_start, dpix_bout_end, True)
				responseproperties["responsesumheadingangle"].append(sumHA)
				responseproperties["responsesumabsheadingangle"].append(sumabsHA)
				responseproperties["responseinitdirectionheadingangle"].append(initdirHA)
				#responseproperties["responseinitsumabsha"].append(initsumabs)
				bout_tdist = 0
				bout_cumdpix = 0
				peak_dpix = 0
				peak_dist = 0
				for c0 in range(dpix_bout_start, dpix_bout_end):
					bout_tdist = bout_tdist + hs_distances[eventtime][c0]
					bout_cumdpix = bout_cumdpix + hs_dict[eventtime][c0]
					if hs_distances[eventtime][c0] > peak_dist:
						peak_dist = float(hs_distances[eventtime][c0])
					if hs_dict[eventtime][c0] > peak_dpix:
						peak_dpix = float(hs_dict[eventtime][c0])
				peak_speed = float(peak_dist) / float(fileloading.msecperframe)
				responseproperties["responsepeakspeed"].append(peak_speed)
				responseproperties["responsepeakdpix"].append(peak_dpix)
				responseproperties["responsecumulativedistance"].append(bout_tdist)
				responseproperties["responsecumulativedpix"].append(bout_cumdpix)
				responseproperties["responsespeed"].append(float(bout_tdist) / float(bout_time))
				responseproperties["responsevelocity"].append( float(bout_disp) / float(bout_time))
				responseproperties["responsefulldpixdata"].append(hs_dict[eventtime][startframe:endframe])
				responseproperties["responsefulldata"].append(hs_distances[eventtime][startframe:endframe])
		for k3,v3 in responseproperties.items():
			if len(v3) == 0:
				continue
			if("responsefull" in k3):
				binnedlist.append(ProcessedData(k3,np.nanmean(v3,axis=0), (event,),name))
			else:
				binnedlist.append(ProcessedData(k3,v3, (event,),name))
	return binnedlist


# Finding the index values of various chunks of time we are analyzing for movement
# Called by activity_time_data
def determine_indices(timestamp_data_array, timeinterval, timestart, timeend, timestamp_data_dict, dropped_seconds):
	indexstart = 0
	indexend = 0
	tdelta = timeend - timestart
	intervalnumber = int(tdelta.total_seconds()) / timeinterval # How many different chunks of time to break it into (ie, 10 minutes in active minutes per 10 minutes)
	indexstart = timestamp_data_dict[timestart] # The index value for the first occurrence of the start time, so this is the inclusive value
	indexend = len(timestamp_data_array) # The index value for the last occurrence of the end time, so this is the inclusive value
	intervalindices = []
	intervalindices.append(indexstart)
	timetest = timestart
	timestamp_data_dict_keys = timestamp_data_dict.keys()
	if timeinterval >= 1:
		for z2 in range(0,int(intervalnumber)):
			z2 = z2+1
			timetest1 = timetest + datetime.timedelta(seconds=(timeinterval * z2))
			if timetest1 not in dropped_seconds:
				indext = timestamp_data_dict[timetest1]
				intervalindices.append(indext-1)
				intervalindices.append(indext)
	else: # If we are going to be doing every frame instead of by a set time, can't go less than second intervals using the time approach
		# Therefore this part of the code is not relevant really we don't want to do less than one second intervals
		for indext in range(indexstart+1,indexend): # Need to add the one because I already put the first index in earlier
			intervalindices.append(indext)
	intervalindices = np.array(intervalindices, dtype=int) # searchsorted efficiency depends on having same datatype
	return intervalindices

# waking activity, or active sec / hr that are not inactive minutes
# data_array has already been copied at initial function call
def sleep(data_array, threshold, sleepbouts, intervalindices1, intervalindices3600, prefix = ""):
	activitydict = {
			prefix + "wakingactive":[]
			}
	if len(sleepbouts) > 0:
		sleepbouts = np.array(sleepbouts)
	maska = np.zeros(np.shape(data_array)[0], dtype = bool)
	maska[sleepbouts] = True
	data_array[maska] = np.nan
	binnedlist = []
	for m in range(0,len(intervalindices3600)-1,2):
		activetime = 0
		printdata = []
		startindex1 = np.searchsorted(intervalindices1, intervalindices3600[m])
		endindex1 = np.searchsorted(intervalindices1, intervalindices3600[m+1]) + 1 # Add one because range below is non-inclusive
		shortarray1 = intervalindices1[startindex1:endindex1]
		for m3 in range(0, len(shortarray1)-1,2):
			activetimebool = False
			for m4 in range(shortarray1[m3], shortarray1[m3+1]+1):
				if data_array[m4] > threshold:
					activetimebool = True
				if activetimebool == True:
					activetime = activetime + 1
					break
		activitydict[prefix + "wakingactive"].append(activetime)
	for k2,v2 in activitydict.items():
		binnedlist.append(ProcessedData(k2,v2, ("1","3600")))
	return binnedlist


def flexactivity(data_array, threshold, intervalnumindices, intervalindices, timebintup, prefix = ""):
	activitydict = {
			prefix + "active": []
			}
	binnedlist = []
	for m in range(0,len(intervalindices)-1,2):
		activetime = 0
		printdata = []
		startindex = np.searchsorted(intervalnumindices, intervalindices[m])
		endindex = np.searchsorted(intervalnumindices, intervalindices[m+1]) + 1 # Add one because range below is non-inclusive
		shortarray = intervalnumindices[startindex:endindex]
		for m3 in range(0, len(shortarray)-1,2):
			activetimebool = False
			for m4 in range(shortarray[m3], shortarray[m3+1]+1):
				if data_array[m4] > threshold:
					activetimebool = True
				if activetimebool == True:
					activetime = activetime + 1
					break
		activitydict[prefix + "active"].append(activetime)
	for k2,v2 in activitydict.items():
		binnedlist.append(ProcessedData(k2,v2, timebintup))
	return binnedlist

def bout_flexactivity(boutproperties, bout_startsl0, intervalindices0, timebin):
	intervalindices = np.array(intervalindices0, dtype = int)
	bout_startsl = np.array(bout_startsl0, dtype = int)
	binnedlist = []
	datadict = {}
	for k0 in boutproperties.keys():
		datadict[k0] = []
	# indexstart, indexend, which is why we need to traverse by 2
	for m in range(0,len(intervalindices)-1,2):
		testintarray_start = np.searchsorted(bout_startsl, intervalindices[m])
		testintarray_end = np.searchsorted(bout_startsl, intervalindices[m+1])
		# Make an empty dictionary with same keys as the original bout data and with empty lists
		# This new dictionary will instead contain the binned data
		for k,v in boutproperties.items():
			shortlist = np.array(v[testintarray_start:testintarray_end])
			if "numberofbouts" in k:
				datadict[k].append(np.sum(shortlist))
			elif "seizurecount" in k:
				datadict[k].append(np.sum(shortlist))
			# Persistence or % in same direction
			elif "initdirectionheadingangle" in k:
				if(np.count_nonzero(~np.isnan(shortlist)) > 0):
					per_clock = float(np.count_nonzero(~np.isnan(shortlist[shortlist > 0]))) / float(np.count_nonzero(~np.isnan(shortlist)))
					per_cclock = float(np.count_nonzero(~np.isnan(shortlist[shortlist < 0]))) / float(np.count_nonzero(~np.isnan(shortlist)))
					if per_clock > per_cclock:
						datadict[k].append(per_clock)
					else:
						datadict[k].append(per_cclock)
				else:
					datadict[k].append(np.nan)
			else:
				datadict[k].append(np.nanmean(shortlist))
	for k2,v2 in datadict.items():
		if(len(v2) == 0):
			continue
		if(np.isnan(v2).all()):
			continue;
		binnedlist.append(ProcessedData(k2,v2, (timebin,)))
	return binnedlist


# Main function that instantiates everything
def process_all_data():
	# Leaving in to remind how to profile specific regions of code
	#pr = cProfile.Profile()
	#pr.enable()
	# Load in all the data through the fileloading.py script
	(fish_list, timestamp_data_array, timestamp_data_dict, dropped_seconds, eventsectionlist) = fileloading.loading_procedures()
	all_fish_bouts = {}
	dpix_all_fish_bouts = {}
	
	# Obtain indices for the binning used to analyze bouts and also activity (classic sleep plots) data
	timestart = timestamp_data_array[0]
	timeend = timestamp_data_array[len(timestamp_data_array)-1]
	allbins = fileloading.activitytimes + fileloading.boutbins
	allbinsset = set(allbins)
	indexdict = {}
	for t in allbinsset:
		binindices = determine_indices(timestamp_data_array, int(t), timestart, timeend, timestamp_data_dict, dropped_seconds)
		indexdict[str(t)] = binindices
		# Adding the start and end indices for all possible binning for every eventsection, even if you don't need every single one (the numerator bins aren't used)
		for es1 in eventsectionlist:
			indstart, indend = find_indices(es1.starttime, es1.endtime, timestamp_data_dict, binindices)	
			es1.add_indices(t, indstart, indend)
	# Finding the bouts, both with distance and dpix, for each fish
	for fish in fish_list:
		# Calculating the distances moved between each frame for both slow-speed data and high-speed movies
		fish_distances = calculate_distance(fish)
		hs_fish_distances = hs_calculate_distance(fish)
		boutstarts,boutends = findBoutIndices(fish_distances, fileloading.thresholdvalues[0], fileloading.thresholdframes[0])
		boutproperties, sleepbouts = CalculateBoutProperties(fish_distances, timestamp_data_array, boutstarts, boutends, fish.rho_array, fish.theta_array, fish.x_array, fish.y_array)
		dboutstarts,dboutends = findBoutIndices(fish.dpix, fileloading.thresholdvalues[1], fileloading.thresholdframes[1])
		# The "True" indicates that the well center and rho data (and displacement) is already done, and those lists will return as empty
		dboutproperties, dsleepbouts = CalculateBoutProperties(fish.dpix, timestamp_data_array, dboutstarts, dboutends, fish.rho_array, fish.theta_array, fish.x_array, fish.y_array, "dpix_", True)
		
		# Processed data object, with access to binned_time((tuple that can be 2 or 1)), data itself, name, and then will add the graphing parameters to it (x-axis, y-axis), high-speed or slow-speed (default)
		# Fish contain a list of Processed data objects
		# EventSection contains a list of fish with truncated or full data sets (depends if we copy)?, and a fxn to sort them based on genotype and create the separate lists to iterate through (if it == 2 or 1 or more give warning about how it's not functional)
		
		# finding waking activity, or active sec / hr that are not inactive minutes
		fish.add_binned_data(sleep(fish_distances.copy(), fileloading.activitytimesthresholds[0], sleepbouts, indexdict["1"], indexdict["3600"]))
		fish.add_binned_data(sleep(fish.dpix.copy().astype(float), fileloading.activitytimesthresholds[1], dsleepbouts, indexdict["1"], indexdict["3600"], "dpix_"))
		for timebin in fileloading.boutbins:
			fish.add_binned_data(bout_flexactivity(boutproperties, boutstarts, indexdict[str(timebin)], str(timebin)))
			fish.add_binned_data(bout_flexactivity(dboutproperties, dboutstarts, indexdict[str(timebin)], str(timebin)))
		for r in range(0, len(fileloading.activitytimes)-1, 2):
			tup_time = (str(fileloading.activitytimes[r]),str(fileloading.activitytimes[r+1]))
			fish.add_binned_data(flexactivity(fish_distances, fileloading.activitytimesthresholds[0], indexdict[str(fileloading.activitytimes[r])],indexdict[str(fileloading.activitytimes[r+1])], tup_time))
			fish.add_binned_data(flexactivity(fish.dpix, fileloading.activitytimesthresholds[1], indexdict[str(fileloading.activitytimes[r])],indexdict[str(fileloading.activitytimes[r+1])], tup_time))
		for es2 in eventsectionlist:
			if es2.type != "time":
				fish.add_binned_data(CalculateEventProperties(es2.name, es2.events, fish.hs_dict, hs_fish_distances, fileloading.hsthresholdvalues[1], fileloading.hsthresholdframes[1], fish.hs_pos_x, fish.hs_pos_y))
	#pr.disable()
	#pr.print_stats(sort='time')
	return eventsectionlist, fish_list

# Start here
fileloading.initialize_args()
if( fileloading.graphonly ):
	graphsstatsandfilter.main(fileloading.graphparameters, fileloading.lightbaseline, fileloading.obendfilter, fileloading.cbendfilter)
else:
	eventsectionlist, fish_list = process_all_data()
	setupgraphsandsavedata.savedataandplot(eventsectionlist, fish_list)
	graphsstatsandfilter.main(fileloading.graphparameters, fileloading.lightbaseline, fileloading.obendfilter, fileloading.cbendfilter)
```
