## Supplemental Software for "A Customizable Low-Cost System for Massively Parallel Zebrafish Behavior Phenotyping": setupgraphsandsavedata.py

```
#!/usr/bin/python

import os,sys,glob,re
import numpy as np
import scipy
import datetime
import time
from datetime import timedelta
from scipy import stats
from scipy.stats import norm
from scipy.stats import mstats

import fileloading # prepares all the files

import pandas as pd
import statsmodels.api as sm

def savedata(genodict, iddict, name, xlabel, ylabel, t = None):
	ribgraphname = "ribgraph_mean_" + name
	for geno,data in genodict.items():
		idstr = '-'.join(iddict[geno])
		np.savetxt(ribgraphname + "_" + geno + ".data", np.array(data,dtype=np.float64), delimiter = ',', header=idstr)

def listtoNanarrayfreq(list):
	nparray = np.asarray(list,dtype=np.float32)
	#nparray[nparray==0.0]=np.nan
	return nparray

def listtoNanarray(list):
	nparray = np.asarray(list,dtype=np.float32)
	nparray[nparray==0.0]=np.nan
	return nparray

def savedataandplot(eventsectionlist, fish_list):
	# Sorting out the Fish objects based on genotype
	genotypes = set()
	for fish in fish_list:
		genotypes.add(fish.genogroup + "-" + fish.realgenotype)
	for es in eventsectionlist:
		graphtitlemid = es.type + "_" + es.name
		# Getting list of names of relevant binned data
		binnednames = set()
		for b in fish_list[0].binned_data:
			if(es.type == "time"): # standard time event
				if(b.slow_speed): # activity and bout graphs
					binnednames.add((b.name, b.time_bin))
			else: # high-speed event
				if(b.event_type == es.name):
					binnednames.add((b.name, b.time_bin))
		for bd in binnednames:
			splitfishlist = {}
			splitfishids = {}
			for g in genotypes:
				splitfishlist[g] = []
				splitfishids[g] = []
			bdname = ""
			for f in fish_list:
				splitfishids[f.genogroup + "-" + f.realgenotype].append(str(f.idnumber))
				for BD in f.binned_data:
					if (bd == (BD.name, BD.time_bin)):
						if((len(BD.time_bin) > 1) and BD.slow_speed): # activity (sleep) plots
							splitfishlist[f.genogroup + "-" + f.realgenotype].append(BD.binned_data[es.indexdict[BD.time_bin[1]][0]:es.indexdict[BD.time_bin[1]][1]])
							bdname = BD.name + "_" + str(BD.time_bin[0]) + "over" + str(BD.time_bin[1])
						elif((len(BD.time_bin) == 1) and BD.slow_speed): # bout graphs
							splitfishlist[f.genogroup + "-" + f.realgenotype].append(BD.binned_data[es.indexdict[BD.time_bin[0]][0]:es.indexdict[BD.time_bin[0]][1]])
							bdname = BD.name + "_" + str(BD.time_bin[0])
						else: # high-speed response plots
							if(BD.event_type == es.name):
								splitfishlist[f.genogroup + "-" + f.realgenotype].append(BD.binned_data)
								bdname = BD.name + "_" + str(BD.time_bin[0])
			savedata(splitfishlist, splitfishids, graphtitlemid + "_" + bdname, "X", "Y")
```
