## Supplemental Software for "A Customizable Low-Cost System for Massively Parallel Zebrafish Behavior Phenotyping": queuecheck_behavior.py

```
#!/usr/bin/python

import os,sys,glob

ids = []
# Movies for high-speed tracking
for m in glob.glob("hsmovie*avi"):
	ids.append(m.split(".")[0].split("_")[-1])
# Current name for long movies that aren't tracked in this
for m2 in glob.glob("testlog*avi"):
	longid = m2.split(".")[0].split("_")[-1]
	if longid in ids:
		ids.remove(longid)

completednumbers = []
for mfile in glob.glob('hsmovie*motion2'):
	statinfo = os.stat(mfile)
	if statinfo.st_size != 0:
		#print(statinfo.st_size)
		completednumbers.append(mfile.split('.')[0].split('_')[-1])
redo = [i for i in ids]
redo3 = list(set(redo).difference(completednumbers))
str1 = ""
for n in redo3:
	str1 = str1 + str(n) + ","
print (str1)
```
