## Supplemental Software for "A Customizable Low-Cost System for Massively Parallel Zebrafish Behavior Phenotyping": sort_pvalues.py

```
#!/usr/bin/python
import glob,argparse

parser = argparse.ArgumentParser(description='options for sorting p-values')
parser.add_argument('-statsfile', type=str, action="store", dest="statsfile", default="linearmodel*out")
parser.add_argument('-ofile', type=str, action="store", dest="ofile", default="")
parser.add_argument('-include', type=str, action="store", dest="includetxt", default="ribgraph")
parser.add_argument('-exclude', type=str, action="store", dest="excludetxt", default="weirdword")
parser.add_argument('-cutoff', type=float, action="store", dest="cutoff", default=0.05)
parser.add_argument('-overwriteanova', action="store_true", dest="overwriteanova", default=False) # Use the pval from linear mixed model instead of fram anova
parser.add_argument('-showfulldata', action="store_true", dest="showfulldata", default=False)
parser.add_argument('-sortalpha', action="store_true", dest="sortalpha", default=False)

args = parser.parse_args()
statsfile = args.statsfile
ofile = args.ofile
includetxt = args.includetxt
excludetxt = args.excludetxt
cutoff = args.cutoff
overwriteanova = args.overwriteanova
showfulldata = args.showfulldata
sortalpha = args.sortalpha # default will be to sort by lowest pval, but this is another option

for sfilename in glob.glob(statsfile):
	pvallist = []
	sfile = open(sfilename, 'r')
	if ofile == "":
		ofile = "pvals_" + sfilename
	outfile = open(ofile, 'w')
	lines = sfile.readlines()
	for line in lines:
		#anova:  ribgraph_mean_time_day2dfall_numberofbouts_3600_controlgroup-het.data : N of control, test, Mean of array control, test, Variance of array control, test, SSMD, H-stat, P-value:  43 35 1196.7162790697676 1193.72 272040.68322336394 316867.31017142854 0.0039044380458149457 0.10660666975064714 0.7440409692821573
		if line.startswith("anova:"): # automatically skipping all the failed ones
			pval = line.strip().split()[-1] # I always leave pval at the end
			graph = line.strip().split(":")[1].strip()
			fulldata = line.strip().split(":")[3].strip()
			if float(pval) < cutoff:
				pvallist.append([float(pval), graph, fulldata])
		else:
			if not overwriteanova: # don't even look at the linear mixed model info
				continue
			if line.startswith("ribgraph"):
				ribgraph = line.strip() # saving and then it will match up with the next instance of hitting the "mutornot"
			if line.startswith("mutornot[T.wt] "):
				if len(line.split()) > 3:
					lmmpval = line.split()[4]
					coef = line.split()[1]
					if float(lmmpval) == 0:
						lmmpval = 0.001
					if float(lmmpval) < cutoff:
						pvallist.append([float(lmmpval), ribgraph, ""])
	for stat in pvallist:
		if stat[2] == "": # if it's lmm data
			for s in range(0, len(pvallist)):
				if stat[1] == pvallist[s][1]: # found the lmm data
					pvallist[s][0] = stat[0] # replacing pval in the non-lmm data with lmm pval
	filteredpvallist = [x for x in pvallist if not x[2]==""] # eliminate all the original lmms
	if sortalpha:
		filteredpvallist.sort(key=lambda x: x[1])
	else:
		filteredpvallist.sort(key=lambda x: x[0])
	for final in filteredpvallist:
		if showfulldata:
			if includetxt in str(final[1]):
				if excludetxt not in str(final[1]):
					outfile.write(str(final[0]) + " " + str(final[1]) + str(final[2]) + '\n')
		else:
			if includetxt in str(final[1]):
				if excludetxt not in str(final[1]):
					outfile.write(str(final[0]) + " " + str(final[1]) + '\n')
# don't forget to count the ones <0.05
```
