## Supplemental Software for "A Customizable Low-Cost System for Massively Parallel Zebrafish Behavior Phenotyping": splitgenotypes.py

```
#!/usr/bin/python
import glob

alphatonum = {'A': 0, 'B': 12, 'C': 24, 'D': 36, 'E': 48, 'F': 60, 'G': 72, 'H': 84, 'I':96, 'J':108, 'K':120, 'L':132, 'M': 144, 'N':156, 'O':168, 'P':180, 'Q':192}
alphatonum_col = {'A': 0, 'B': 1, 'C': 2, 'D': 3, 'E': 4, 'F': 5, 'G': 6, 'H': 7, 'I':8, 'J':9, 'K':10, 'L':11, 'M': 12, 'N':13, 'O':14, 'P':15, 'Q':16}
alphatonum_colflip = {'A': 7, 'B': 6, 'C': 5, 'D': 4, 'E': 3, 'F': 2, 'G': 1, 'H': 0}

def Reverse(lst):
	return [ele for ele in reversed(lst)]

def zipper(a,b):
	list = [a[i] + b[i] for i in range(len(a))]
	return list

gfile = open("genotyping", 'w')
for file in glob.glob('*matrix*'):
	f = open(file, 'r')
	line1 = f.readline()
	lines = f.readlines()
	wells = {}
	for line in lines:
		if len(line.split()) != 13:
			print( "Either row not correct length or duplicated gene set, length = ", len(line.split()))
			if len(line.split()) != 26:
				print( "REAL ERROR REAL ERROR REAL ERROR, length = ", len(line.split()))
			wells[line.split()[0]] = zipper(line.split()[1:13], line.split()[14:26])
			# IF YOUR CAMERA IS FLIPPED AROUND, UNCOMMENT BELOW LINE AND COMMENT ABOVE LINE
			#wells[line.split()[0]] = Reverse(zipper(line.split()[1:13], line.split()[14:26]))
		else:
			wells[line.split()[0]] = line.split()[1:13]
			# IF YOUR CAMERA IS FLIPPED AROUND, UNCOMMENT BELOW LINE AND COMMENT ABOVE LINE
			#wells[line.split()[0]] = Reverse(line.split()[1:13])
	valuelist = []
	for v in wells.values():
		valuelist = valuelist + v
	valueset = set(valuelist)
	finaldata = {}
	for v2 in valueset:
		finaldata[v2] = []
	for a in wells.keys():
		for v3 in range(0, len(wells[a])):
			num = ((v3)*8)+1 + alphatonum_col[a]
			# IF YOUR CAMERA IS FLIPPED AROUND, UNCOMMENT BELOW LINE AND COMMENT ABOVE LINE
			#num = ((v3)*8)+1 + alphatonum_colflip[a]
			finaldata[wells[a][v3]].append(num)
	for k in finaldata.keys():
		finaldata[k].sort()
	print(file)
	print(finaldata)
	gfile.write(file)
	gfile.write('\n')
	for key in finaldata.keys():
		title = file.split('_')[0] + "_" + key
		gfile.write(title + ": ")
		gfile.write(str(finaldata[key]).strip().strip("[").strip("]").replace(" ", ""))
		gfile.write('\n')
```
