## Supplemental Software for "A Customizable Low-Cost System for Massively Parallel Zebrafish Behavior Phenotyping": track.slurm

```
#!/bin/bash
#SBATCH -p express # Partition to submit to
#SBATCH -n 1 # Number of cores requested
#SBATCH -N 1 # Ensure that all cores are on one machine
#SBATCH -t 15 # Runtime in minutes
#SBATCH --mem=8000 # Memory per cpu in MB (see also --mem-per-cpu)
#SBATCH --job-name=arrayJob
#SBATCH --array=147,277,278,408,669,670,31,419,682,171,48,693,188,572,574,337,214,87,600,219,229,360,238,370
#SBATCH -o hostname_%A_%a.out # Standard out goes to this file
#SBATCH -e hostname_%A_%a.err # Standard err goes to this filehostname

#module load Anaconda3/5.3.1

xvfb-run -a python highspeedmovieanalysis.py -r "newrois" -m "hsmovieThu, Aug 6, 2020_$SLURM_ARRAY_TASK_ID.avi" -e "fulltestrun_final_01_27_2020"
```
