## Supplemental Software for "A Customizable Low-Cost System for Massively Parallel Zebrafish Behavior Phenotyping": pdb.h

```
/* Audio Library for Teensy 3.X
 * Copyright (c) 2014, Paul Stoffregen,
 *
 * Development of this audio library was funded by PJRC.COM, LLC by sales of
 * Teensy and Audio Adaptor boards.  Please support PJRC's efforts to develop
 * open source software by purchasing Teensy or other PJRC products.
 *
 * Permission is hereby granted, free of charge, to any person obtaining a copy
 * of this software and associated documentation files (the "Software"), to deal
 * in the Software without restriction, including without limitation the rights
 * to use, copy, modify, merge, publish, distribute, sublicense, and/or sell
 * copies of the Software, and to permit persons to whom the Software is
 * furnished to do so, subject to the following conditions:
 *
 * The above copyright notice, development funding notice, and this permission
 * notice shall be included in all copies or substantial portions of the Software.
 *
 * THE SOFTWARE IS PROVIDED "AS IS", WITHOUT WARRANTY OF ANY KIND, EXPRESS OR
 * IMPLIED, INCLUDING BUT NOT LIMITED TO THE WARRANTIES OF MERCHANTABILITY,
 * FITNESS FOR A PARTICULAR PURPOSE AND NONINFRINGEMENT. IN NO EVENT SHALL THE
 * AUTHORS OR COPYRIGHT HOLDERS BE LIABLE FOR ANY CLAIM, DAMAGES OR OTHER
 * LIABILITY, WHETHER IN AN ACTION OF CONTRACT, TORT OR OTHERWISE, ARISING FROM,
 * OUT OF OR IN CONNECTION WITH THE SOFTWARE OR THE USE OR OTHER DEALINGS IN
 * THE SOFTWARE.
 */

#ifndef pdb_h_
#define pdb_h_

#include "kinetis.h"

// Multiple input & output objects use the Programmable Delay Block
// to set their sample rate.  They must all configure the same
// period to avoid chaos.

#define PDB_CONFIG (PDB_SC_TRGSEL(15) | PDB_SC_PDBEN | PDB_SC_CONT | PDB_SC_PDBIE | PDB_SC_DMAEN)


#if F_BUS == 120000000
  #define PDB_PERIOD (2720-1)
#elif F_BUS == 108000000
  #define PDB_PERIOD (2448-1)
#elif F_BUS == 96000000
  #define PDB_PERIOD (2176-1)
#elif F_BUS == 90000000
  #define PDB_PERIOD (2040-1)
#elif F_BUS == 80000000
  #define PDB_PERIOD (1813-1)  // small ?? error
#elif F_BUS == 72000000
  #define PDB_PERIOD (1632-1)
#elif F_BUS == 64000000
  #define PDB_PERIOD (1451-1)  // small ?? error
#elif F_BUS == 60000000
  #define PDB_PERIOD (1360-1)
#elif F_BUS == 56000000
  #define PDB_PERIOD (1269-1)  // 0.026% error
#elif F_BUS == 54000000
  #define PDB_PERIOD (1224-1)
#elif F_BUS == 48000000
  #define PDB_PERIOD (1088-1)
#elif F_BUS == 40000000
  #define PDB_PERIOD (907-1)  // small ?? error
#elif F_BUS == 36000000
  #define PDB_PERIOD (816-1)
#elif F_BUS == 24000000
  #define PDB_PERIOD (544-1)
#elif F_BUS == 16000000
  #define PDB_PERIOD (363-1)  // 0.092% error
#else
  #error "Unsupported F_BUS speed"
#endif

#endif
```
