## Supplemental Software for "A Customizable Low-Cost System for Massively Parallel Zebrafish Behavior Phenotyping": teensy36_dma_fish_box.ino

```
// extra triggers
#define T1 7
#define T2 8
#define T3 14  // also A0
#define T4 21  // also A7

#define CAL_AIN A0
#define CAL_VIS_MIN 0
#define CAL_VIS_STEP 5
#define CAL_VIS_N 100

#define N_STEP_SAMPLES 100
#define STEP_SAMPLE_DELAY 5  // ms

//#define CAL_VIS_MAX 500  // max vis duty to test during calibration
//#define CAL_VIS_DELAY 300  // ms
//#define CAL_VIS_N (CAL_VIS_MAX - CAL_VIS_MIN) / CAL_VIS_STEP

#define MIN_BRIGHTNESS 500  // photodiode adc units
#define MAX_RISE_TIME 10  // ms
#define MAX_FALL_TIME 10  // ms
//#define CAL_ACCEPT_THRESHOLD 0.90  // get with 90% of target intensity

#include <EEPROM.h>
#define CAL_VIS_ADDR 0

int vis_cal[CAL_VIS_N];
int min_vis_brightness = -1;

int step_samples[N_STEP_SAMPLES];

int triggers[] = {T1, T2, T3, T4};
int analog_inputs[] {-1, -1, 0, 7};

#define VISPIN 3
#define MUTEPIN 4

//#define VISFREQ 3662.109
//#define VISFREQ 10000
#define VISFREQ 6000
#define VIS_MAX_DUTY 600  // should give just a bit more than 300 mA

#define MAX9744_I2CADDR 0x4B

#include <Wire.h>
#include <DMAChannel.h>
#include "pdb.h"

DMAChannel dma(false);

// 128
#define N_SAMPLES 128
static volatile uint16_t sinetable[] = {
   2047,    2147,    2248,    2348,    2447,    2545,    2642,    2737,
   2831,    2923,    3012,    3100,    3185,    3267,    3346,    3422,
   3495,    3564,    3630,    3692,    3750,    3804,    3853,    3898,
   3939,    3975,    4007,    4034,    4056,    4073,    4085,    4093,
   4095,    4093,    4085,    4073,    4056,    4034,    4007,    3975,
   3939,    3898,    3853,    3804,    3750,    3692,    3630,    3564,
   3495,    3422,    3346,    3267,    3185,    3100,    3012,    2923,
   2831,    2737,    2642,    2545,    2447,    2348,    2248,    2147,
   2047,    1948,    1847,    1747,    1648,    1550,    1453,    1358,
   1264,    1172,    1083,     995,     910,     828,     749,     673,
    600,     531,     465,     403,     345,     291,     242,     197,
    156,     120,      88,      61,      39,      22,      10,       2,
      0,       2,      10,      22,      39,      61,      88,     120,
    156,     197,     242,     291,     345,     403,     465,     531,
    600,     673,     749,     828,     910,     995,    1083,    1172,
   1264,    1358,    1453,    1550,    1648,    1747,    1847,    1948,
};

static volatile uint16_t squaretable[] = {
0,    0,    0,    0,    0,    0,    0,    0,    0,    0,    0,
0,    0,    0,    0,    0,    0,    0,    0,    0,    0,    0,
0,    0,    0,    0,    0,    0,    0,    0,    0,    0,    0,
0,    0,    0,    0,    0,    0,    0,    0,    0,    0,    0,
0,    0,    0,    0,    0,    0,    0,    0,    0,    0,    0,
0,    0,    0,    0,    0,    0,    0,    0,    0, 4095, 4095,
4095, 4095, 4095, 4095, 4095, 4095, 4095, 4095, 4095, 4095, 4095,
4095, 4095, 4095, 4095, 4095, 4095, 4095, 4095, 4095, 4095, 4095,
4095, 4095, 4095, 4095, 4095, 4095, 4095, 4095, 4095, 4095, 4095,
4095, 4095, 4095, 4095, 4095, 4095, 4095, 4095, 4095, 4095, 4095,
4095, 4095, 4095, 4095, 4095, 4095, 4095, 4095, 4095, 4095, 4095,
4095, 4095, 4095, 4095, 4095, 4095, 4095,
};

volatile uint16_t *waveform = sinetable;

static volatile uint16_t dmatable[N_SAMPLES];

bool playing = false;
int freq = 900;
float amp = 0.0;
int waveform_index = 0;
unsigned long dur = 2;  // ms
elapsedMillis note_timer;
//bool enable_note_timer = false;

elapsedMillis delay_timer;
bool enable_delay = false;
int delay_ms = 0;


void mute() {
  digitalWrite(MUTEPIN, LOW);
}

void unmute() {
  digitalWrite(MUTEPIN, HIGH);
}

void setup_sound() {
  for (int16_t i=0; i<N_SAMPLES; i+=1) {
    dmatable[i] = 2048;
  }
  dma.begin(true); // allocate the DMA channel first
  
  SIM_SCGC2 |= SIM_SCGC2_DAC1; // enable DAC clock
  DAC1_C0 = DAC_C0_DACEN | DAC_C0_DACRFS; // enable the DAC module, 3.3V reference
  // slowly ramp up to DC voltage, approx 1/4 second
  for (int16_t i=0; i<2048; i+=8) {
    *(int16_t *)&(DAC1_DAT0L) = i;
    delay(1);
  }
  
  // set the programmable delay block to trigger DMA requests
  SIM_SCGC6 |= SIM_SCGC6_PDB; // enable PDB clock
  PDB0_IDLY = 0; // interrupt delay register
  //PDB0_MOD = PDB_PERIOD; // modulus register, sets period
  PDB0_MOD = 375;  // 375 gives 1 kHz: 48M / 128K (128K = 128 samples * 1000 times / sec)
  PDB0_CH0C1 = 0x0101; // channel n control register? disable adc pre-trigger?
  PDB0_SC = PDB_CONFIG | PDB_SC_LDOK; // load registers from buffers
  PDB0_SC = PDB_CONFIG | PDB_SC_SWTRIG; // reset and restart
  
  dma.sourceBuffer(dmatable, sizeof(dmatable));
  dma.destination(*(volatile uint16_t *)&(DAC1_DAT0L));
  dma.triggerAtHardwareEvent(DMAMUX_SOURCE_PDB);
}

void change_frequency(float hz) {
  int period = F_BUS / (N_SAMPLES * hz);
  PDB0_MOD = period;
  PDB0_SC = PDB_CONFIG | PDB_SC_LDOK;
}

void set_waveform(int i) {
  switch (i) {
    case 1:
      Serial.println("Set waveform to square");
      waveform = squaretable;
      break;
    case 0:
    default:
      Serial.println("Set waveform to sine");
      waveform = sinetable;
      break;
  }
}

void decay_sound(float decay_value) {
  for (int16_t i=0; i<N_SAMPLES; i+=1) {
    dmatable[i] = 2048 + (waveform[i] - 2048.) * decay_value;
  }
}

void start_playing() {
  dma.enable();
  playing = true;
  note_timer = 0;
}

void stop_playing() {
  dma.disable();
  playing = false;
}

void note_on(float amp, int freq) {
  change_frequency(freq);
  decay_sound(amp);
  start_playing();
  Serial.print("amp: "); Serial.println(amp);
  Serial.print("freq: "); Serial.println(freq);
  Serial.print("dur: "); Serial.println(dur);
  Serial.print("waveform: "); Serial.println(waveform_index);
}

void note_on(float amp) {
  note_on(amp, freq);
}

void note_on() {
  //Serial.println(DAC1_DAT0L);
  unmute();
  note_on(amp, freq);
}

void note_off() {
  stop_playing();
  mute();
  *(int16_t *)&(DAC1_DAT0L) = 2048;
  return;
  // slowly bring dac output back to 2048
  int v = *(int16_t *)&(DAC1_DAT0L);
  for (int i=v; i > 2048; i--) {
    *(int16_t *)&(DAC1_DAT0L) = i;
    delay(1);
  }
  for (int i=v; i < 2048; i++) {
    *(int16_t *)&(DAC1_DAT0L) = i;
    delay(1);
  }
  Serial.println(*(int16_t *)&(DAC1_DAT0L));
}


void play_note() {
  note_on();
  while(note_timer < dur);
  note_off();
}

void set_volume(int vol) {
  if (vol > 63) vol = 63;
  if (vol < 0) vol = 0;
  Wire.beginTransmission(MAX9744_I2CADDR);
  Wire.write(vol);  // vol from 0 to 63 inclusive
  if (Wire.endTransmission() != 0) {
    Serial.println("Error volume not set");
  }
}


void zero_calibration() {
  for (int i=0; i<CAL_VIS_N; i++) {
    vis_cal[i] = -1;
  }
}

void save_calibration() {
  for (int i=0; i<CAL_VIS_N; i++) {
    int addr = CAL_VIS_ADDR + i * 2;
    byte hb = (vis_cal[i] >> 8);
    byte lb = (vis_cal[i] & 0xFF);
    // Serial.print(lb); Serial.print(",");
    // Serial.println(hb);
    EEPROM.write(addr, hb);
    EEPROM.write(addr + 1, lb);
  }
}

void load_calibration() {
  min_vis_brightness = -1;
  for (int i=0; i<CAL_VIS_N; i++) {
    int addr = CAL_VIS_ADDR + i * 2;
    byte hb = EEPROM.read(addr);
    byte lb = EEPROM.read(addr + 1);
    // Serial.print(lb); Serial.print(",");
    // Serial.println(hb);
    vis_cal[i] = (((int)hb) << 8) | lb;
    if (vis_cal[i] == 0xFFFF) vis_cal[i] = -1;
    if ((vis_cal[i] != -1) && (min_vis_brightness == -1)) {
      min_vis_brightness = vis_cal[i];
    }
  }
}

void report_calibration() {
  for (int i=0; i<CAL_VIS_N; i++) {
    Serial.println(vis_cal[i]);
  }
}

void setup() {
  Serial.begin(9600);  
  //pinMode(IRPIN, OUTPUT);
  pinMode(VISPIN, OUTPUT);
  //digitalWrite(IRPIN, LOW);
  digitalWrite(VISPIN, LOW);
  analogWriteResolution(10);
  analogWriteFrequency(VISPIN, VISFREQ);
  //Serial.println(*(int16_t *)&(DAC1_DAT0L));
  pinMode(MUTEPIN, OUTPUT);
  mute();
  
  // set digital volume to max
  Wire.begin();
  set_volume(63);
  
  setup_sound();
  //Serial.println(*(int16_t *)&(DAC1_DAT0L));
  load_calibration();
}

void sweep_freq() {
  // hz: 5000, 2500, 1000, 500, 250, 100, 50
  int fs[] = {75, 150, 375, 750, 1500, 3750, 7500};
  for (int i=0; i<7; i+=1) {
    change_frequency(fs[i]);
    // do nothing here
    start_playing();
    for (int i=0; i<100; i+=1) {
      decay_sound(sin(i / 100. * PI / 2.) * 0.1);
      delay(1);
    }
    delay(100);
    for (int i=100; i>0; i-=1) {
      decay_sound(sin(i / 100. * PI / 2.) * 0.1);
      delay(1);
    }
    stop_playing();
    delay(1000);
  }
}

void set_vis_duty(int duty) {
  if (duty > VIS_MAX_DUTY) duty = VIS_MAX_DUTY;
  analogWrite(VISPIN, duty);
}

void read_trigger() {
  byte i = Serial.read() - 49;  // convert '1' - '4' to 0-3
  if (i > 3) {
    Serial.print("Invalid trigger: ");
    Serial.println(i);
    return;
  }
  char s = Serial.read();
  int t = triggers[i];
  switch (s) {
    case 'u':
      pinMode(t, INPUT_PULLUP);
      break;
    case 'i':
      pinMode(t, INPUT);
      break;
    case 'd':
      pinMode(t, INPUT_PULLDOWN);
      break;
    case 'D':
      pinMode(t, INPUT_DISABLE);
      break;
    case 'o':
      pinMode(t, OUTPUT);
      break;
    case 'h':
      digitalWrite(t, HIGH);
      break;
    case 'l':
      digitalWrite(t, LOW);
      break;
    case 'r':
      Serial.print("t");
      Serial.print(i + 1);
      Serial.println(digitalRead(t));
      break;
    case 'a':
      t = analog_inputs[i];
      if (t == -1) {
        Serial.println("Not a valid analog pin");
      } else {
        Serial.print("t");
        Serial.print(i+1);
        Serial.println(analogRead(t));
      }
      break;
  }
}

void step_light(int step_intensity) {
  pinMode(CAL_AIN, INPUT_DISABLE);
  set_vis_duty(0);
  delay(100);
  set_vis_duty(step_intensity);
  for (int i=0; i<N_STEP_SAMPLES; i++) {
    step_samples[i] = analogRead(CAL_AIN);
    delay(STEP_SAMPLE_DELAY);
  }
  delay(100);
  set_vis_duty(0);
  for (int i=0; i<N_STEP_SAMPLES; i++) {
    Serial.print(i * STEP_SAMPLE_DELAY);
    Serial.print(",");
    Serial.println(step_samples[i]);
  }
}

int measure_rising_step(int step_intensity) {
  // assumes start is 0
  int i = 0;
  set_vis_duty(step_intensity);
  for (i=0; i<N_STEP_SAMPLES; i++) {
    step_samples[i] = analogRead(CAL_AIN);
    delay(STEP_SAMPLE_DELAY);
  }
  // process rising edge
  int target = step_samples[N_STEP_SAMPLES-1];
  int thresh = round(target * 0.632);  // TODO make configurable
  for (i=N_STEP_SAMPLES-2; i>=0; i--) {
    if (step_samples[i] < thresh) break;
  }
  if (i < 0) {
    return STEP_SAMPLE_DELAY * N_STEP_SAMPLES;
  } else {
    return i * STEP_SAMPLE_DELAY;
  }
}

int measure_falling_step(int step_intensity) {
  set_vis_duty(step_intensity);
  int i = 0;
  for (i=0; i<N_STEP_SAMPLES; i++) {
    step_samples[i] = analogRead(CAL_AIN);
    delay(STEP_SAMPLE_DELAY);
  }
  // process falling edge
  int target = step_samples[0];
  int thresh = round(target * 0.368);
  for (i=0; i<N_STEP_SAMPLES; i++) {
    if (step_samples[i] < thresh) break;
  }
  return i * STEP_SAMPLE_DELAY;
}

void calibrate_light() {
  pinMode(CAL_AIN, INPUT_DISABLE);
  set_vis_duty(0);
  delay(N_STEP_SAMPLES * STEP_SAMPLE_DELAY);
  // step through duty cycles
  //for (int duty=CAL_VIS_MIN; duty<=CAL_VIS_MAX; duty += CAL_VIS_STEP) {
  bool hit_min_brightness = false;
  bool always_increasing = true;
  int pv = -1;
  for (int i=0; i<CAL_VIS_N; i++) {
    int duty = CAL_VIS_MIN + CAL_VIS_STEP * i;
    int rt = measure_rising_step(duty);
    int v = analogRead(CAL_AIN);
    int ft = measure_falling_step(0);
    if ((rt > MAX_RISE_TIME) || (ft > MAX_FALL_TIME)) {
      vis_cal[i] = -1;
    } else {
      vis_cal[i] = v;
      if (v > MIN_BRIGHTNESS) hit_min_brightness = true;
      if (v < pv) always_increasing = false;
      pv = v;
    }
    /*
    set_vis_duty(duty);
    delay(CAL_VIS_DELAY);
    int v = analogRead(CAL_AIN);
    */
    Serial.print(duty); Serial.print(",");
    Serial.print(rt); Serial.print(",");
    Serial.print(ft); Serial.print(",");
    Serial.println(v);
  }

  // check that a minimum brightness was achieved
  if (!hit_min_brightness) {
    Serial.print("Calibration failed to reach minimum brightness[");
    Serial.print(MIN_BRIGHTNESS);
    Serial.println("]");
    zero_calibration();
  }
  if (!always_increasing) {
    Serial.print("Calibration failed: intensity decreased during steps");
    zero_calibration();
  }
  // clean up calibration
  // zero all valid points before invalid points
  for (int i=CAL_VIS_N-1; i>=0; i--) {
    if (vis_cal[i] == -1) {
      for (int j=0; j<i; j++) {
        vis_cal[j] = -1;
      }
      break;
    }
  }
  for (int i=0; i<CAL_VIS_N; i++) {
    if (vis_cal[i] != -1) {
      min_vis_brightness = vis_cal[i];
      break;
    }
  }
  // reset things back to normal
  set_vis_duty(0);
}

bool set_vis_brightness(int target) {
  if (target == 0) {
    set_vis_duty(0);
    return true;
  }
  //Serial.print("svb: "); Serial.println(target);
  for (int i=1; i<CAL_VIS_N; i++) {
    if (vis_cal[i] == -1) continue;  // invalid calibration point
    if (vis_cal[i] == target) {
      int duty = CAL_VIS_MIN + CAL_VIS_STEP * i;
      //Serial.print(" at point: "); Serial.println(duty);
      set_vis_duty(duty);
      return true;
    }
    if (vis_cal[i] > target) {
      
      // linearly interp between i and i-1
      int d0 = CAL_VIS_MIN + CAL_VIS_STEP * (i - 1);  // y
      int d1 = d0 + CAL_VIS_STEP;
      int v0 = vis_cal[i-1];  // x
      if (v0 == -1) return false;
      int v1 = vis_cal[i];
      float m = (d1 - d0) / ((float)(v1 - v0));
      float b = d1 - m * v1;
      int duty = round(m * target + b);
      /*
      Serial.print(" interp: ");
      Serial.print(d0); Serial.print(",");
      Serial.print(d1); Serial.print(",");
      Serial.print(v0); Serial.print(",");
      Serial.print(v1); Serial.print(",");
      Serial.print(m); Serial.print(",");
      Serial.print(b); Serial.print(",");
      Serial.println(duty);
      */
      if ((duty < 0) || (duty > VIS_MAX_DUTY)) return false;
      set_vis_duty(duty);
      return true;
    }
  }
  return false;
}

bool test_calibration() {
  pinMode(CAL_AIN, INPUT_DISABLE);
  set_vis_duty(0);
  delay(MAX_FALL_TIME);
  for (int i=0; i<CAL_VIS_N; i++) {
    int target = (i * MIN_BRIGHTNESS) / (CAL_VIS_N - 1);
    if (target < min_vis_brightness) {
      Serial.print(target); Serial.print(",");
      Serial.println(-1);
      continue;
    }
    if (!set_vis_brightness(target)) {
      Serial.print("Failed to set brightness: ");
      Serial.println(target);
      return false;
    }
    delay(N_STEP_SAMPLES * STEP_SAMPLE_DELAY);
    int v = analogRead(CAL_AIN);
    set_vis_duty(0);
    delay(N_STEP_SAMPLES * STEP_SAMPLE_DELAY);
    float dev = 1. - abs(1. - (v / ((float)target)));
    /*
    if (1. - abs(1. - (v / (float)target)) < CAL_ACCEPT_THRESHOLD) {
      Serial.print("Failed to reach brightness: ");
      Serial.print(v);
      Serial.print(" != ");
      Serial.println(target);
      return false;
    }
    */
    Serial.print(target); Serial.print(",");
    Serial.print(v); Serial.print(",");
    Serial.println(dev);
  }
  set_vis_duty(0);
  return true;
}

void loop() {
  if (Serial.available()) {
    char c = Serial.read();
    switch (c) {
      // lights
      case 'v':
        set_vis_duty(Serial.parseInt());
        break;
      case 'b':  // set to a particular brightness using the calibration
        // print 1 if brightness set, 0 if brightness failed
        Serial.println(set_vis_brightness(Serial.parseInt()));
        break;
      // sound
      case 'f':
        freq = Serial.parseInt();
        change_frequency(freq);
        break;
      case 'a':
        amp = Serial.parseFloat();
        break;
      case 'd':
        dur = Serial.parseInt();
        break;
      case 'D':
        delay_ms = Serial.parseInt();
        //enable_delay = true;
        delay(delay_ms);
        break;
      case 'p':
        //note_on();
        play_note();
        break;
      case 'w':
        waveform_index = Serial.parseInt();
        set_waveform(waveform_index);
        break;
      case 'g':
        set_volume(Serial.parseInt());
        break;
      // triggers
      case 't':
        read_trigger();
        break;
      // calibration
      case 's':
        step_light(Serial.parseInt());
        break;
      case 'c':
        calibrate_light();
        break;
      case 'T':
        test_calibration();
        break;
      case 'S':
        save_calibration();
        break;
      case 'L':
        load_calibration();
        break;
      case 'R':
        report_calibration();
        break;

    }
  }
  /*
  if (playing & (note_timer > dur)) {
    note_off();
  }
  */
}
```
