## Supplementary Material for "A Customizable Low-Cost System for Massively Parallel Zebrafish Behavior Phenotyping"

#### Assembly Instructions

All parts are listed in the BillofMaterials spreadsheet. Parts in multi-packs can be used for multiple builds, and several types of screws/nuts/washers can be substituted with alternatives. FabricationFiles.zip includes technical drawings for custom parts: DXF files for 2D manufacturing and STL files for 3D printing. If a laser cutter and 3D printer are locally unavailable, the following online fabricators can produce these components:

**For 2D manufacturing**
- https://www.emachineshop.com/
- https://www.bigbluesaw.com/
- https://www.protolabs.com/

**For 3D printing**
- https://www.protolabs.com/
- https://www.3dhubs.com/
- https://www.shapeways.com/
- https://www.sculpteo.com/en/

**For PCB manufacturing**
- https://www.sunstone.com/
- https://oshpark.com/
- https://www.4pcb.com/

##### Enclosure

1) Laser-cut all necessary pieces (**Supplementary Figure 1A-B**).
2) Attach the 80/20 rails (Part #1) to the aluminum bottom plate (**Supplementary Figure 1A**) with ¼″-20 threaded socket head screws (Part #2).
3) Use T-slot screw and nut (Part #3) to attach shelf supports facing inwards (Part #4) 11 inches from the bottom of each of the four 80/20 rails (Part #1) (**Supplementary Figure 1C**, **Supplementary Figure 2A-B**).

   ***Warning**: Shelf support position can be adjusted by slightly untightening the T-slot screw and nut set, but the screw can easily detach from the nut, allowing it to fall to the bottom of the rail. Multiple pieces may need to be dissassembled to recover the nut.
4) Secure the HDPE panels in **Supplementary Figure 1B** to the 80/20 rails by loosely affixing T-slot screws and nuts to each hole in the panel and sliding the panels into their respective positions. Tighten screws as necessary.
5) Attach the top aluminum panel (**Supplementary Figure 1A**) to the enclosure using socket head screws (Part #2).

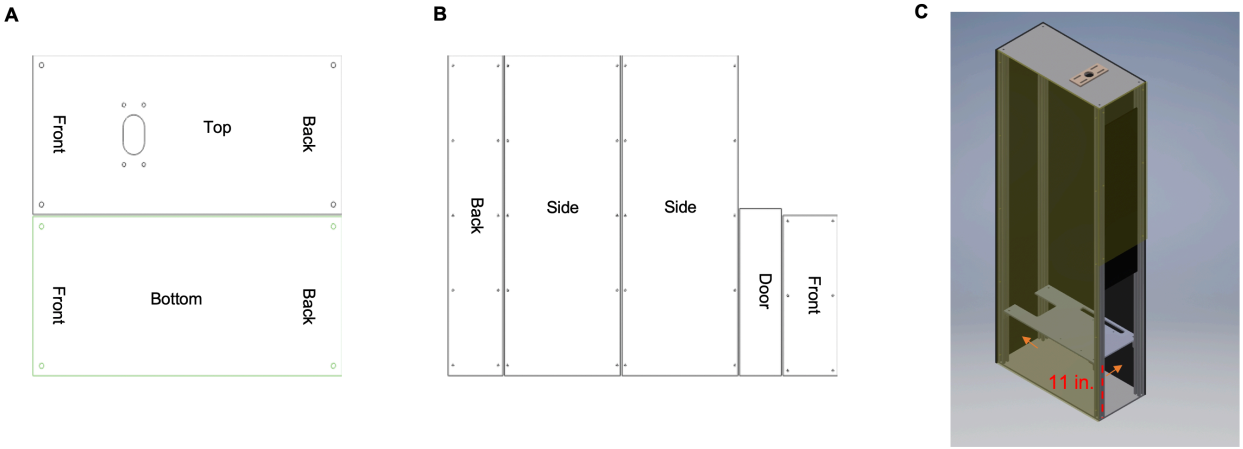

***Supplementary Figure 1. Box Enclosure Components (A)*** *2D drawing (Enclosure Top Bottom.dxf) of the top and bottom aluminum plates. The four holes located at the corners of the plate are for attaching socket head screws (Part #2). The four holes near the front of the top plate are for attaching the camera mount (alternative camera mounts with different hole patterns are also included and discussed in Camera Mounting).* ***(B)*** *2D drawing (Enclosure Walls.dxf) of the enclosure panels with corresponding holes for T-slot screws. The panel without holes is a sliding door at the front of the box* ***(C)*** *A 3D rendering of the enclosure. Orange arrows refer to approximate hole placement for tubing and wires (precise positioning not required). The arrow pointing at the back of the box is for water tubing input and light power cords. The arrow pointing at the right of the box is for tubing water output and wires for light and sound control.*

6) Secure fish chamber shelf (**Supplementary Figure 2**) to the supports (Part #4) using socket head screws (Part #5), hex nuts (Part #6), and washers (Part #7). Confirm that shelf is level and adjust supports if necessary.
7) Drill a ~1-inch hole on the back panel and side panel for water tubing and wires (**Supplementary Figure 1C**).
8) Seal box seams with black masking tape to eliminate external light.
9) Assemble light shelf (**Supplementary Figure 3**) and add to the box. The white LED panel (Part #8) must be wired to electronics before adding (see Electronics Videos).

    ***Warning**: If the suggested LED panel is out of stock, substitute light panels with appropriate wattage, lumens, and shape. Wattage/lumens differences may require additional system optimization, while size/shape (11.7 x 9.3 x 1.5 inches) differences may require modification of the shelf design.

10) Infra-red light (Part #9) may flicker when viewed through the camera. To prevent flickering, cover the photocell (round sensor on IR light housing) with black masking tape.

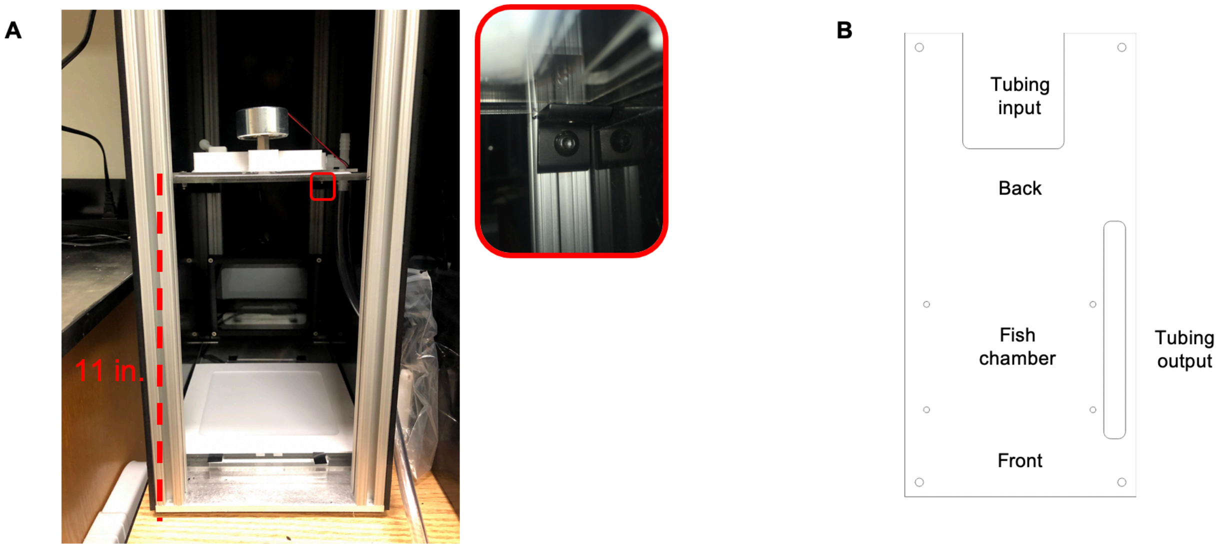

***Supplementary Figure 2. Box Interior Components. (A)*** *Interior of the setup. The fish chamber shelf is located 11 inches from the bottom. Shelf supports are shown attached to the rails (red inset). Light shelf rests on bottom with both white light panel and IR light (Part #9) (**Supplementary Figure 3**).* ***(B)*** *2D drawing (Fish Chamber Shelf.dxf) of fish chamber shelf oriented for placement in box. The four corners of the shelf are for securing to the shelf supports (Part #4), while the four smaller holes for securing the fish chamber (see Fish Chamber Setup). The slot to the right is for the output tubing nozzle and wires for the surface transducer.*

#### Light Setup

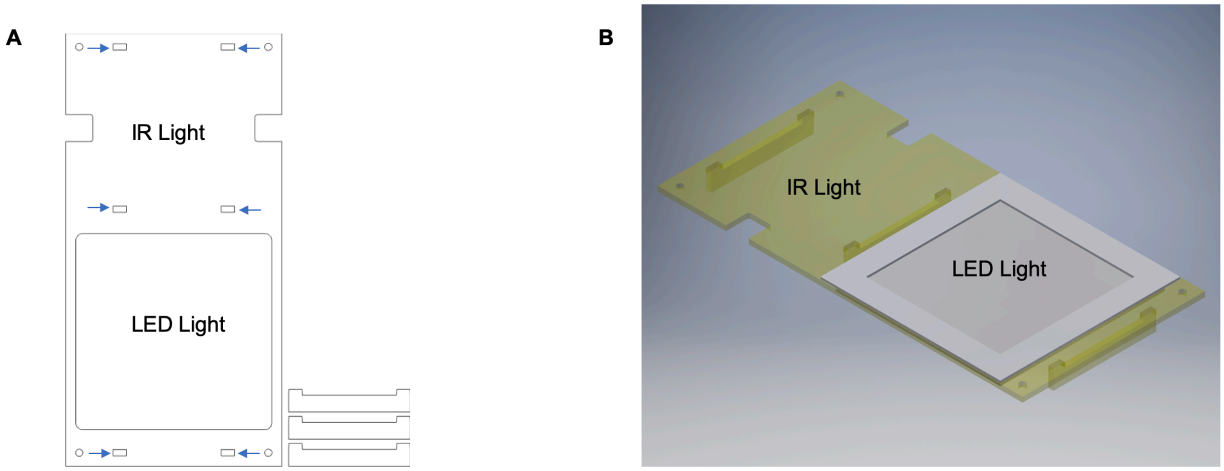

***Supplementary Figure 3 Assembling Light Shelf (A****)* *The acrylic shelf (Light Shelf.dxf) for the LED light (Part #8) includes three acrylic legs (CAD file). Insert legs to the specified slots (blue arrows) to elevate the shelf from the bottom of the box. Depending on the accuracy of the laser cutter, you may need to glue or tape the legs to the plate.* ***(B)*** *The LED light rests on the shelf, with wires fed underneath the shelf and through the back of the box. Cut and strip the power cord to connect to the teensy electronics board (see Electronics Videos). Place the light shelf in the bottom of the box and orient the IR light (Part #9) to face slightly down and toward the front of the box. The IR light reflects off the surface of the LED light. Also see Supplementary Figure 2A light placement.*

#### Camera Mounting

1) If acrylic is transparent, wrap in black masking tape to prevent light from entering the box.
2) Attach acrylic camera mount to the aluminum plate using M5 screws (Part #10), corresponding hex nuts (Part #11), and washers (Part #12) along with the corner bracket (Part #13). The corner bracket was modified to accommodate M5 screws (see Notes 1 and 2 below).
3) Assemble the camera: carefully cut out a round piece of the IR filter (Part #14) and place between the 50mm lens (Part #15) and the Grasshopper3 camera (Part #16). Attach the mount that comes with camera.
4) Secure camera with lens pointing into box, and tighten the camera to the corner bracket using a socket head screw (Part #17) (**Supplementary Figure 4**). Temporarily loosen this screw when adjusting lens focus. Camera orientation will affect downstream data analysis.
5) Connect camera to computer using USB 3.1 locking cable (Part #18). Tighten locks to prevent disruptions in data acquisition.

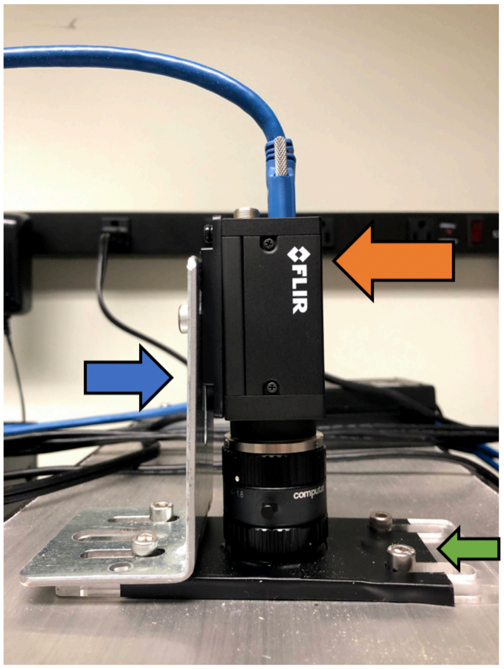

***Supplementary Figure 4. Camera Mounting*** *Blue arrow: corner bracket. In our setup, two additional slots were added to the bracket to match the mount and the aluminum top. Green arrow: camera mount. Orange arrow: Grasshopper 3 high speed camera. To prevent damage to the USB port, avoid straining/bending the cable.*

*Note 1: We found that M5 screws fit perfectly in the laser-cut holes of the aluminum panel, but results may vary between laser cutters. Alternatively, holes can be bored with a ¼″-20 tap to create threads for ¼″-20 screws.

*Note 2: In initial versions of our setup, holes at the top of the box accomodated double slots in the camera mount and bracket. However, the corner bracket (Part #13) that stabilizes the camera was modified by a machine shop to match this setup. For future iterations, we created 2D DXF files (Camera Mount Single Slot.dxf, Enclosure Top Bottom Single Slot.dxf) to accommodate single slots in the camera mount bracket. M5 screws (Part #10) can secure the bracket, but the single slot version requires wider washers than those listed in the BillofMaterials.xlsx.

#### Fish Chamber Setup

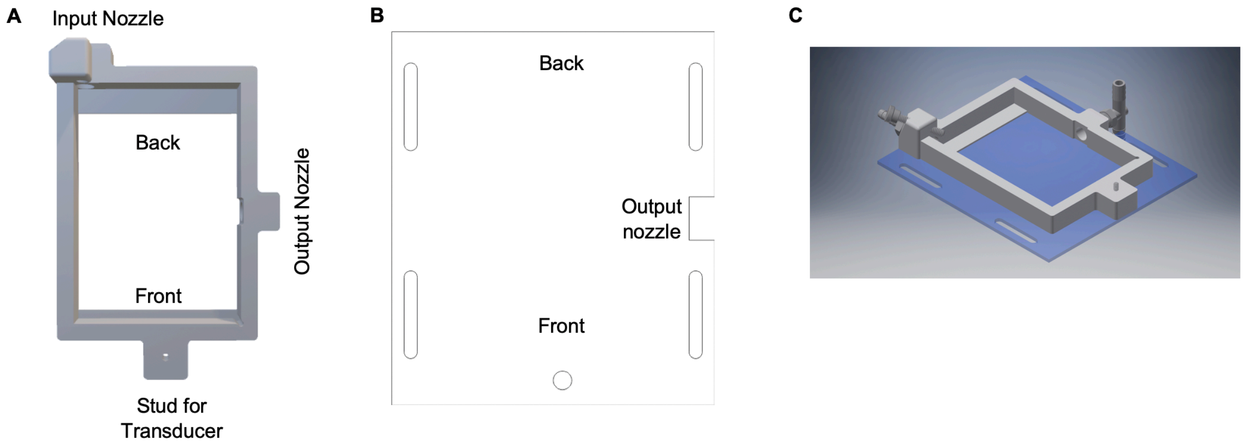

***Supplementary Figure 5. Fish Chamber Assembly.*** **(A)** The 3D printed fish chamber (Fish Chamber Frame.stl). The lip towards the back of the chamber raises the fish plate slightly to allow water flow under and around the plate. Nylon screw (Part #19) is attached to the top left corner and secures the fish plate, preventing movement when the surface transducer vibrates. Holes are tapped for all attachments (¼″ NPT tap for the output and input nozzles, ¼″-20 tap for the nylon screw, and M5 tap for the socket head screw). **(B)** 2D drawing (Fish Chamber Floor.dxf) of the fish chamber floor. Slots on the floor plate allow position adjustments relative to the fish chamber shelf, to align with the camera. **(C)** 3D rendering of the assembled fish chamber.

1) Tap threads (**Supplementary Figure 5A**) for the nylon screw that secures the fish plate (Part #19), the tubing input/output nozzles (Part #20, Part #21), and the socket head screw (Part #22) for the surface transducer (Part #52). Taps are listed under **Useful Tools** in BillofMaterials.
2) Attach tubing input and output (Parts #20 and #21) and wrap the threads with plumbing tape.
3) Screw in the M5 socket head screw (Part #22) to the plate with the end pointing up. Next, screw on the hex standoff (Part #23) for the surface transducer. *Note: the surface transducer (Part #52) will screw onto the M5 stud with standoff, but must first be connected to the electronics board.
4) Secure the fish chamber to the fish chamber floor (**Supplementary Figure 5B, Supplementary Figure 5C**) using acrylic adhesive. Ensure proper sealing to prevent leaks. However, too much adhesive could mar the acrylic surface and decrease visibility and tracking ability. Follow manufacturer's instructions regarding adhesive drying/setting.
5) Seal edges on both the inside and outside with silicone sealant. Avoid marring surface of acrylic.

6) Use the M5 threaded 22 mm long socket head screws (Part #10), hex nuts (Part #11), and washers (Part #12) to screw in fish chamber floor to the chamber shelf (**Supplementary Figure 2B**). Position can be adjusted as necessary to align with camera.

#### Heat Shock Assay Setup

The heat shock assay is an optional component and requires a three-way valve, input/output tubing, water pump, and tub (See **Supplementary Figure 6A** for water flow diagram).

1) Attach the water pump to the tub (Parts #24 and #25). Attach the pump power cord (Part #26).
2) Length of the inner tubing (Part #27) should be distance from input nozzle to tub + at least 2 extra inches. Equivalent for output tubing / output nozzle (Part #28).
3) Cut tubing to respective lengths.
4) Attach the pump and the 3-way valve (Part #29), leaving enough output tubing for easy access to the valve.
5) Attach output tubing to 3-way valve pointing down into the tub.
6) Attach input tubing from Step 4 to the last end of the 3-way valve.
7) Attach the output tubing from Step 4 to the output nozzle and submerge in tub.
8) Follow manufacturer's instructions for the water pump. Maintain water level above the pump's input valve.

*Note: One pump generates enough pressure for more than one box. We use an 8-way garden water splitter (https://www.ebay.com/itm/252874426549?ul_noapp=true) for multiple boxes, with a 2-way valve for each box to regulate water flow. These splitters have an even numbers of nozzles, and unused nozzles must be sealed off to maintain water pressure. Options for splitting water flow with with odd numbers of boxes include: 1) Adding and closing tubing and a 2-way valve (Part #30), 2) Sealing nozzles with acrylic adhesive, 3) Melting nozzles shut, or 4) 3D Printing a custom part.

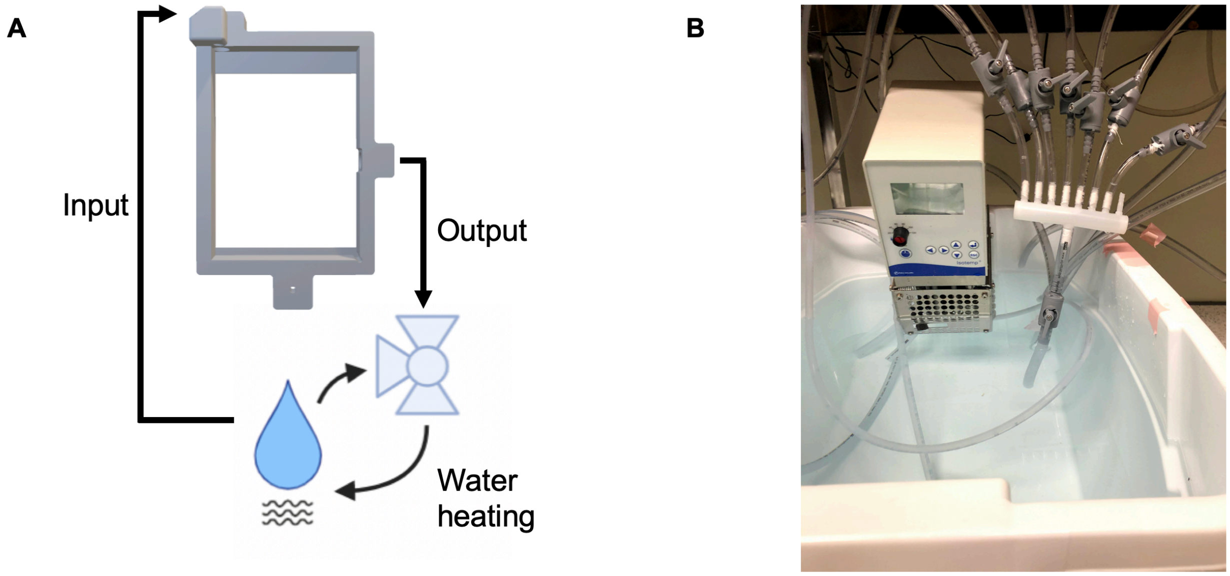

***Supplementary Figure 6. Heat Shock Assembly. (A)*** *Diagram of water flow. A single box contains one input and one output nozzle connected to the fish chamber. Water is heated and circulated using a 3-way valve inside a tub. Once the water reaches optimal temperature, the valve is manually switched to direct water into the input tubing. Water flows underneath the plate containing the fish and out through the output port back into the tub.* ***(B)*** *Our setup includes up to 8 outlets into 8 different inputs, each for a different box. To maintain optimal flow for each box, a 2-way valve (Part #30) is placed on each input tube line to decrease or increase water flow depending on the pressure.*

#### Optomotor Assay Setup

The optomotor assay is an optional component of the box. Hardware includes a projector (Part #31), a corner bracket (Part #13), socket head screw (Part #32), washer (Part #33), and wires to connect to the computer and power (Parts #34-38). Substitute manufacturer-provided HDMI and power cords with those listed in the BillofMaterials.

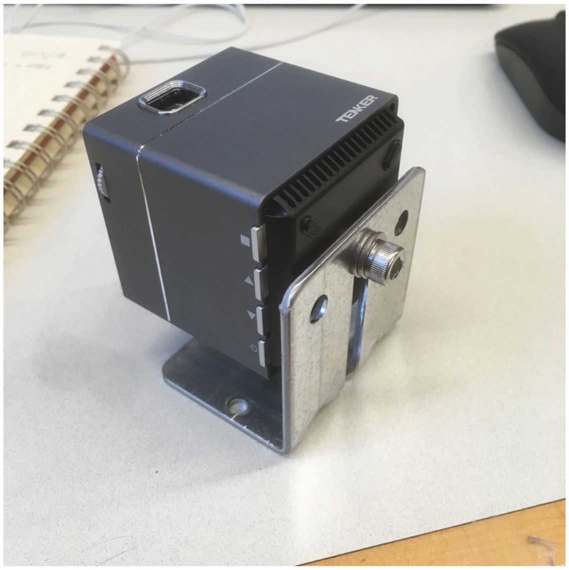

***Supplementary Figure 7 Optomotor Projector Assembly.*** *Plug in the USB-to-micro-USB and mini HDMI-to-HDMI cords before attaching the corner bracket. Align the bracket and tighten the socket head screw to secure the projector. Use the HDMI to Display port adaptor to connect to the Display Port cord to the HDMI end of the mini HDMI to HDMI cord. This setup stabilizes the projector and stands on top of the white light to project onto the fish chamber shelf. Place a sheet of paper or projection material between the fish chamber and the shelf to make movies visible to fish.*

Grating movies (vdark20_vlight250_For.avi, vdark20_vlight250_Rev.avi) are included in the supplementary files folder along with a script to create your own with varying bar width or contrast (makeOMRgrating.py and dependency png.py). Run python makeOMRgrating.py -h for more info. The script creates PNG frames that must be stitched to generate an AVI movie. We recommend using ImageJ (https://imagej.nih.gov/ij/download.html)

Once ImageJ is installed,

1) Open the application.
2) Load files by clicking File->Import->Image Sequence
3) Click on one of the images and click Open.
4) Confirm "Number of Images" matches the number of frames and check the Sort names numerically box. Click OK.
5) Click File->Save As->AVI… Confirm frame rate matches input. Refer to comments in makeOMRgrating.py for more info.

### Data Acquisition Setup/Software

#### Computer Requirements

| | |
|---|---|
| Processor: | Intel Core i7 4.50 GHz |
| Internet connection: | Ethernet (LAN) |
| Memory (RAM): | Recommended: 16 GB |
| Hard Drive: | 1TB SSD |
| Operating System: | Windows 10 |

One computer per box is required. To prevent interruptions during experiments, disable Windows Update and automatic restarts/shutdowns/sleeping modes.

#### PCB Board Construction/Teensy Installation

Light and tapper control relies on a custom printed circuit board (PCB). The gerber files (SleepBox_gerbers.zip) can be sent to a PCB manufacturer for fabrication. Assemble the board with the electronics and attachments specified from the BillofMaterials (Parts #39-52) and according to the instructional videos (Electronics Videos.zip). PCB design files (DesignFiles.zip, subject to MIT license) enable design modification.

*Assembling Electronics Enclosure*

Completed electronics boards are enclosed to prevent potential damage (**Supplementary Figure 8**). The casing includes two 3D printed ends attached with acrylic boards for ease access. The surface transducer and the white light must be connected to the board before adding the enclosure.

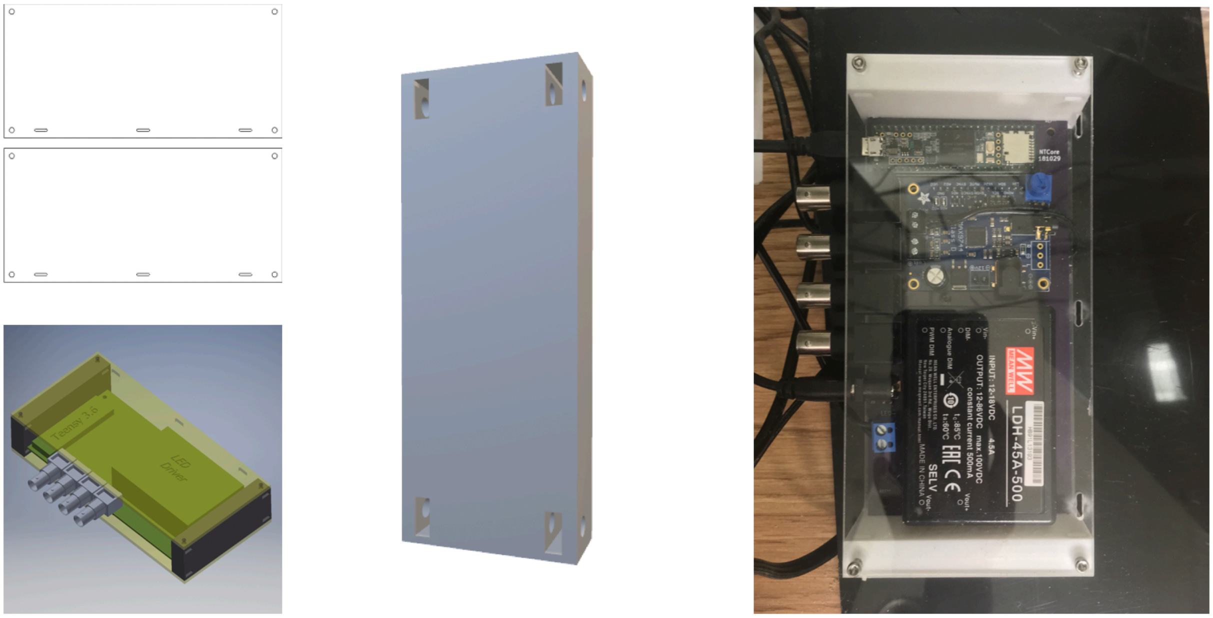

***Supplementary Figure 8. Electronics Board Enclosure Assembly.*** *The PCB and electronic components must be assembled before enclosure. Assembly instructions are in the Electronics Videos. If the PCB does not fit into the 3D printed side slot (Simple Board Side.stl), sand slot edges until the PCB board fits snugly. Use the ½" long socket head screw and corresponding hex nuts (Parts #50 and #51) to attach the acrylic enclosure panels (Electronics Enclosure Top Bottom.dxf) to the 3D printed side pieces. Insert hex nuts into the slots in the side pieces, and screw the board into the hex nut. Enclose the electronics board in a plastic bag to prevent water damage in the event of a leak from the output tube.*

##### *Optimizing the Electronics Setup*

The electronics are controlled through Teensy microcontroller and Arduino software. Download the latest version of Arduino (https://www.arduino.cc/en/Main/Software) and Teensyduino (https://www.pjrc.com/teensy/td_download.html).

1) Arduino must be installed first before Teensyduino (add-on for Arduino software)
2) Download the supplementary folder teensy36_dma_fish_box and open the Arduino file **teensy36_dma_fish_box**
3) Upload the file to the Teensy microcontroller. In Tools->Port, select the COMport corresponding to the teensy board and click Upload.
4) You will be prompted to press the button on the Teensy to continue uploading the file. The button should be blinking until pressed. Click Verify to confirm script was uploaded.
5) If the upload was successful, open the serial monitor and type **a1 p;**
   You should hear a loud tap from the surface transducer.

*Note: You can control the surface transducer and the white light manually using this script, by opening the serial monitor and typing commands. See Command Table for Teensy Control.

*Light Calibration*

Light panels must be calibrated to deliver consistent light across boxes. The required photodiode is listed in the BillofMaterials.

1) Connect the photodiode BNC to trigger 3 on the PCB (the BNC second from the teensy USB), place the photodiode directly on the light panel, and turn on the photodiode.
   *Note: the BNC connector must be secured properly by pushing the cable in until it clicks.
2) Open the Arduino serial monitor and connect to the teensy.
3) Send 'c' to start the calibration. Program will output lines of the following:

   *duty cycle, rise time, fall time, measured intensity*

   The first few lines should have intensity of less than a few units (ideally 0), while rise and fall times should be close to 500.
4) After calibration, send 'T' to test the calibration. It should start outputting lines of the following:

   *target intensity, actual intensity, accuracy (less than or equal to 1)*

   Make note of the minimum target intensity and the worst accuracy.
5) If the calibration is satisfactory, save by sending 'S'. The calibration will automatically load when the Teensy is powered on and can be inspected by sending 'R'.

*Command Table for Teensy Control.*

| **Teensy Command** | **Function** |
|---|---|
| *Sound* | |
| f<frequency in hertz> | Set the frequency of the sound in hertz |
| a<amplitude> | Set the amplitude of the sound where amplitude range from 0 (no sound) to 1 (max) |
| d<duration> | Set the duration of the sound in milliseconds |
| D<delay duration> | Wait command in milliseconds |
| P | Play configured sound |
| w<waveform index> | Select waveform type:<br>0: sine wave<br>1: square wave |
| g<amplifier gain> | Set the digital gain of the audio amplifier from 0 (no sound) to 63 (max). The default is 63 |
| *Light* | |
| v<duty cycle> | Set the uncalibrated brightness of the light from 0 (light off) to 600 (max) |
| c | Start light calibration routine |
| T | Test the current light calibration |
| S | Save the current calibration to non-volatile memory |
| L | Load the calibration saved in non-volatile memory |
| R | Report the current calibration. Any line with a -1 denotes an invalid calibration point where the light output was either 0 or the rise or fall time failed to pass the threshold (10 ms max) |
| b<brightness> | Set the calibrated output of the light panel from 0 (light off) to a maximum determined by the calibration. |

The Arduino script for the Teensy microcontroller contains several commands. An example command for an auditory pre-pulse inhibition sequence would be: **a0.01 f1400**

**d5 p D300 a1 f1400 d5 p;** with the semi-colon signifying the end of the command string. These commands are written in an events file with specific delivery times. See *Data Acquisition Control* for more information.

#### Camera Setup

*Installation*

A Grasshopper3 USB3 (FLIR) high-speed camera is used to record animals. Standard FLIR drivers work with their free FlyCapture2 camera control software. Since our 1-second movies are captured at exactly 285 fps, we use a specific driver iteration and software version to achieve this. Because more recent versions cannot achieve our settings, we include ours in Supplementary Software. The default analysis pipeline assumes that high-speed movies are captured at 285 fps, and must be modified as necessary for other capture speeds (classifications of movements may change based on framerate, etc).

We added a C-Mount 50mm fixed focal lens (Computar) at a working distance of 0.5 m. An IR filter (Lee) is inserted, since white light is used to maintain day/night cycles and deliver stimuli. The camera must be focused manually, while zoom can be adjusted manually or automatically in the FlyCapture2 software.

*Software*

Download and install FlyCapture2 (Included in Supplementary Software; FlyCapture_2.10.3.145_x64.exe).

*Updating the Camera Driver*

The most recent driver iteration does not allow for proper video acquisition, so a previous driver iteration must be installed. After installing FlyCapture2, upload the driver file gs3-u3-2.20.3-00.ez2 using the Driver Control GUI located in FlyCapture2 applications.

***Warning:** Do not disconnect the camera at any point during driver installation. An interruption may corrupt camera software.

1) Open Driver Control GUI
2) Confirm camera is connected and its name appears at the Device box
3) Open the gs3-u3-2.20.3-00.ez2 file
4) Click Upload

Camera settings can be adjusted using FlyCapture2. Some settings are obligatory for data acquisition, while others must be optimized based on desired assays and box environment (**Supplementary Figure 9**).

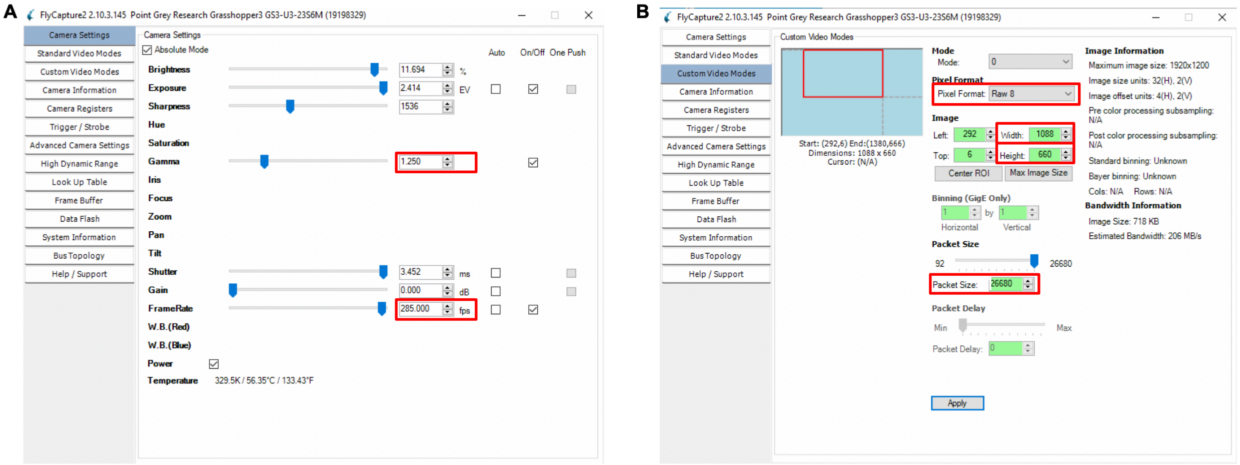

***Supplementary Figure 9. FlyCapture2 Settings. (A)*** *Red boxes indicate required values for the video acquisition. Some of these values can be adjusted based on lighting, but framerate must be set to 285 fps. To adjust values, uncheck boxes under Auto* ***(B)*** *Red boxes indicate required camera view orientation values. The Top and Left values are used to center the camera view, and will thus not match values shown here. Pixel format sometimes reverts to Mono 8, and must be returned to Raw 8 before each experiment to acquire the expected fps.*

#### Data Acquisition Control

Data acquisition is controlled by custom National Instruments (NI) LabVIEW software. Install the recent version of LabVIEW per manufacturer's instructions, together with the following packages need to be installed:

- LabVIEW Vision Development Module
- NI-IMAQdx

All scripts are included in Supplemental Software. LabVIEW expertise is not required to run the scripts, but will aid in adding new options to the software. To use the camera in either FlyCapture2 or LabVIEW, switch the driver for the corresponding software using Device Manager in Windows:

1) Open Device Manager (**Supplementary Figure 10**) using the search box located in the bottom left hand corner of the screen. You can also pin the Device Manager icon to the start menu for ease access.

2) Scroll through the Devices and locate PointGrey or FLIR.
3) Right click on PointGrey/FLIR and select “Update Driver”.
4) Click “Browse My Computer for Driver Software”.
5) Click “Let Me Pick from a List of Available Drivers” on My Computer
6) IMAQdx is the driver for LabVIEW. Click Next to update the driver.
7) The driver for LabVIEW does not always update normally and may produce an error message saying the update was unsuccessful. Restart your computer to complete the update.

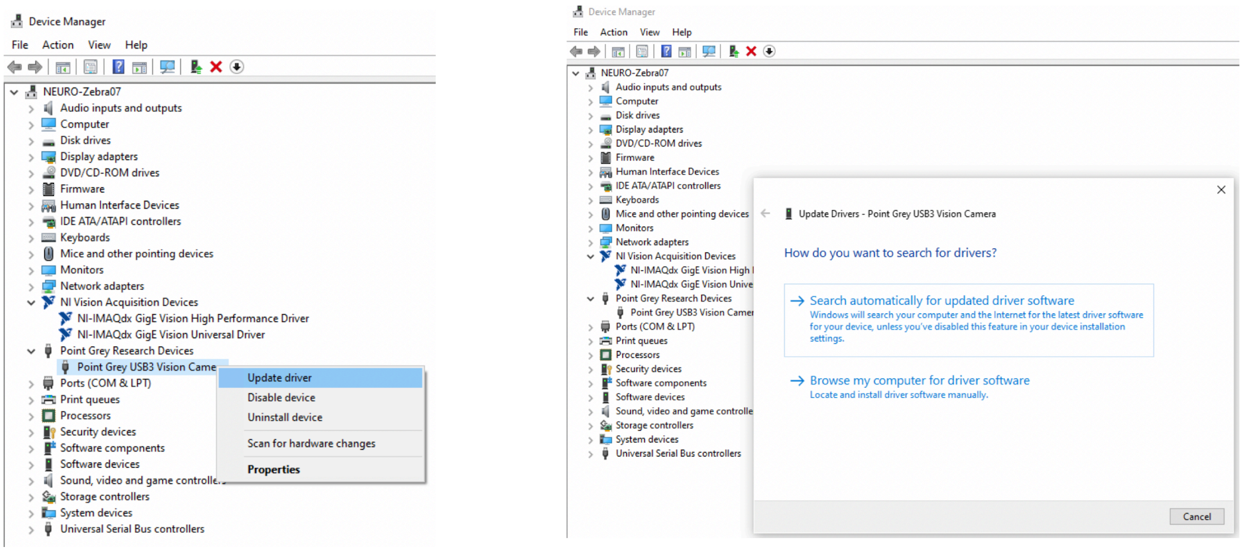

***Supplementary Figure 10 Switching the Camera Driver***. *In Windows, use the Device Manager to switch between LabVIEW and camera drivers. Initially, the camera will appear as PointGrey USB3 Vision Camera, and can be switched to LabVIEW IMAQdx driver.*

#### Starting a Behavior Run

##### *Setting ROIs*

Establish a consistent naming system for your input/output files, especially if using multiple setups. One approach is to use identical names for every run, and rename when transferring data from the computer to the analysis cluster. Path names in LabVIEW can be set to default after first use by clicking Edit->”Make current values default”.

1) Create a folder for your behavior run.
2) Open FlyCapture2. Confirm that settings are correct and that the fish plate is in focus and fully in view. Click the save button to save a PNG image of the plate in your new folder.

3) Open the LabVIEW file **Generate ROIs.vi**. This script uses the PNG image from Step 2 to generate ROI files for downstream analysis. Set the paths for the image and output files (**Supplementary Figure 11A)**. While we typically overwrite the ROI file from a previous run, users may need to generate an empty text file to accept output the first time the script is run.
4) Click the white arrow to start the program.
5) The cursor will turn into a crosshairs pointer. Click on the top left edge of the picture, then click the “Top Left” button. Repeat for the top right edge and bottom right edge.
6) Script will generate ROI files upon clicking the bottom right edge button. Green ROI boxes should appeared on your image in the script (**Supplementary Figure 11B**). Confirm that ROIs match individual wells, and repeat Steps 4-6 if necessary.
7) The **Generate ROIs.vi** script works with multiple plate formats. Users can adjust the number of rows/columns as desired.

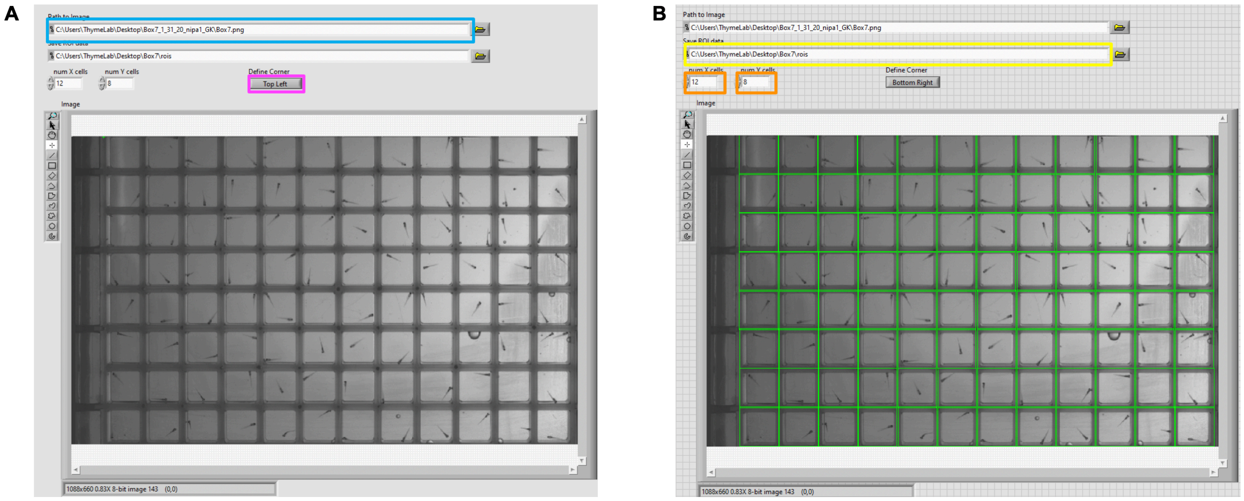

***Supplementary Figure 11 Using the Generate ROIs Script****. **(A)** Running the Generate ROIs script displays a PNG image designated in the input path (blue box) and changes the cursor to a crosshair. Clicking on the image then designates the point listed in the Define Corner field (magenta box), while clicking on the Define Corner field switches to the next point. Users sequentially designate the Top Left, Top Right, and Bottom Right corners of the plate. **(B)** After setting the bottom right position and clicking on the Define Corner button, the script finishes and outputs ROI files to the specified path and name (yellow box). The green boxes represent the ROIs for each well. If alignment is poor, re-run the script until optimal alignment alignment is achieved. The zoom tool (magnifying glass on left toolbar) may assist in accurate point designation. For different plate formats, users can change the number of ROI in the x and y directions (orange boxes).*

*Creating an Events File*
The main LabVIEW script reads an events file and relays commands at designated times. These files can be made in Notepad/WordPad or automatically generated with a Python script. An example line would be:

9:30:00 AM 1 a0.01 f1400 d5 p;

Each line contains:
- a time point
- an event identifier
    - 0 to not collect a movie,
    - 1 to collect a 1-sec 285 fps movie,
    - 2 to collect a 30-min 30 fps movie, or
    - 3 to load a movie
- A command string for the Teensy or the name of the movie to load.

Included in the supplementary files is an example **eventsfile** containing stimulus blocks. Time, event IDs, and command strings must be separated using tabs instead of spaces. However, elements within the Arduino command string are separated with spaces. Failure to adhere to this tabs/spaces format will cause the program to fail or interrupt an experiment. The eventsfile must also use correct time format (no 0s in front of single digit hours).

*Start Data Acquisition*
Turn off white light before starting. The first line of the eventsfile should turn the light on, and succesful execution of this command indicates that the Data Acquisition.vi script is working. We typically set the time soon after the start of the run and confirm lights-on before closing the box:

9:30:00 AM 0 b200;

1) Use Device Manager to Update the Camera Driver to IMAQdx. You may need to restart the computer if there is an error (**Supplementary Figure 10**).
2) Open **Data Acquisition.vi**, which controls the Teensy microcontroller, the camera, and generates movies and testlog files for the experiment (**Supplementary Figure 12A, Supplementary Figure 12B).**

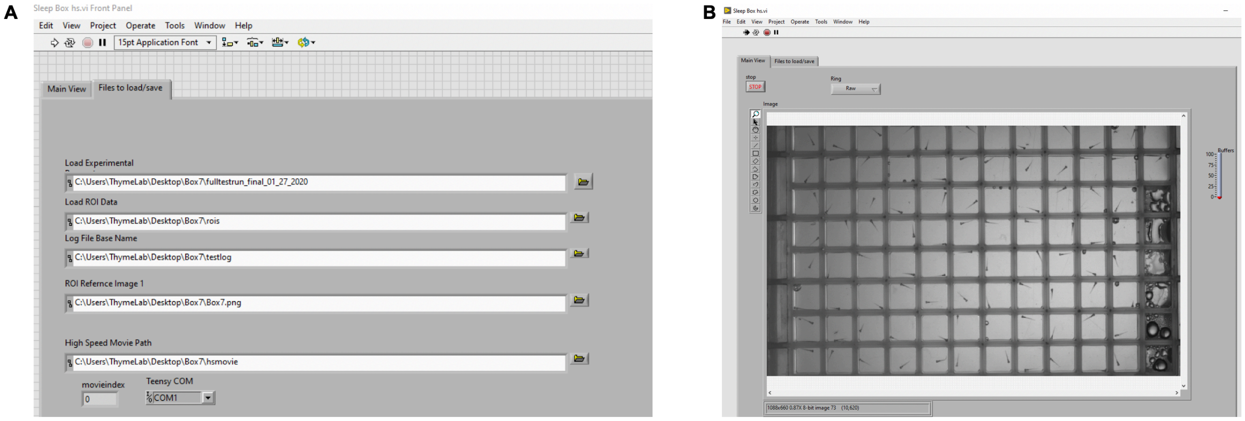

***Supplementary Figure 12. Using the Data Acquisition Script.*** ***(A)*** *The main program (and its dependencies: VLCPlayer.vi, Summarize Motion hs.vi, Scheduler hs.vi, ROI Reference hs.vi, Map to Well (ROI) Simple hs.vi, and Detect Motion hs.vi) for the behavior run requires three input files and two output file paths (and one folder for movies if desired). The Teensy COM must be switched to the correct port. The program uses the experimental file as its time course and the ROIs file with a reference image to determine where animals are within the camera feed. The program will continue until no additional commands are given by the experimental file. *Warning: include a final command for long after you plan to stop acquistion, as the system will terminate soon after the last command.* ***(B)*** *The main view shows the camera feed. Observation is critical, as this view shows recorded data. Absence of a feed indicates errors in data collection.*

3) Set the paths for each file to the folder you created:
    a. Load Events: events file containing the timepoint commands of the behavior run
    b. Load ROI Data: binary file created from the **Generate ROIs.vi** program (**Supplementary Figure 11**).
    c. Log File Base Name: name given to the three logs generated from the experiment: timestamps, motion data, and centroid data, as well as any slow-speed movies (do not include spaces or punctuation).
    d. ROI Reference Image: PNG image taken in FlyCapture2.
    e. High Speed Movie Path: path name for the high-speed movies collected during the run (do not include spaces or punctuation).
    f. Load Movie Folder: The path to a folder containing desired movies to use in an experiment, such as moving gratings for optomotor response. *Note: This path does not need to be filled for the script to run..
4) Change the Arduino settings to match microcontroller comport.
5) Click start run.
6) Once the run is done, close LabVIEW and manually switch off light. See Optimizing Electronics Setup for more info on manually controlling the microcontroller.

### Data Analysis Software

#### Introduction

The analysis software was developed in the Python programming language. This guide assumes basic familiarity with Linux command-line interfaces. It is highly recommended to analyze the data in a high-performance computing cluster. Many clusters use the SLURM workload manager, and we have included example SLURM job submission scripts.
***Warning**: As an important general rule, do not use spaces or punctuation in file naming. Only a few specific characters, such as an underscore, are compatible with Linux and these analyses.

To use analysis software on different systems, we include a Python environment file (**behavior_environment.yml**).

1) Computing clusters typically include many software packages. If so, load the most recent version of Anaconda for Python3 (i.e module load Anaconda3/<release#>). If not, install Anaconda for Python3. We used version 5.3.1, and Python2-based Anaconda will generate errors.
2) After loading or installing Anaconda, enter **conda env create -f behavior_environment.yml.** This will begin the installation process.
3) After installation, activate the environment by entering **source activate behavior_env**, or deactivate by entering **conda deactivate behavior_env.**
4) Load Anaconda and activate your environment for any future analyses.

#### High-Speed Movie Tracking

The first step of the analysis determines the centroid of each animal. For a detailed explanation of the approach, refer to comments within the Python scripts.

1) Activate the **behavior_env** environment.
2) Within the experiment data folder, run **highspeedmovieanalysis.py** with arguments for the ROI string text file, a high speed movie, and the events file (i.e. python highspeedmovieanalysis.py -r "rois_string" -m "hsmovieFri, Nov 22, 2019_<movie#>.avi" -e “eventsfile.txt”). An example SLURM submission script (**track.slurm**) is provided in Supplemental Software. Arguments for the script include:

```
-h, --help  # optional argument to remind user of available flags
-r # string version of the rois file
-m # movie file
-e # events file
-p # pixel threshold (default=3)
```

-f # movie fps (default=285)

***Warning:** Depending on the computing cluster configuration and the number of simultaneous jobs, analysis will fail or produce empty data files for individual movies. Run **queuecheck_behavior.py** to isolate these “dropouts,” and resubmit them for analysis. Specifically, **queuecheck_behavior.py** identifies movies for which the analysis produced no files or empty data files named *.motion2 or *.centroid2. In the unlikely event that the analysis fails during file generation, **queuecheck_behavior.py** will not detect dropouts, and errors will appear during downstream operations. These types of failures can be diagnosed by confirming that output *.motion2 and *.centroid2 files are similarly sized between movies (run “ls -l”). Repeating **highspeedmovieanalysis.py** for failed movies should correct any issues.

#### Processing High-Speed and Slow-Speed Data

The next step is to create a matrix of the genotypes/conditions for each animal in the plate using **splitgenotypes.py**. Example matrices (**gene_matrix)** are included in the supplementary files (**gene_matrix** for single mutants/condition; **duplicatedgene_matrix** for paralog double mutants). **splitgenotypes.py** includes logic identifiers for all combinations of fish genotypes as well as drug administration (see step 4 below). Users can also designate invalid wells, such as empty wells or wells with bubbles that preclude data acquisition.

1) Create genotype matrix text file: identifyingname_matrix. We generate these matrices by copying a Google spreadsheet of genotypes into a text document with the Vi editor. Example from Google spreadsheet:

| | 1 | 2 | 3 | 4 | 5 | 6 | 7 | 8 | 9 | 10 | 11 | 12 |
|---|---|---|---|---|---|---|---|---|---|---|---|---|
| A | b | B | hom | hom | hom | het | het | hom | hom | hom | het | B |
| B | het | het | hom | het | hom | het | hom | het | hom | het | hom | B |
| C | hom | het | het | het | hom | het | het | hom | het | het | het | B |
| D | het | het | hom | hom | het | het | het | hom | hom | hom | hom | het |
| E | hom | hom | hom | het | hom | hom | Het | het | hom | het | hom | het |
| F | hom | het | hom | hom | hom | het | Het | hom | het | het | hom | hom |
| G | hom | hom | e | hom | hom | het | hom | het | hom | het | hom | het |
| H | hom | het | hom | 2 | het | hom | hom | hom | het | hom | het | het |

***Warning:** Names must not include any spaces, tabs, or punctuation, and must exactly match the identifiers summarized in step 4. White space (tabs) separates wells.

2) Run **splitgenotypes.py**
   This script parses the matrix file, counts the number of each well for each identifier, and creates a file called **genotyping**.
   ***Warning:** Genotypes will be scrambled if the camera is mounted backwards. Comments within the script detail how to modify if needed.
3) Open **genotyping**.
   To identify groups for statistical analysis, type and asterisk (*) at the beginning of each identifier line. If more than two groups are selected with an asterisk at this stage, the next script will generate all permutations of the groups comparisons. Two samples, one control group and one test group, are required for the statistical analysis.
4) Run **makeslurmfiles.py**
   This script parses **genotyping** and generates necessary slurm files and output directories for analysis. Confirm that generated files adhere to nomenclature **submission_script_gene_het_vs_gene_hom.slurm**. **makeslurmfiles.py** uses specific genotype identifiers to automatically determine control and test groups. Faulty nomenclature (including different capitalization) will bypass genotype identifiers and necessitate manual test and control group input. New identifiers can be added to the script as needed. A "logic dictionary" at the top of the script lists current group identifiers: dmso, drug, wt, wtwt, hetwt, wtandhet, hetandwt, hethet, het, homwt, wthom, hethom, homhet, hom, homhom. The script should be modified as needed for a particular computing cluster and for changes to file names. However, a number of options are available if needed and do not require modifying the script:

   -statsfile # prefix for output statistics file (default = "linearmodel")
   -gfile # genotyping file (default = "genotyping")
   -efile # events file (default = "fulltestrun_final_01_27_2020")
   -prefix # prefix for slow-speed data (default = "../testlog")
   -hprefix # prefix for high-speed data (default = "../hsmovie")
   -pfile # plot parameters file (default = "../PlotParameters")
   -esfile # sections file (default = "../sectionsfile")
   -scriptpath # path to the folder containing the Python analysis scripts in the ProcessMotion folder; there is a default, but this value MUST be changed for your system
   -other # can add any other flag applicable to the processionmotiondata.py script, such as the -graphonly flag that skips production of data files and goes directly into graph generation.

5) Modify/create a **sectionsfile**

This file groups phases of the experiment for statistical analysis. Confirm that this file follows the exact timeline of movie/event generation, as it uses the LabVIEW timestamp file. We include an example sectionsfile and corresponding **eventsfile** that illustrates potential event groupings. For instance, a long habituation block can be divided into sections to represent the habitution process vs post-habituation testing.
***Warning:** sectionsfile errors are a common cause for analysis script failures. Confirm that the start/end times of each section fall within the overall experiment timestamps by cross-referencing with the LabVIEW testlog.timestamp file. Downstream checks will generate warnings in case of discrepancies. The sectionsfile should not contain any extra spaces, blank lines, or punctutation, and should use 24h time nomenclature without additional 0s in front of single-digit hours.

6) Run **processmotiondata.py** by submitting the slurm scripts automatically generated by **makeinputgenotyping.py**, or by replicating the **script_gene_het_vs_gene_hom.slurm** format.
*Note: **makeslurmfiles.py** from Step 4 will have generated a **jobsubmission.sh** shell script for automatic submission of multiple slurm files. Simply type ./jobsubmission.sh to submit all of its listed slurm files. If only one comparison is needed, users can instead enter “sbatch submission_script_gene_het_vs_gene_hom.slurm”.
7) **processmotiondata.py** analyzes the high-speed and slow-speed data and uses three supporting scripts (**fileloading.py**, **setupgraphsandsavedata.py**, and **graphsstatsandfilter.py**) to load data and create processed data files and graphs. Three additional files contain code for classes and organize data (**Fish.py**, **EventSection.py**, and **ProcessedData.py**). The list below summarizes available arguments for **processmotiondata.py** (Comments in **fileloading.py** provide additional descriptions):

```
 -h, --help # optional argument to remind user of available flags
 -graphonly # skips production of data files and goes straight to graph or re-graphing
(default=False)
 -j # PlotParameters file; required flag, no default
 -t # testlog.timestamp file; required flag, no default
 -e # eventsfile; required flag, no default
 -c # testlog.centroid file; required flag, no default
 -d # testlog.motion file; required flag, no default
 -m # prefix for high-speed movies; required flag, no default
 -g # genotype ids from a inputgenotypeids_scripted file generated by setup script;
required flag, no default
 -s # sectionsfile; required flag, no default
```

-n # number of wells in the plate (default=96)
-i # milliseconds per highs-speed movie frame (default=3.508772)
-v # threshold values for slow-speed data, distance and dpix (default="0.5,3.0")
-w # threshold values for high-speed data, distance and dpix (default="0.9,3.0")
-f # threshold of frame number for slow-speed, distance and dpix (default="1,3")
-x # threshold of frame number for high-speed, distance and dpix (default="2,3")
-a # bins for the activity/sleep data (default="1/60,60/600,60/3600,1/3600")
-y # thresholds, distance and dpix, for activity/sleep data (default="1,10")
-b # bins for slow-speed bout data (default="60,600,3600")
-z # filters for identifying potential seizures (default="4.0,300,1300,70")
-l # baseline light level, used to identify what is a dark flash to filter o-bends (default=200)
-o # obendfilters, currently identified based on response time and sum of absolute value of heading angle (default="60,10", ie O-bends = responses with time >60 and sum absolute heading angle > 10)
-p # cbendfilters, currently identified based on response velocity and response cumulative dpix (default="0.2,1500", ie C-bends = responses with response velocity > 0.2 and cumulative dpix > 1500)

*Note: C-bend and O-bend filtering likely depend on the high-speed framerate and other baseline conditions in the setup. We visually inspected movies and scatter plots of corresponding metrics to isolate values representing different types of responses.

8) Once analysis is complete, use **sort_pvalues.py** to define graphs with significantly different values between comparison groups. Arguments for this script include:
-statsfile # name of input file (default = "linearmodel*out", ie all *.out files starting with linearmodel in the working directory)
-ofile # name of outfile (default is automatic generation based on input file name)
-include # name fragment of graphs to include for analysis of p-values (For example, include only darkflash data, etc) (default="ribgraph*", representing all analyses)
-exclude # name fragment of graphs to exclude
-cutoff # p-value cutoff (default=0.05)
-overwriteanova # substitute the ANOVA p-value for the linear mixed model p-value calculated for slow-speed time sections (default=False)
-showfulldata # show all the Ns, SSMDs (default=False)
-sortalpha # sort p-values by graph names instead of by the p-values (default=False)

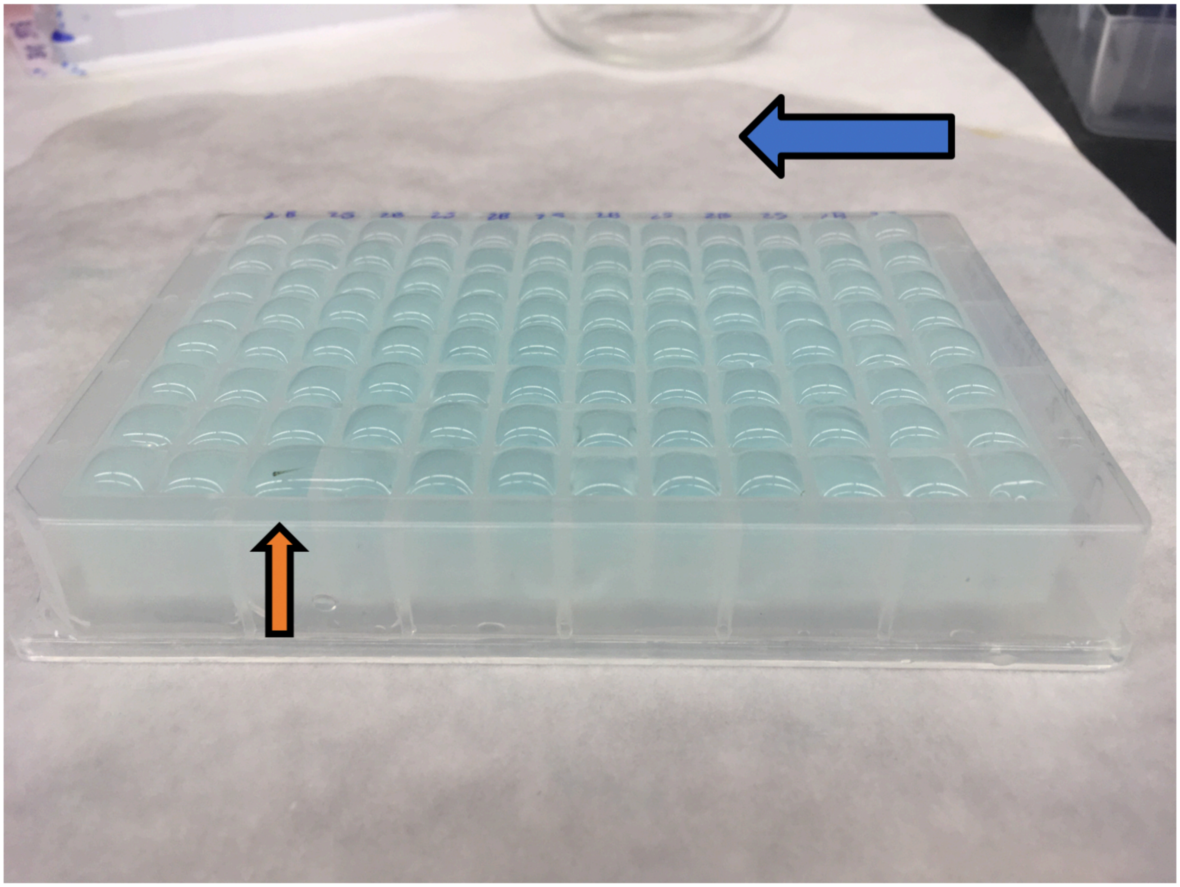

***Supplementary Figure 13. Fish Plate Loading.*** *Water will evaporate if plate remains in box for extended times. Accordingly, we cover plates with a semi-permeable seal (Thermo Fischer Catalog #4311971). Larvae are loaded into a 96-well plate (Agilent Part #201242-100) and placed over ice to slow their movements. Next, each well is overfilled as shown above. The seal is then pressed down onto the plate at one edge and slowly flattened across the plate (Blue arrow) along with excess water. This method minimizes bubble formation inside wells, which can obscure data acquisition. Warning: Larvae may commence movement or float to the top of the well before sealing and can move to other wells (Orange arrow). If your fish regularly do not sink to the bottom after icing (essential for sealing the film), check the components of your blue water or other embryo medium.*

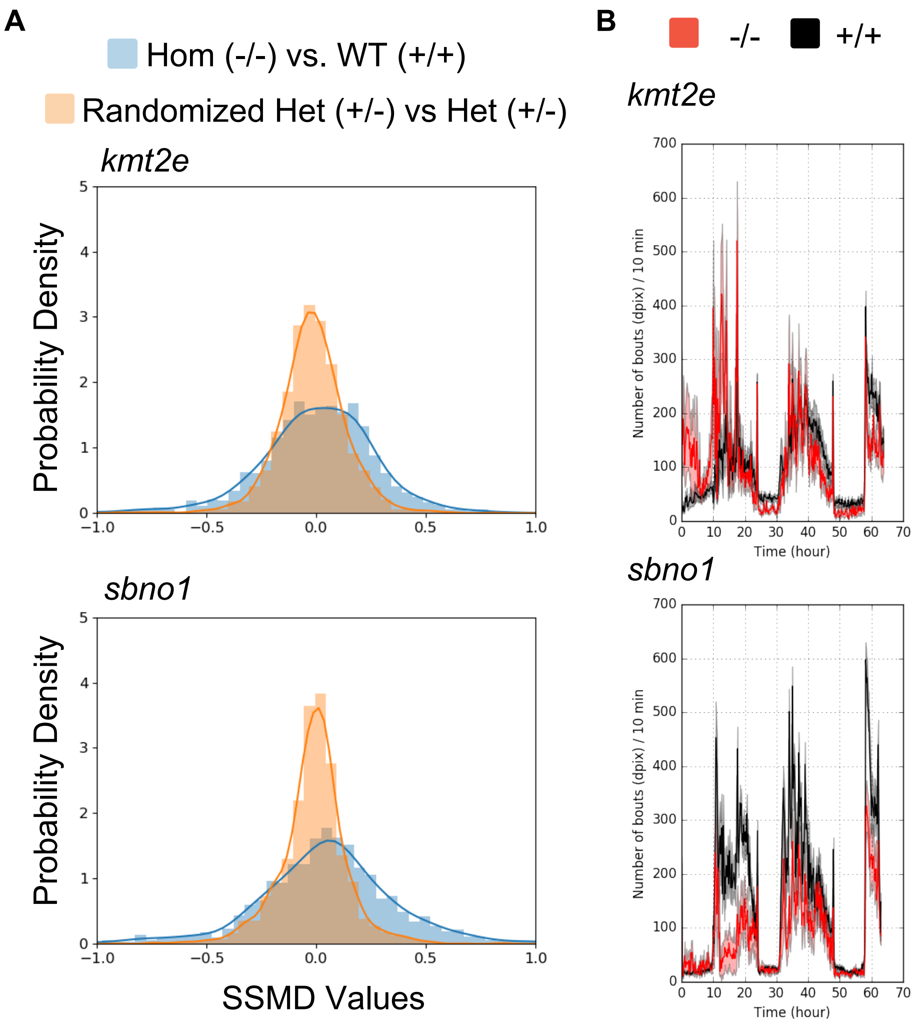

***Supplementary Figure 14. Mutant SSMD Comparison. (A)*** *Histograms of SSMD values for kmt2e and sbno1 mutants. kmt2e: N -/- = 7, +/+ = 29, +/- = 24 for both Het groups. sbno1: N -/- = 8, +/+ = 24, +/- = 27 for both Het groups.* ***(B)*** *Example graphs showing strong frequency of movement phenotypes for both mutants. These two mutants are among the top five strongest phenotypes observed in a previously published screen of 165 mutants (Thyme, et al., 2019). More subtle and specific phenotypes would almost certainly be obfuscated by genetic background differences if animals from different clutches were compared (see main text and Figure 8).*
